# Kingdom-wide analysis of the evolution of the plant type III polyketide synthase superfamily

**DOI:** 10.1101/2020.04.28.059733

**Authors:** Thomas Naake, Hiroshi A. Maeda, Sebastian Proost, Takayuki Tohge, Alisdair R. Fernie

## Abstract

The emergence of type III polyketide synthases (PKSs) was a pre-requisite for the conquest of land by the green lineage. To study the deep evolutionary history of this key family, we used phylogenomic synteny network and phylogenetic analyses of whole-genome data from 126 species spanning the green lineage. This study thereby combined study of genomic location and context with changes in gene sequences. We found that two major clades, CHS and LAP5/6 homologs, evolved early by a segmental duplication event prior to the divergence of Bryophytes and Tracheophytes. We propose that the macroevolution of the type III PKS superfamily is governed by whole-genome duplications and triplications. Intriguingly, the combined phylogenetic and synteny analyses in this study shed new insights into changes in the genomic location and context that are retained for a longer time scale with more recent functional divergence captured by gene sequence alterations.

## Introduction

During plant evolution, the number of specialized metabolites and the enzymes responsible for their synthesis exploded (1, 2). The number of protein folds however, remained restricted (1, 3). This is likely because novel biosynthetic pathways generally originate by gene duplication events and/or by functional divergence of existing genes (2). Commonly, duplicated genes, from already enzymatically active enzymes, were subjected to differential mutations resulting in a broader substrate specificity and a lower activation energy of catalysis, led single enzymes catalyzing multiple reactions and thereby synthesizing multiple products (1).

450-500 million years ago, Charophycean freshwater green algae began to colonize land (4-6). Early land plants needed to adapt quickly to their altered environment leading to the innovation of novel metabolic pathways, including phenylpropanoids, sporopollenin and lignin biosynthesis (1, 7). They achieved this by ‘recycling’ enzymes from existing core pathways, co-opting them and evolving novel functionalities (2, 8-12). The availability of plant and algae genomes is enabling us to trace the diversification of plant enzyme families undergoing evolutionary alterations and shaping the vast plant chemical diversity seen today (13-16). These studies whilst highly informative were, however, restricted to phylogenetic analyses of gene/protein sequences and did not take into account the genomic context for all analyzed species which would reveal the deep ancestral history and the points of diversification of a gene family.

Given their strategic importance within the phenylpropanoid pathway we postulate that PKSs may have played a major role in the colonization of land, by providing the precursors for the synthesis of flavonoids (17) and sporopollenin (18, 19). The type III PKS superfamily is a prime example of how the recruitment of an existing pathway led to the diversification of metabolic routes (12, 20). PKS enzymes are likely derived from the β-ketoacyl acyl carrier protein (ACP) synthases of fatty acid biosynthesis (21), as they share a protein fold (12, 21). Indeed PKSs, including type III PKS, like their predecessors from fatty acid metabolism, catalyze the sequential head-to-tail condensation of two-carbon acetate units derived from a malonate thioester into a growing linear polyketide chain (12). All type III PKSs share a common αβαβα structural fold, a conserved catalytic triad consisting of Cys-His-Asn and act as homodimers consisting of ∼40 kDa monomeric subunits.

Type III PKSs are widely distributed in bacteria (22, 23), fungi (22, 24) and ubiquitously present in plants (12). More than 20 functionally different plant type III PKSs have been described (Fig. 1) which share 30-95% sequence identity (20). Among the most prominent members are chalcone synthases (CHSs) (20). CHSs catalyze the entry point of flavonoid metabolism and are well characterized in a number of model species (25-27). The diverse functions of the 20 types of enzymes arose due to their differences in (i) substrate specificities, (ii) numbers of condensations and (iii) cyclization reactions all of which are ultimately governed by the sequence of the genes which encode them (12, 20).

**Fig. 1:**
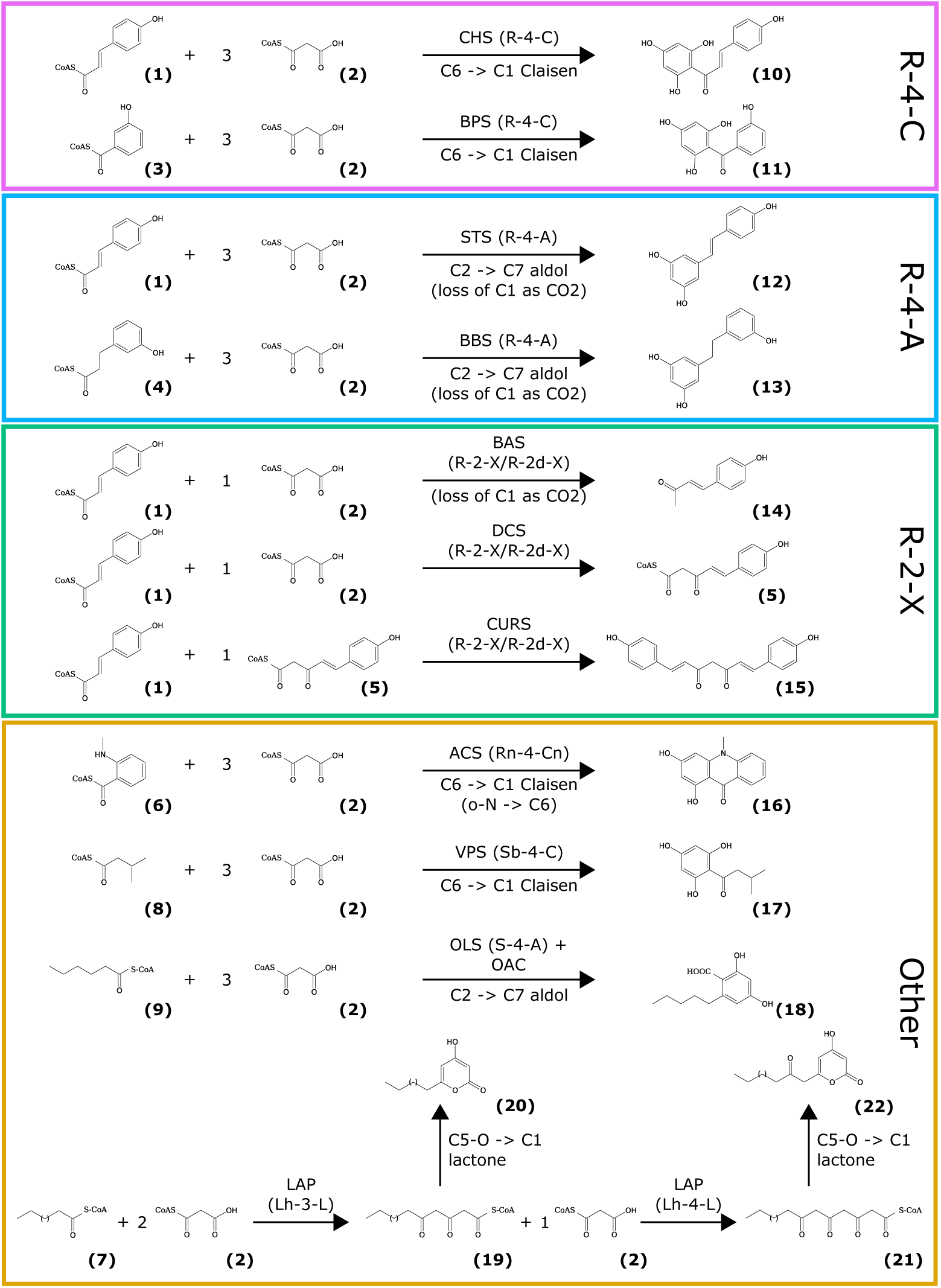
Overview on important reactions catalyzed by type III PKS. The reaction type is defined on the basis of combinations of three features according to pPAP (28, 29): 1) starter substrate: three categories for the starter substrates based on their acyl group: ring (R), short chain (S, C2 to C12) or long chain (L, up to C26). Additional characters are added to specify acylgroups (branched chain, b; carboxylate group, c; hydroxy group, h; nitrogen, n); 2) number of condensations: indicates the number of methylenecarbonyl units in the intermediates. Additional characters are added to specify other substrates than malonyl-CoA (methylmalonyl-CoA, m; ethylmalonyl-CoA, e; acetoacetyl-CoA, a; diketide-CoA, d); 3) mechanism of intramolecular cyclization: Claisen, C; aldol, A; lactone, L; no cyclization, X; nitrogen-carbon, n. The features 1) and 2) are interrelated since Claisen- or aldol-type cyclizations require typically at least four and lactonization at least three carbonyl units in the intermediate. The pPAP software will define four major categories for a classified reaction type (R-4-C for CHS and BPS; R-4-A for STS and BBS; R-2-X for BAS, DCS and CURS; Other for ACS, VPS, OLS and LAP). (1): *p*-coumaroyl CoA, (2): malonyl CoA, (3): 3- hydroxybenzoyl CoA, (4): dihydro-*m*-coumaroyl CoA, (5): p-coumaroyl diketide CoA, (6): *N*- methylanthraniloyl CoA, (7): fatty acid acyl CoA, (8): isovaleroyl CoA, (9): hexanoyl CoA, (10): naringenin chalcone, (11): 2,3’,4,6-tetrahydroxybenzophenone, (12): resveratrol, (13): 3,3’,5-trihydroxybibenzyl, (14): benzalacetone, (15): bisdemethoxycurcumin, (16): 1,3- dihydroxy-*N*-methylacridone, (17): phloroisovalerophenone, (18): olivetolic acid, (19): triketide intermediate, (20): triketide pyrones, (21): tetraketide intermediate, (22): tetraketide pyrones. CHS: chalcone synthase, BPS: benzophenone synthase, STS: stilbene synthase, BBS: bibenzyl synthase, BAS: benzalacetone synthase, DCS: diketide-CoA synthase, CURS: curcumin synthase, ACS: acridone synthase, VPS: phloroisovalerophenone synthase, OLS: 3,5,7-trioxododecanoyl-CoA synthase/olivetol synthase, LAP: acyl-CoA synthase/hydroxyalkylpyrone synthase/less adhesive pollen.

In order to study the evolution of type III PKS we here utilized a phylogenomic network approach (30-32), to study the syntenic relationships between genomic regions of a myriad of species spanning the green lineage. Synteny, the conservation of gene content and order within or between genomes, infers a shared evolutionary history. Synteny analysis, therefore, provides a means to examine the ancient history of gene evolution, since gene sequences can change their functionality by mutations, while synteny can be retained over a longer time scale. Such approaches allow the inference of the orthology, timing and mode of duplication of pairs/ groups of genes (33). Here we modified the approach in order to allow cross-Kingdom analysis and combined it with the phylogenetic approach to investigate the relationship between functional divergence of various genes and their genomic location. These combined analyses revealed an early segmental duplication event that led to the emergence of the LAP and CHS clades. We also provide evidence that the evolution of the type III PKS superfamily is governed by genome region duplication and triplication events following the emergence of the LAP and CHS clades. We propose an evolutionary route for the CHSs governed by a whole-genome triplication event and its subsequent diversification in a Fabales-specific clade. Our combined results are further discussed in the context of early land plant colonization and the maintenance of presence of type III PKS genes in their genomic context.

## Results

### PKS copy numbers widely vary among different plant and green algae species

Flavonoid and sporopollenin biosynthesis evolved on the terrestrialization of the green lineage. To study the diversification of the PKS superfamily, the fully-sequenced genomes of 126 species spanning the green lineage were queried for the number of PKS copies they possessed. To provide a robust classification of protein families, we used OrthoFinder (34) and MCL (35). We detected 63,344 and 61,643 groups containing more than one protein sequence for OrthoFinder and MCL clustering, respectively. Within this dataset, the type III PKS superfamily formed one protein group with 1,621 different protein sequences detected by either of the clustering methods, of which 1,551 protein signatures/sequences were jointly detected by both. Within these groups, all previously characterized and described PKS sequences were recovered (see Supplementary File 1). Reassuringly, the protein family of β-ketoacyl ACP synthases, which exhibits sequence similarity to PKS, formed a separate group in both the OrthoFinder and MCL output. *PKS*s are present in all land plants albeit in varying copy numbers. However, *PKS* were not found, or only found in low copy numbers, in the Chlorophyta, and are absent in Chlorokybophyceae, Mesostigmaphyceae and Coleochaetaphyceae of the Charophyta (see Supplementary Text). By contrast, type III PKS were detected in *Penium margaritaceum* (36).

### Synteny network analysis detects clade-specific and reaction type-specific clusters

To study the diversification of the type III PKS superfamily we followed a synteny network approach (32). Whole genomes of 126 species were compared in a pairwise manner, followed by robust block detection of regions containing type III *PKS* genes and network analysis to detect syntenic clusters within the network. The resultant network contained 706 vertices corresponding to syntenic regions containing single or multiple type III PKS genes of which 166 vertices corresponded to regions with tandem-duplicated genes from a total of 105 species (Supplementary Table S2). Tandem-duplicated genes may play important roles in providing genetic redundancy, gene dosage balance, genetic robustness, and to provide an additional means for divergence in transcriptional regulation and protein sequence (37-40). The highest number of tandem-duplicated PKS genes in one syntenic region was 23 (*Arachis duranensis,* containing mainly ‘R-4-A’-type PKS sequences, Fig. 1 and Fig. 2). *Arachis ipaensis* (21 genes) and *Vitis vinifera* (20 genes) had the second- and third-highest numbers of tandem-duplicated genes (also containing mostly ‘R-4-A’-type sequences), respectively.

**Fig. 2:**
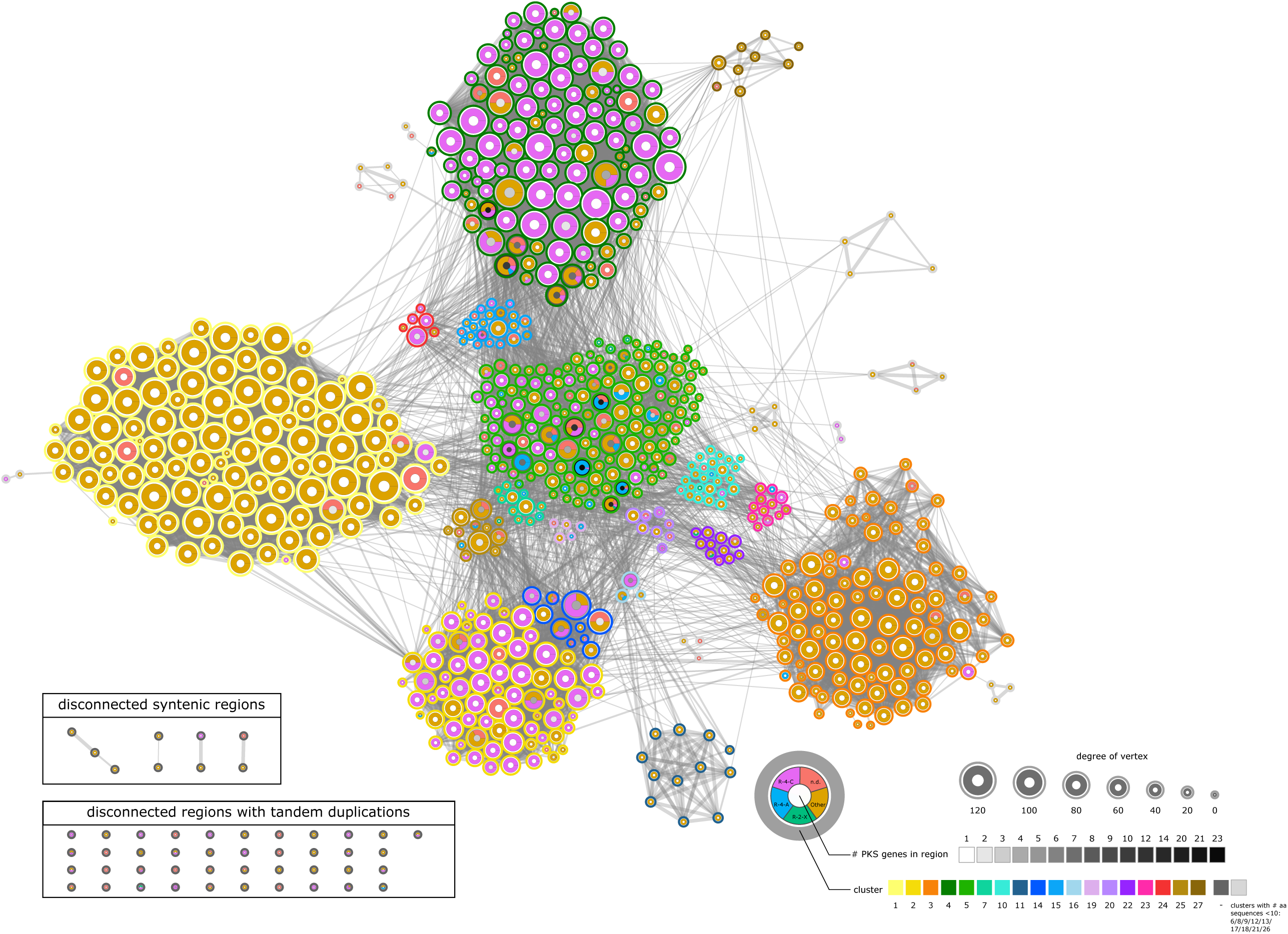
Network showing synteny between regions containing type III PKS genes. The network contains four ‘R-4-C’-enriched syntenic clusters corresponding to CHS function (2, 4, 5 and 14) and four syntenic clusters with LAP5 and LAP6 orthologs (1, 3, 11 and 27). The syntenic clusters showed species-specific distribution (Supplementary Figure 5). Each vertex depicts information on the pPAP classification of PKS genes within the genomic region, the number of PKS genes within the region (number of tandem duplicates) and the cluster membership. Clusters were detected by applying the community detection algorithms, “fastgreedy”, “walktrap”, “leading eigenvector” and “multilevel” on the weighted network, followed by calculation of the sum of distance between the different methods (0 if cluster is identical, 1 if cluster is different) and application of affinity propagation clustering to define the number of clusters and their membership. Vertex size reflects degree of the vertex (unweighted number of edges connecting to the vertex).

For most of the possible syntenic relationships, the network was characterized by an absence of synteny between PKS-containing regions. The network was sparse with 2.4% of all possible edges (5,851 of 248,865 possible edges). We applied a robust cluster detection approach on the network using four different algorithms that resulted in the detection of 27 syntenic clusters (Fig. 2).

The previously characterized dicot *CHS* genes, such as *Solanum lycopersicum Solyc09g091510/SlCHS1* and *Solyc05g053550/SlCHS2* were located to the syntenic cluster 2, while *Arabidopsis thaliana AT5G13930/TT4* and *Zea mays Zm00001d052673/C2* were found in syntenic cluster 4 (Fig. 2). Of the 27 clusters, some were clade-specific. For instance, cluster 4 with the second highest number of *PKS* did not contain syntenic regions in the majority of Asterids species. This suggests that either Asterids never had *PKS* locating to cluster 4 or that specific synteny was lost or the regions/genes deleted (see Supplementary Table S3).

Syntenic regions specific to the Commelinids were found in clusters 9, 21, 27 and partly 11 and 22. BLAST analysis of PKS sequences revealed that the annotation of bisdemethoxycurcumin synthases was exclusive for Commelinid species. Genes in cluster 9 were mainly annotated as ‘Other’ by pPAP and bisdemethoxycurcumin synthase (BCURS) by BLAST analysis (BCURS are canonically classified as ‘R-2-X’ by pPAP (29)). Genes in cluster 22 were mainly annotated as ‘R-2-X’ or ‘Other’ via pPAP and as BCURS by BLAST (type ‘Other’ by pPAP and acridone synthase by BLAST in the *Citrus* genus). These two clusters indicate a Commelinid-specific invention of BCURS and PKS of the ‘Other’ type. Cluster 21 contains genes of unknown function annotated as ‘Other’ by pPAP and CHS-like by BLAST (gray color in Fig. 2).

PKS in the clusters 3 and 11 were almost exclusively classified as ‘Other’ by pPAP and annotated as type III polyketide synthase A or CHS-like based on BLAST analysis. PKS in the clusters 1 and 27 were almost exclusively annotated as ‘Other’ by pPAP and type III polyketide synthase B or CHS-like by BLAST (Fig. 2). The syntenic cluster 3 showed depletion of syntenic regions of members of the Commelinids with the exception of *Elaeis guineensis* and *Phoenix dactylera*. This cluster was specific for members of the Commelinids. By contrast, syntenic cluster 1 showed depletion of syntenic regions of members of the Commelinids with the exception of *Musa acuminata, Sorghum bicolor, Oropetium thomaeum, Hordeum vulgare* and *Leersia perrieri*, while the syntenic cluster 27 exclusively contains Commelinid PKS. Clusters 1 and 27 contain a characterized *LAP5*, while 3 and 11 contain characterized *LAP6* genes (18, 19) (e.g. *AT4G34850/LAP5* in cluster 1 and *AT1G02050/LAP6* in cluster 3) and homologs (Supplementary Table S4).

### Synteny network analysis detects four enriched clusters of ‘R-4-C’-type PKS corresponding to chalcone synthase

Further analyses of the pPAP-classified PKS showed that specific kinds of *PKS* genes show certain distributions among different clusters. In particular, we found that ‘R-4-C’-type *PKS* corresponding to the CHS function are overrepresented in the syntenic clusters 2 (and tightly associated 14), 4, and 5 (Fisher’s Exact test, odds ratio: 13.33, p-value < 2.2e-16) and the ‘R-4-C’-classified PKS were only found to a minor extent in some other syntenic clusters (Supplementary Figure 5). This observation led to the hypotheses that the major CHS clusters 2/14, 4 and 5 either evolved independently three times, *or*, a scenario that is more likely, evolved by duplication of the gene regions following the loss of the synteny between these three CHS-enriched clusters.

### Phylogenetic analysis indicates timing of the appearance of ‘R-4-C’-enriched clusters

To further evaluate these two hypotheses, we performed a phylogenetic analysis using PKS amino acid sequences to link their syntenic cluster membership with sequence divergence reflected in the phylogenetic tree. The analysis incorporated sequences from over 180 species thereby dramatically extending previous phylogenetic analyses on the type III PKS (41). Here, we included 1,607 different amino acid sequences and obtained detailed phylogenetic relationships using the Maximum likelihood method (Fig. 3) having high transfer bootstrap expectation values (42) for all major clades (Supplementary Figure 4). The phylogenetic tree showed that ‘R-4-C’/CHS are mainly present in one very large clade (Fig. 3) that is dominated by sequences of the type ‘R-4-A’, ‘R-4-C’ and ‘Other’. This clade also contains the sequences of AT5G13930/TT4 from *Arabidopsis thaliana*, Solyc09g091510/*Sl*CHS1 and Solyc05g053550/*Sl*CHS2 from *Solanum lycopersicum* and Zm00001d052673/C2 from *Zea mays*. By contrast, the other clade contains mainly sequences of the type ‘R-2-X’ and ‘Other’.

**Fig. 3:**
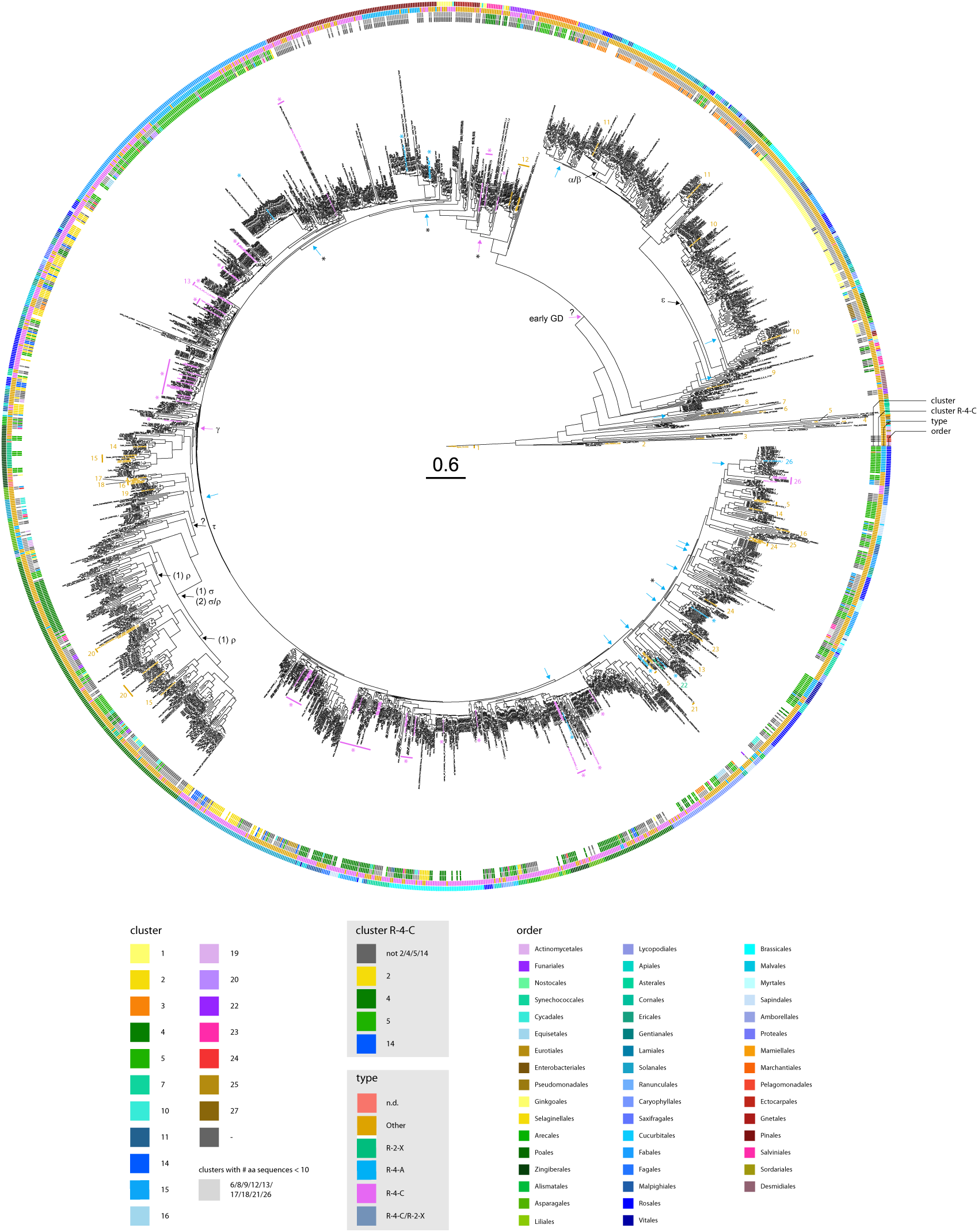
Phylogenetic gene tree of type III PKS amino acid sequences with additional information on syntenic cluster membership, type of the sequence according to pPAP classification and the taxonomic order of the species to which the sequences refers to. The phylogenetic tree indicates that the LAP ortholog containing clade and the ‘R-4-C’-containing clade evolved by an early duplication event (early GD). The ‘R-4-C’-containing syntenic clusters 2/14, 4 and 5 containing clades form highly related but mostly distinct clades in the phylogenetic tree indicating that cluster 2/14, 4 and 5 evolved by duplication events. The LAP5/6 clade contains orthologs of LAP5 and 6 from *Arabidopsis thaliana* (Supplementary Table S4). Blue arrows indicate ‘R-4-A’ sequences that evolved independently several times. Blue arrows with star (*) indicate STS sequences of *Vitis vinifera, Arachis duranensis, Arachis ipaensis* and from Gymnosperms. Magenta arrows indicate duplication events involving (proto) ‘R-4-C’-type PKS sequences. Magenta arrows with a star indicate the origin of ‘R-4-C’ sequences from these events. Proteome files of 126 species were queried for type III PKS sequences via a HMM profile of known PKS sequences. For species, for which proteome files were not available, PKS genes were selected based on previous annotation. Amino acid sequences were aligned via hmmalign and sites with more than 20% missing values were removed. The gene tree containing 1607 unique sequences were build using RAxML using 1000 bootstrap replications. Supplementary Figure 3 shows extended information on the sequences. The tree shows high transfer bootstrap expectation values (42) for all major clades (cf. Supplementary Figure 4). Experimentally validated sequences are font-colored in purple for chalcone synthases (CHS, indicated by purple star), blue for stilbene synthases (STS, indicated by blue star), green for benzalacetone synthase and orange for other PKS sequences. 1: triketide and tetraketide pyrone synthase, PKS18; 2: phloroglucinol synthase; 3: RppA; 4: quinolone synthase; 5: β-ketoacyl carrier protein synthase III; 6: 2’-oxoalkylresorcylic acid synthase, ORAS; 7: CsyB; 8: 2′-oxoalkylresorcinol synthase, ORS; 9: hydroxyalkyl α-pyrone synthase, LAP; 10: hydroxyalkyl α-pyrone synthase, LAP5; 11: hydroxyalkyl α-pyrone synthase, LAP6; 12: stilbenecarboxylate synthase, SCS; 13: valerophenone synthase, VPS, VPS annotated with ‘R-4-C’ also show prenylflavonoid synthase function; 14: diketide-CoA synthase, DCS; 15: curcuminoid synthase, CS/CURS; 16: octaketide synthase, OS; 17: chromone synthase; 18: aleosone synthase; 19: pyrrolidine ketide synthase; 20: alkylresorcylic acid synthase, ARS; 21: acridone synthase, ACS; 22: benzalacetone synthase; 23: olivetol synthase, OLS: 24: 2- pyrone synthase, 2-PS; 25: orcinol synthase; 26: benzophenone synthase, BPS.

Sequences of the type ‘R-4-C’/CHS corresponding to the syntenic clusters 2, 4, 5 and 14 located in neighboring, yet mostly distinct subclades. Comparing the ‘R- 4-C’-dominated clade (Fig. 3) and the distribution of sequences present in the syntenic clusters (Supplementary Figure 5), it seems likely that the ‘R-4-C’-type genes in the syntenic clusters originated from the same evolutionary event by duplication/triplication events. From this result, we hypothesize that ‘R-4-C’-type *PKS* were initially contained in a protocluster that contained primordial ‘R-4-C’-type *PKS*. This was most likely followed by one or two early duplication events, depending if the LAP or CHS clade evolved first, creating the syntenic protoclusters 2/5/14 and 3, followed by loss of synteny between the protoclusters. Phylogenetic analysis further suggests that type III *PKS* from the type ‘R-4-C’ found in syntenic cluster 5, mainly from the Fabales, originated from type III *PKS* sequences from syntenic cluster 2, since the ‘R-4-C’-type sequences are located closely in the phylogenetic tree (Fig. 3).

Considering the situation that we found for syntenic cluster 5, it seems most probable that the primordial *CHS* genes were initially present in syntenic protocluster 2/14, followed by a loss of synteny between syntenic protocluster 2/14 and 5 (Supplementary Figure 5) leading to the present situation of having four major CHS-enriched clusters. Interestingly, ‘R-4-C’-type PKS sequences from *Physcomitrella patens* were detected in the syntenic cluster 5 and 15 while PKS from *Azolla filiculoides, Marchantia polymorpha, Salvinia cucullata* and *Selaginella moellendorffii* were present in syntenic clusters 4 and/or 5. Within the phylogenetic tree, these sequences located both within the LAP5/6 and the ‘R-4-C’-containing clade. Furthermore, syntenic cluster 5 contains the duplicated regions of ‘R-4-A’-type PKS sequences of *Arachis duranensis, Arachis ipaensis* and *Vitis vinifera* corresponding to the STS function.

### Independent and multiple evolution of ‘R-4-A’-type PKS with stilbene synthase function

The clades containing the ‘R-4-A’-type STS of *Vitis vinifera* and of the *Arachis* genus are located within the ‘R-4-C’-type dominated clade (clusters 2/14, 4 and 5), which suggests that convergent evolution led to the emergence of type ‘R-4-A’-type PKS after the emergence of the ‘R-4-C’-type protocluster. The synteny network and the phylogenetic analysis suggest that syntenic cluster 5 is a relic from syntenic regions that were separated by the emergence of the ‘R-4-C’-type dominated clade since sequences of syntenic cluster disseminate along the phylogenetic tree (Fig. 3). In addition to these early evolutionary events, phylogenetic analysis further highlights several independent evolutionary events underlying the ‘R-4-A’-type III PKS. STS from *Vitis vinifera* and the *Arachis* genus evolved within the same syntenic cluster 5 independently *or*, although less likely given their evolutionary distances, evolved once and diverged into their sequences while maintaining STS function (Fig. 3). Other ‘R-4-A’-type PKS sequences evolved in several far-related taxa within the monocots, eudicots and Gymnosperms (Fig. 3). The Gymnosperm-specific ‘R-4-A’-type PKS form a monophyletic clade within the phylogenetic tree suggesting that the appearance of ‘R-4-A’ happened before their speciation. These ‘R-4-A’-type PKS correspond mostly to pinosylvin-forming stilbene synthases and evolved most probably from CHS or a protoform within the gymnosperms.

As in a previous study (43), the ‘R-4-A’-type bibenzyl synthases from the species of the Orchidaceae form a distinct clade from their ‘R-4-C’-type orthologs, suggesting their divergence into subfamilies before the speciation of this family. Next to the evolution of ‘R-4-A’-type sequences, we further recapitulated the proposed evolutionary trajectory of VPS from *Humulus lupus* and OLS from *Cannabis sativa* and proposed a possible evolutionary route of aleosone synthase, chromone synthase and OS in *Aloe arborescens* (see Supplementary Text).

### Combining syntenic network and phylogenetic analysis reveals timing of type III PKS evolution

Having performed independent phylogenetic and synteny analysis we next attempted to see what added value could be obtained by combined these analyses. Interestingly, the phylogenetic tree indicates a Commelinid-specific clade that mainly contains members of the syntenic clusters 4, 5, 10 and 22. It seems most probable that the PKS genes located in syntenic clusters 10 and 22 originate either from a duplication event in clade 5 and 4, resulting in the formation of 5 and proto 4/10/22, followed by a duplication of protocluster 4/10/22 and loss of synteny resulting in 10 and 22 (see (1) in Fig. 5) *or* by duplication of the clade 4 and 5 and diversification in the syntenic clusters 4, 5, 10 and 22 and loss of synteny between the syntenic clusters (see (2) in Fig. 5). Furthermore, the phylogenetic analysis indicates that the Angiosperm-specific clade containing LAP homologs (Supplementary Table S4) and members of the syntenic clusters 1, 3, 11 and 27 evolved from the protoclusters A and A* by a segmental duplication event (Fig. 3 and 5). This clade contains *At*LAP5 or *At*LAP6 orthologs within the Bryophyta, Marchantiophyta, Lycopodiophyta, Pinophyta, Ginkgophyta and Gnetophyta, instead of *At*CHS orthologs (Fig. 3). This finding suggests that the LAP5/6 orthologs in this clade originated before the divergence of these afore-mentioned taxa from Angiosperms. The clade containing the LAP5/6 orthologs in angiosperms either originated from one duplication event of the protocluster A leading to the formation of a syntenic cluster B, followed by a second duplication event leading to two regions of B that correspond to the proto 3/11 and proto 1/27 cluster (see (3) in Fig. 5) *or* by two duplication events of the regions A or A* leading to the proto 3/11 and proto 1/27 syntenic clusters followed by a loss of synteny between the two protoclusters (see (4) in Fig. 5). This clade in the phylogenetic tree indicates that one subclade, either 3/11 or 1/27, originated from the other by a duplication event before the divergence of the monocots and eudicots, followed by loss of synteny between the monocot-specific and eudicot-specific clusters. Within the subclade corresponding to the syntenic clusters 3 and 11, cluster 3 indicates a duplication event within the Brassicales, leading to the emergence of two copies of LAP6 orthologs within the Brassicales, which still share synteny with other members of the syntenic cluster 3 (Fig. 2), indicating a recent duplication event. However a caveat regarding the quality of the Gymnosperm genomes should be noted (for details see Supplementary Text).

### Drivers of genomic maintenance in syntenic regions

The results above show that the genomic regions containing type III *PKS* remain syntenic over a long evolutionary timeframe. This raises the questions why the synteny between PKS-containing genomic regions was maintained and if genes corresponding to specific biological processes contribute to their maintenance. We conducted a GO enrichment analysis on the genes that were reported as being syntenic and checked for enrichment against the background (all genes showing synteny for each species or all genes present in the data sets). We performed this analysis by using syntenic regions containing *PKS* genes (Supplementary Figures 6 and 7) and regions where we excluded the *PKS* genes prior to analysis (Fig. 4 and Supplementary Figure 8) to remove bias that originates from the presence of *PKS* genes. All comparisons showed similar results in their overrepresentations, with the exception of the comparisons where PKS genes were included and that showed specific enrichment of PKS-related terms. We also checked the overrepresentation for GO terms of the CHS-enriched syntenic clusters 2, 4, 5 and 14 by performing enrichment tests against the genes of syntenic regions containing PKS regions to test if the CHS-enriched syntenic clusters differ from the PKS-containing syntenic regions. The analysis did not show fundamentally different results from the other comparisons, rather the terms where specialized terms of enriched terms that were found in the other set comparisons (Supplementary Figure 9). This indicates that the CHS-enriched syntenic clusters share an evolutionary past with the PKS-containing background.

**Fig. 4:**
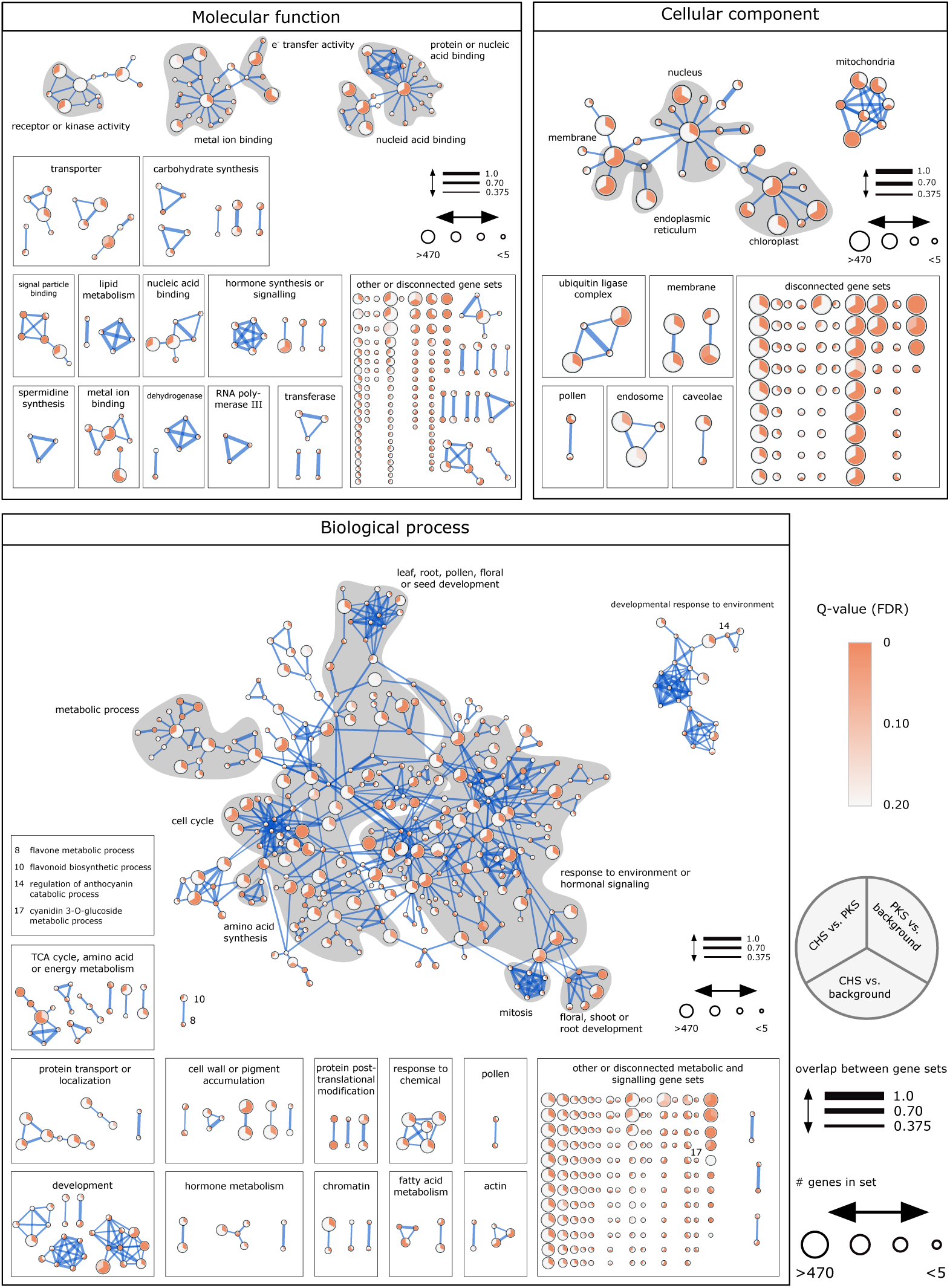
Gene Ontology enrichment of syntenic regions where *PKS* genes were excluded. *PKS* genes were removed prior to conducting enrichment analysis. Three enrichment sets were compared: A) Genes of *PKS*-containing syntenic regions were checked for enrichment against background (all genes in syntenic regions), B) Genes of syntenic regions in CHS-enriched clusters 2, 3, 5 and 14 against background, C) Genes of syntenic regions in CHS-enriched clusters 2, 3, 5 and 14 against syntenic genes of all species. The syntenic regions were enriched to a total of 509 (biological process), 230 (molecular function) and 108 (cellular compartment) significant terms (FDR-corrected q-value < 0.05). Enriched terms were categorized into higher categories. Many enriched terms in the category ‘Biological process’ can be linked to flavonoid-related processes (‘leaf, root, pollen, floral or seed development’, ‘response to environment or hormonal signaling’). The size of the vertex corresponds to the number of gene with the same GO term. Terms with a FDR-corrected q- value of <0.2 are displayed. Edges correspond to the similarity between terms based on their gene set overlap (50% Jaccard similarity and 50% overlap between terms with a cutoff of 0.375).

When excluding *PKS* genes, for the category biological process, the syntenic regions did show few enriched terms that are involved in flavonoid biosynthesis, for instance ‘cyanidin 3-O-glycoside metabolic process’, ‘flavonone metabolic process’ two flavonoid biosynthetic processes and isoflavonoid metabolic process (Supplementary Table S5 and S6). These terms were found partly in syntenic regions of Brassicales showing long syntenic regions or only in a smaller subset of species. The GO enrichment analysis revealed four (flavonoid) regulation-related processes and seven upstream biological processes (Supplementary Tables S5-8).

Unexpectedly, significant enrichment was detected for GO terms that are related to direct and indirect effects of flavonoids/polyketides on biological processes (genes in syntenic regions: 139 terms, all genes: 149 terms). These terms included processes that are linked to hormone metabolism, growth, development in response to hormonal signaling and responses to biotic and abiotic stress (Supplementary Tables S5 and S6).

## Discussion

During evolution, the biochemical repertoire of plants was expanded to synthesize a myriad of small molecules (1, 2). This chemical diversity was achieved by the recruitment of enzymes involved in primary metabolism and duplication of biosynthetic genes (1, 44-46). Here we presented a large-scale synteny analysis for the type III PKS superfamily in the green lineage and outlined the most likely evolutionary route for biochemical diversification of this important entry point of specialized metabolism. The products of the enzymatic activity of type III, flavonoids and building blocks for the pollen exine layer, are ubiquitously distributed in the plant kingdom and have been suggested to be pivotal prerequisites for colonizing the land. Indeed, the metabolic innovation of CHS rendered the shikimate pathway, a pathway only found in microorganisms, fungi and plants, considerably more important (47), such that nowadays it can carry 25% of the total C assimilatory flux (48) rendering it a high-flux bearing pathway, which is capable of producing a wide spectrum of polyphenolic compounds that play vital roles in development and biotic and abiotic interactions (49).

### Type III polyketide synthases evolved when colonizing terrestrial ecosystems

For non-land plants (non-embryophytes), that were included in the study here, we found PKS signatures only for some species of the Chlorophyta and *Ectocarpus siliculosus* (Ochrophyta). The PKS signatures from the Chlorophyta do not fall into the generic length range of PKS protein sequences suggesting that these PKS sequences are unlikely complete and functional proteins. However, experimental evidence in support of this supposition is currently lacking. The PKSs from *Ectocarpus siliculosus* fall within the range of generic PKS sequences found in vascular plants, however, all three sequences are located close to the outgroup of the phylogenetic tree. Previously, the *Ectocarpus siliculosus* genome was predicted to encode three type III *PKS* that may be involved in phloroglucinol biosynthesis (50, 51) that correspond to the sequences we found here. The shikimate pathway is also fully encoded in the genome of *Ectocarpus siliculosus*, however some downstream pathways, including the pathways leading to phenylpropanoids and salicylic acid, are lacking (50, 51). Genome mining of other brown algae species detected homologs of type III PKS (52-54). However, in the diatoms *Phaeodactylum tricornutum* and *Thalassiosira pseudonana* or in non-photosynthetic stramenopile oomycetes no type III PKS were found (51). Three putative type III PKS have been detected in *Pseudochattonella farcimen*, a unicellular alga belonging to the Dictyochophyceae within the Ochrophyta (55). These findings suggest that a lateral gene transfer of type III PKS, possibly by an ancestral actinobacterium, occurred following the separation of the diatoms from other members of the Ochrophyta, but before the brown algae diverged from the Pelagophytes and Dictyochophytes (51). Similarly, genes of other pathways, including the genes for mannitol, alginate and hemicellulose biosynthesis, are present in brown algae but absent in diatoms, suggesting a lateral gene transfer from an ancestral actinobacterium to *Ectocarpus siliculosus* (51, 56, 57).

It was previously stated that land plants originated from the Streptophytes of the Charophycean line (58) and that land plants and Zygnematophyceae form a sister relationship (59-62), rather than to the aquatic Chlorophycean group (47). For the two Charophyta species studied where full genomes were available, *Klebsormidium nitens* and *Chara braunii*, no type III PKS sequence were detected. Previously, members of the Charophyceae including the Charales, Coloeochaetales and Zygnematales, but not basal Charophyceae (Klebsormidiales and Chlorokybales) were shown to have cell walls similar to primary walls of embryophytes. Some of the analyzed Coleochaete have lignin or lignin-like containing cell walls, suggested to be derived from radical coupling of hydroxycinnamyl alcohols (63). The streptophyte *Klebsormidium flaccidum* harbored an ortholog of the *phenylalanine ammonia lyase* (*PAL*) gene (63). de Vries et al. (64) analyzed the genetic complement in streptophyte and chlorophyte algae with regard to the phenylpropanoid pathway and found cinnamyl/sinapyl alcohol dehydrogenase orthologs, coumarin, lignin and flavonoid biosynthetic gene orthologs in these taxa, and 4-coumarate:CoA ligase (4CL) among Streptophyta, but not Chlorophyta. These findings indicate that Streptophyta acquired phenylpropanoid activity, but no activity of type III PKS *or* that type III PKS were lost again in the course of evolution, the latter becoming more probable if the hypothesis of Stebbins and Hill that Charophyceae have secondarily returned to an aquatic habitat, after adaptation to terrestrial or amphibian life (65), is correct. Recently, the genomes of *Mesostigma viride(66)* and *Chlorokybus atmophyticus* (66) and *Penium margaritaceum* (36), a member of were published. *Mesostigma viride* and *Chlorokybus atmophyticus* did not show any type III PKS signatures within their genomes. By contrast, *Penium margaritaceum* was reported to contain eleven copies of type III PKS (36). This indicates that type III PKS appeared earliest with the Zygnematophyceae, but not with the earlier appearing Mesostigmatophyceae, Chlorokybophyceae, Klebsormidiophyceae and Charophyceae. To our knowledge, no full genome is available yet for a member of the Coleochaetophyceae, the next basal class to the Zygnematophyceae. Such a genome and additional genomes of the Zygnematophyceae represent (an) intriguing model(s) to study further the emergence of type III PKS in the green lineage and will yield higher resolution as to which step in evolution type III PKS emerged.

### Independent evolution of stilbene synthase function in different taxa

The evolution of type ‘R-4-A’-type *PKS* was studied previously. Schröder et al (67) found that STS and CHS sequences differ in charged/uncharged amino acids in homosites across the whole sequences. Tropf et al. (68) performed a phylogenetic analysis of CHS and STS sequences and found that STS grouped with CHS sequences and subsequently concluded, that STS evolved from CHS multiple times independently, while STS and CHS have a common evolutionary origin. Their mutagenesis studies showed that three amino acid changes in a CHS hybrid/chimera are sufficient to obtain STS activity. Austin et al. (69) revealed the structure of functional STS in *Pinus sylvestris* and identified the structural basis of STS sequences from CHS ancestors corresponding to a cryptic thioesterase activity in the active site, due to alternative hydrogen bonding network (‘aldol switch’). Guided by the STS protein structure, Austin et al. (69) performed mutagenesis studies to convert CHS into STS. Here, we found similarly that the type ‘R-4-A’ was associated to different clades in the phylogenetic trees supporting previous findings that STS evolved independently. The clade of gymnosperm STS seemed to appear after the divergence of gymnosperms with angiosperms since the ‘R-4-A’-type containing clade of gymnosperms locates close to the ‘R-4-C’- and ‘Other’-type containing clade of gymnosperms. We found that ‘R-4-A’-type sequences of *Vitis vinifera* and *Arachis sp.* are located in the same syntenic cluster. The fact that the STS sequences of *Vitis vinifera* and *Arachis sp.* do not form a monophyletic clade, however, suggests that independent evolutionary events generated the ‘R-4-A’ function in these species. In the *Arachis* genus, the ‘R-4-A’ function most likely evolved before the speciation of *Arachis duranensis* and *A. ipaensis* since both ‘R-4-A’-containing genomic regions locate to the same syntenic cluster and the ‘R-4-A’-type sequences to the same clade in the phylogenetic tree. Generally, we found that ‘R-4-A’ function is independent of the syntenic regions (Supplementary Figure 5 and 12). The phylogenetic analysis suggests, that also in gymnosperms the ‘R-4-A’ function evolved before speciation of the gymnosperm species under study.

### GO maintenance of synteny indicates no general formation of PKS-containing syntenic regions across the green lineage

The type III *PKS*-containing gene regions maintain their synteny over a long time and across a wide range of Taxa (Fig. 2 and Supplementary Figure 5). By contrast, some type III *PKS*-containing regions show no synteny to other regions, which could be attributed to low assembly quality (Supplementary Figure 13 C), small scaffold size (Supplementary Figure 13 E right plot) or to the high taxonomic distance from Angiosperms (Supplementary Figure 13 E).

Plant specialized metabolic pathways may be encoded as regulon-like gene clusters which consist of mostly non-homologous genes that are physically linked and functionally related via biosynthetic pathways and coregulated (70). Gene clusters for specialized metabolic pathways were previously shown for alkaloids (71), diterpenoid phytoalexins (72, 73), triterpenoids (74, 75), hydroxamic acids (76) and syringyl lignin (77). It was hypothesized that gene clustering provides a selective advantage due to more efficient inheritance since clustered genes are probably retained in the face of recombination. Furthermore, gene clusters allow for coordinate transcription via genomic and epigenetic mechanisms (1). In order to test whether this was the case for polyketide/flavonoid biosynthetic gene clusters, we conducted for the network containing syntenic *PKS*-containing regions a gene ontology enrichment test and co-expression analysis for *PKS* genes of the syntenic clusters 2, 4, 5 and 14 was carried out using the STRING database (Supplementary Text). The analyses showed no enriched terms for polyketide-related processes except some upstream related processes (related to acetate metabolism, shikimate metabolism) and no co-expression between *PKS*/*CHS* and other genes. In barley a gene cluster consisting of *Cer-c*, a CHS-like diketone synthase, *Cer-q*, a lipase/carboxyl transferase, and *Cer-u*, a cytochrome P450 hydroxylase was recently predicted making it, to our knowledge, the only published example to report a gene cluster containing a *PKS* gene (78). *Cer-c* in our study did not show synteny to other *PKS* genes. However, the results presented here render it unlikely that *PKS*s form gene clusters at least in the many species we studied.

Looking at other processes, we found enrichment of processes that showed biological involvement of polyketide products, especially in processes that are linked to hormonal processes and effects, besides enrichment of processes that did not show any overlap with other processes. It has been suggested, that flavonoid metabolism initially evolved as an internal physiological regulator/chemical messenger (79), rather than as a UV filter as proposed in Kubitzki et al. (47), since enzymatic capabilities and enzyme quantities should be low after recruitment from β-ketoacyl ACP synthases (79). Interestingly, many signaling related processes were enriched in the genomic context of PKS and CHS genes. It is possible that PKS-related genes governing such processes are an evolutionary remnant *or* provide a direct fitness effect, which maintains type III PKS genes in close genomic proximity to these genes. Alternatively or in addition, ‘hijacking’ of genomic regions that contain pivotal genes, such as genes that are linked to transcription and translation, amino acid metabolism, cell development and responses with the environment, could explain the presence of type III *PKS* in these genomic regions.

### Possible evolutionary route of the PKS superfamily, suggested by combined syntenic network and phylogenetic analyses

Type III PKS catalyze the sequential condensation of acetate units to a starter molecule. The reaction sequence mirrors the biosynthetic pathway of fatty acid synthases in primary metabolism (12). It has been hypothesized that type III PKS evolved from β-ketoacyl ACP synthases (12) due to their similar reaction mechanism and the presence of the αβαβα-fold. Using synteny information and information from a large-scale phylogenetic analyses we outlined an evolutionary route for the type III polyketide synthase superfamily (Fig. 5) after the emergence of the first version of a PKS protein. The evolution of type III PKS is primarily governed by an early gene region or genome duplication that formed the two major ‘R-4-C’-containing clusters (2, 4, 5 and 14), and an LAP ortholog-containing clade (Fig. 3). Whole genome duplications have neither been described for the liverwort *Marchantia polymorpha* (80) nor the Lycophyte *Selaginella moellendorffii* (4). However, they were described for *Physcomitrella patens*(81) and the Tracheophyta after the divergence from the Bryophyta (82, 83), indicating that the ‘R-4-C’-specific and LAP5/6 clades evolved possibly by a segmental duplication event before the divergence of the Bryophyta from the Tracheophyta.

**Fig. 5:**
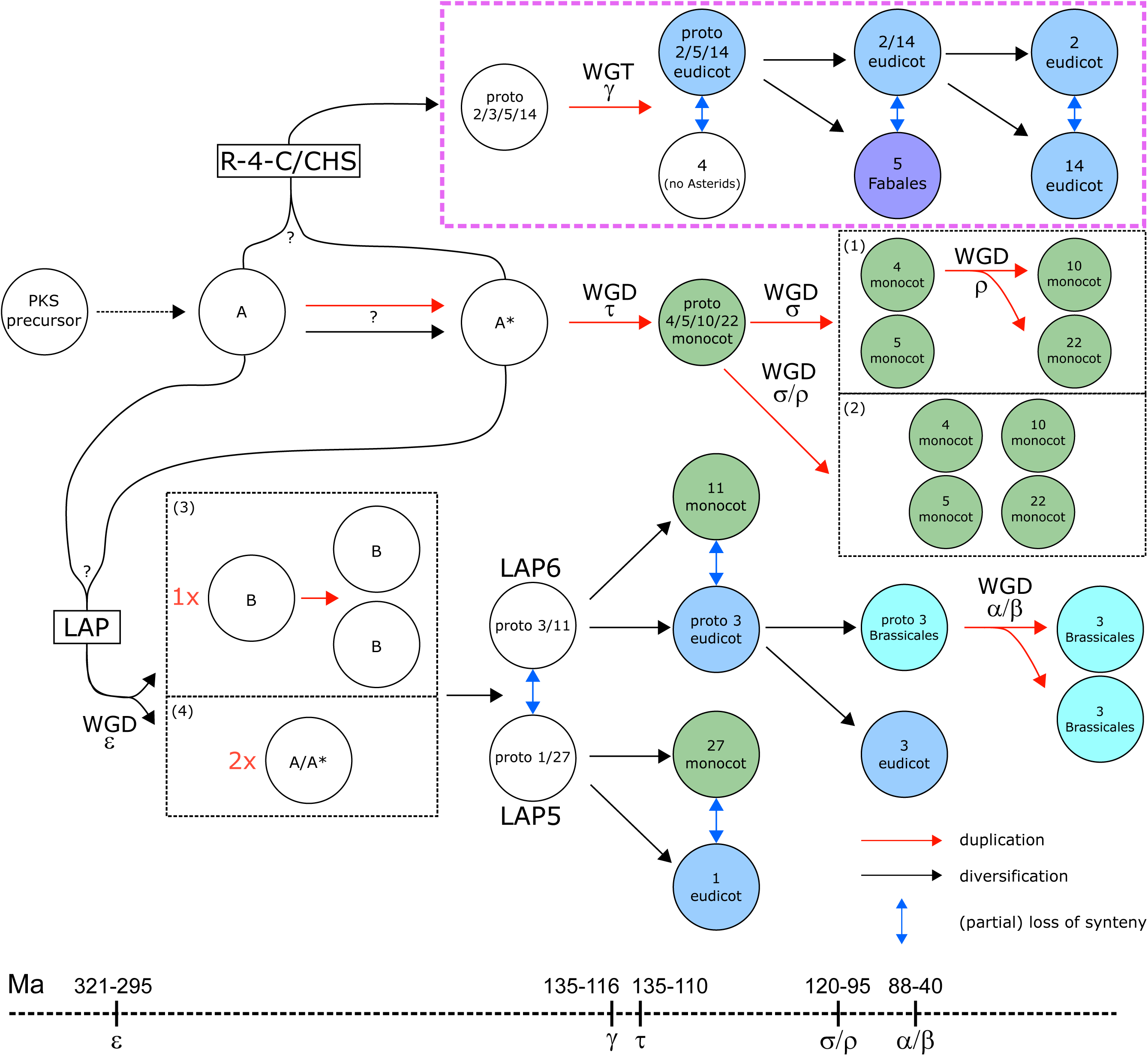
Possible route of evolution of the type III PKS superfamily in the plant kingdom. Given the data, it is unclear if the protocluster evolved into the ‘R-4-C’-containing clade or the LAP5/6 ortholog clade, first. Possible whole-genome duplications/triplications are indicated according to Van de Peer et al. (83) and Clark & Donoghue (82) based on the species distribution, the synteny network and phylogenetic analysis. Only major syntenic clusters are displayed. The location of clusters does approximately indicate the timing. Hypothesis for evolution of CHS: from protocluster A* diversification into protocluster 2/3/5/14, duplication of protocluster and diversification resulting into cluster 3 and protocluster 2/5/14, diversification of protocluster 2/5/14 and loss of synteny between the three clusters. Hypotheses for evolution of monocot-specific cluster: duplication of A* and diversification into proto 4/5/10/22 cluster, (1) duplication of proto cluster into 4/10/22 and 5, followed by a duplication of 4 resulting in 10 and 22; (2) duplication of monocot-specific proto cluster into 4, 5, 10 and 22. Hypotheses for Evolution of LAP5/6 ortholog clade: (3) 1x duplication of A or A* and diversification into protocluster B, subsequent duplication and divergence of this region and loss of synteny. (4) 2x duplication of protoclusters A or A* and loss of synteny between duplicated regions.

After the divergence from the Bryophyta and Gymnosperms, the LAP5 and LAP6-containing clades (including the Angiosperm-specific clusters 1, 3, 11 and 27) formed by a gene region/genome duplication event, possibly the ε whole-genome duplication event 321-295 million years ago (82-84), followed by diversification into eudicot-(1 and 3) and monocot-specific (11 and 27) clusters. Within the Brassicales, possibly the α or β whole-genome duplication event formed the syntenic clusters 14 and 15 88-40 million years ago (82, 83, 85, 86).

The members of the LAP5/6-specific clade are anther-specific and involved in sporopollenin biosynthesis (18, 19). Orthologs can also be found in members of the Bryophyta, Marchantiophyta, Lycopodiophyta, Pinophyta, Ginkgophyta and Gnetophyta families. This suggests that LAP5/6 orthologous sequences evolved before the divergence between the former and the Angiosperms by a segmental duplication event. The divergence between Gymnosperm- and Angiosperm-forming clades was 365.0-330.9 Million years ago in the Carboniferous (87, 88), the divergence between the Bryophyte- and the Tracheophyta-forming) clades occurred 506.4-460.3 Million years ago (88). Since *PKS* can be found in all land plant lines, it can be concluded that the type III *PKS* superfamily is at least 460.3-506.4 million years old.

The two major ‘R-4-C’ containing regions proto 2/5/14 (containing no ‘R-4-C’ sequences from Monocots) and 4 evolved possibly by the γ whole-genome triplication event 135-116 million years ago after the divergence of the Eudicots and the Monocots (82, 83, 89). Intriguingly, we found that the CHSs of *Glycine max* interacting with chalcone reductase (90) locate to syntenic cluster 5 suggesting that a duplication event in the Fabales leading to syntenic cluster 5 facilitated the biosynthesis of isoflavonoids (see Supplementary Text for details).

The monocot-specific clades of syntenic clusters 4, 5, 10 and 22 possibly evolved by τ whole-genome duplication 135-110 million years ago (91), followed by σ and ρ whole-genome duplication (see (1) in Fig. 5) *or* by the σ or ρ whole-genome-duplication in the Monocot lineage (see (2) in Fig. 5) 120-95 million years ago (82, 83, 91, 92).

The question remains, which clade evolved directly from the protocluster A, the LAP5/6 clade or the clade containing syntenic clusters 2, 4, 5 and 14 and others. It is interesting to note that sequences of *Azolla filiculoides* (syntenic cluster 4), *Marchantia polymorpha* (5), and *Physcomitrella patens* (5, 15), Salvinia cucullata (4) and Selaginella moellendorffii (4) are both located in the phylogenetic clade containing the syntenic clusters 2, 4, 5 and 14 *and* containing the LAP5/6 homologs.. Most of the sequences in the syntenic cluster 4, 5 and 15 co-locate to the phylogenetic clade containing the syntenic clusters 2, 4, 5 and 14 and show strong synteny to ‘R-4-C’-containing syntenic clusters and less to the LAP-containing syntenic clusters 1 and 3. This favors the hypothesis that the syntenic cluster 4 and 5 is primordial and existed before the emergence of LAP5/6 containing syntenic clusters, albeit it has to be kept in mind that not all sequences of the Bryophyta, Lycopodiophyta, Marchantiophyta and Polypodipsida showed synteny to other sequences and that the loss of synteny between clusters is similar across different clusters.

Studying the macroevolution of the type III PKS superfamily, the clade containing 2, 4, 5 and 14 *or* the clade containing LAP5/6 evolved from protocluster A *and* the respective other clade evolved by a genome (region) duplication event from protocluster A or A*. Another possibility, although less favored, is that two genome (region) duplication events from A or A* occurred forming the clade containing LAP5/6 and the syntenic clusters 2, 4, 5 and 14. The ‘R-4-C’-type enriched syntenic clusters 2, 5 and 14 do not contain monocot ‘R-4-C’ sequences indicating that the diversification into eudicot ‘R-4-C’ happened after the divergence from the monocots *or* that monocot PKS genes in the syntenic clusters 2, 5 and 14 lost their ‘R-4-C’ function after diverging from the eudicots. Next to their macroscale evolution, we observed gene expression changes for CHS orthologs and conservation of expression pattern for LAP5/6 orthologs (see Supplementary Text for details) indicating a diversification of CHS orthologs after duplication events.

Sporopollenin is a constituent of the spore and pollen grain outer walls of all known land plants. The average pine sporopollenin structure consists of two fatty acid-derived polyvinyl alcohol-like units, each flanked at one end by a α-pyrone at one end and cross-linked by an ester at the other end. Sporopollenin furthermore possesses supposedly covalently linked *p*-coumaric acid and naringenin as structural units (93). Tri- and tetraketide α-pyrones are formed by LAP5 and LAP6 proteins from a broad range of potential acyl-CoA synthetase 5-synthesized fatty acyl-CoA starter substrates (18, 19)Sporopollenin, might have equipped algal zygotes with a UV-protecting outer layer that promoted their movement onto the land (7, 94).As some of the most primitive organisms, some members of the freshwater algae Charophytes are believed to host the phenylpropanoid pathway (94, 95) and it has been suggested that the UV autofluorescent lignin-like material surrounding the zygotes of several charophytic algae species, including *Coleochaete*, is sporopollenin (7, 95, 96). By contrast, Zygnematophyceae zygospores contain algaenan that differs from pollen grains in its chemical position and the biochemical pathway (acetate-malate pathway) leading to its production (97). Furthermore, to our knowledge, the presence of type III PKS-like enzymes in the genomes of Charophyta is not reported with the exception of *Penium margaritaceum* (36). The adaption of a sporopollenin-containing protecting spore wall is considered a synapomorphy of the embryophytes to colonize the land, but appears to be pre-adaptive given that it is present in the Charophyceae, the proposed sister group to the embryophytes (7, 94-96, 98).If LAP5/6 evolved first, the syntenic clusters 1/27 and/or 3/11 duplicated and formed the cluster that later the syntenic cluster that contain ‘R-4-C’-type *PKS* genes followed by a loss of synteny between the clusters 1/27 and 3/11. This sequence of events is consistent with the work of Weng & Chapple (7) who postulated that the emergence of sporopollenin biosynthesis occurred earlier than that of phenylpropanoid metabolism and flavonoid biosynthesis. To our knowledge, it is not clear if sporopollenin biosynthesis required at this point of emergence the presence of α-pyrone. With the data currently at hand, the exact sequence of events can however not be elucidated fully.

One obstacle that remains is that there is no syntenic information currently available on gymnosperm species, due to their large genome sizes and technical difficulties in their assembly. Such information would allow us to refine the evolutionary sequence we presented here to a higher resolution than is currently possible. This fact notwithstanding, we feel that the study here has allowed us to carry out a comprehensive analysis, and one that is unprecedented in scope of the evolution of the type III PKS family in a manner which we believe is highly applicable to myriad of other specialized pathways of the plant kingdom and beyond.

## Methods

### Retrieval of genomic data and processing of proteome files

Protein FASTA files and .gff/.gff3 files were downloaded for 126 species from the sources indicated in Supplementary file 2. If available, functional annotation files (containing GO annotation and InterPro domains) were downloaded from the same sources. Splice variants, if any were annotated, were removed retaining only the variant with the longest coding sequence for each locus.

Annotated transposable elements in the genomes were removed. Additionally, to further remove TEs and remove TEs in genomes where they were not annotated, a local peptide library was built containing known *Arabidopsis thaliana, Oryza sativa, Solanum lycopersicum* and *Zea mays* transposable elements. All species’ protein FASTA files were queried (using BLAST, default settings) against this database and hits were considered a transposable elements, and removed as well, when the protein identity was > 70%, the E-value < 0.05 AND the length > 50. The proteome files were checked for completeness using BUSCO(99) (v4.0.2_cv1, -m proteins, –l chlorophyta_odb10, lineage dataset from 2019-11-20, for the species *Chara braunii, Chlamydomonas reinhardtii, Coccomyxa sp. C169, Cyanidioschyzon merolae, Cyanophora paradoxa, Dunaliella salina, Klebsormidium nitens, Ostreococcus lucimarinus, Volvox carteri, -l stramenopiles_odb10*, lineage dataset from 2019-11-21, for the species *Ectocarpus siliculosus* and *Aureococcus aneophagefferens,* -l cyanobacteria_odb10, lineage dataset from 2919-04-24, *Synechocystis sp. PCC 6803* or -l embryophyta_odb10, lineage dataset from 2019-11-20, for all other species) in the respective docker container.

### Inference of orthogroups, orthologs and gene families using OrthoFinder and MCL

Orthogroups were inferred from protein FASTA files (proteome files) using OrthoFinder(34) (v2.2.7), Python (v2.7.10), diamond (v0.9.9), dlcpar (v1.0), fastme (v2.1.5) and mcl (v14.137) using the command orthofinder.py -f ./ -S diamond. To obtain MCL groups, pairwise-species BLAST files were used as input for MCL(35) clustering using mcxload (--stream-mirror, –stream-neg-log10, -stream-tf ‘ceil (200)’, abc file from BLAST results) and mcl (-I 2) (https://micans.org/mcl/).

### Detection of syntenic regions using i-ADHoRe and MCScanX

For each species, information on the gene orientation (+/-) was extracted from the .gff/.gff3 and one file per scaffold/chromosome was created containing the gene (matching the identifier in the protein FASTA file) and its orientation according to the order in the genome. i-ADHoRe (v3.0.01) was used to detect collinear regions between two genomes using the following settings within the .ini file: table_type=family, cluster_type=collinear, alignment_method=gg2, gap_size=15, cluster_gap=20, max_gaps_in_alignment=20, q_value=0.9, prob_cutoff=0.001, anchor_points=5, level_2_only=true, write_stats=true, number_of_threads=4. For blast_table the output from OrthoFinder or MCL clustering was used, respectively, where each protein (in the first column) referred to an orthogroup/group (in the second column). MCScanX (mcscanx_h, version 3-28-2013) detected collinear regions between two genomes (using –b 0 option) using the homology relations from OrthoFinder or MCL clustering.

### Annotation of PKS genes

PKS protein sequences were blasted against the NCBI database and the fit with lowest E-value was reported for annotation (expect threshold: 10; word size: 6; matrix: BLOSUM62; Gap costs: existence: 11, Extension: 1; Compositional adjustments: Conditional compositional score matrix adjustment). The same parameters were used when blasting other sequences using blastp against the NCBI database. Classification of the PKS type III reaction type was predicted via pPAP(28) (v1.1) using the protein sequences as input (ruby v2.5.1p57, BioRuby 1.5.2, MAFFT v7.310 and HMMER v3.2.1). The number of exons was taken from the PLAZA database (Dicots PLAZA 4.0, Monocots PLAZA 4.0, Gymno PLAZA 1.0, pico-PLAZA 2.0).

To retrieve type III PKS sequences in the species *Chlorokybus atmophyticus* (accession no. RHPI00000000) and *Mesostigma viride* (accession no. RHPH00000000)(66), known CDS sequences of *Penium margaritaceum* were blasted against the assemblies of the two species using the following options ‘Optimize for: Somewhat similar sequences (blastn)’, ‘database: ASM910322v1 GenBank assembly GCA_009103225.1’ (*Chlorokybus atmophyticus*) / ‘database: ASM974604v1 GenBank assembly GCA_009746045.1’ (*Mesostigma viride*), ‘Match score: 2’, ‘Mismatch score -3’, ‘Gap costs: Existence: 5, Extension: 2’, ‘Word size: 7’, ‘filter low complexity regions’, ‘mask for lookup table only’. To retrieve type III PKS sequences of members of the Coleochaetaphyceae, we queried the CDS sequences against the nucleotide collection of the Coleochaetaphyceae (taxid: 131209, other algorithm parameters identical as for *Chlorokybus atmophyticus* and *Mesostigma viride*).

### Tandem gene identification and syntenic network construction and clustering

Analysis of synteny was done according to Zhao & Schranz(32) following a network approach using a custom script. Tandem genes were defined as present when they were detected in one of the four methods and all tandem genes per region (genes that form a component) were treated as a tandem gene region in the following. In a next step and for each method, the value 0.25 was added to a_i,j_ to adjacency matrix A, if syntenic link between (tandem) gene regions i and j, containing PKS genes, exists. Connections of type “i-ADHoRe+MCL and “MCScanX+OrthoFinder” *and* “i-ADHoRe+OrthoFinder and MCScanX+MCL” were removed from the adjacency matrices. Vertices that do not link to others were removed. To determine clusters of the syntenic network, four community structure detection algorithms (all algorithms resulted in ≤20 clusters) were applied separately on the network to retrieve membership: based on greedy optimization of modularity (function fastgreedy.community, modularity=TRUE), via short random walks (function walktrap.community, modularity=TRUE, steps=15), based on the leading eigenvector of the community matrix (function leading.eigenvector.community, steps=15), and based on multi-level optimization of modularity (function multilevel.community, all functions from igraph package v1.2.4.1, R v3.5.0). After this step distances were calculated for each cluster: if cluster assignment was identical, distance was set to 0, otherwise to 1. The final cluster membership was obtained by affinity propagation clustering for cluster detection using the information from all six cluster detection algorithms (apcluster from the apcluster package, v1.4.6, convits=1000, maxits=10000, lam=0.9, nonoise=TRUE). The enrichment test for ‘R-4-C’-type PKS in clusters 2, 4, 5 and 14 (selection criteria: more than 15% of genes are of type ‘R-4-C’ and at least 10 genes in cluster) was performed with the function fisher.test (alternative=“greater”) in the R environment (v3.5.0) after removing the terms for sequences that were not present in the syntenic network. The custom script for network construction can be accessed via www.github.com/tnaake/PKS_synteny.

### Phylogenetic analysis

A multiple sequence alignment were built from characterized PKS protein sequences using MUSCLE (v3.8.31). A HMM protein profile was built using hmmbuild (--fragthresh 0, hmmer v3.2.1). Using hmmsearch the HMM protein profile was queried against the proteome files. Sequences, from sequences for which no proteome file is available, were added manually by NCBI database research. Hits were aligned with the protein profile using hmmalign and the alignment was manually checked. Columns with >20% missing values were excluded from further analysis, as well as frayed C- and N-terminal regions of the alignment (Supplementary File 4). Tree building was done by raxmlHPC-AVX (-f a, -m PROTGAMMALGX, -c 25, -p 12345, -x 12345) using 1000 bootstrap replicates and the genes AAK45681, BAD97390, BAA33495 as outgroups. booster(42) calculated transfer bootstrap expectation values for branches (-a tbe).

Visualization of the tree was performed within the R environment (v3.5.0) and ggtree (v1.17.1). The phylogenetic species tree was obtained by OrthoFinder via species tree inference from All Genes (STAG)(100) using the processed proteome files and midpoint rooting in FigTree (v1.4.3).

### Enrichment analysis

Gene ontology terms were obtained for each species separately by using the processed proteome FASTA files and PANNZER2(101, 102) entering the species name in the field ‘Scientific name of query species’. Subsequently, three types of enrichment analyses were run: (1) enrichment of genes of syntenic regions containing PKS genes with no removal of PKS genes using the syntenic genes of syntenic regions as background, (2) enrichment of genes of syntenic regions containing PKS with removal of PKS genes using the syntenic genes of syntenic regions as background, (3) enrichment of genes of syntenic regions containing PKS genes with no removal of PKS genes using all genes as background (syntenic genes of syntenic regions and other genes), (4) enrichment of genes of syntenic regions containing PKS genes and all genes as background. GO terms of genes in PKS-containing syntenic regions were tested against GO terms of the backgrounds (genes of all syntenic regions or all genes, PKS-BG), genes of ‘R-4-C’-enriched syntenic regions of clusters 2, 4, 5 and 14 were tested against background (CHS-PKS) and genes of ‘R-4-C’-enriched syntenic regions of clusters 2, 4, 5 and 14 were tested against GO terms of genes within PKS-containing syntenic regions (CHS-PKS) using fisher.test (alternative=“greater”) within R (v3.5.0). The enrichment analyses were separately conducted for PKS-BG, CHS-BG and CHS-PKS and p-values were adjusted by Benjamini-Hochberg using p.adjust within the R environment (v3.5.0). Enriched terms were visualized in Cytoscape(103) (v3.6.1) using Enrichment Map(104) (FDR q-value cutoff=0.2, p-value cutoff=0.5, NES (GSEA only)=All, Data Set Edges=Combine edges across data sets (sparser), Cutoff=0.375, Metric=Jaccard+Overlap combined, Jaccard=50%, Overlap=50%). The custom script for enrichment analysis can be accessed via www.github.com/tnaake/PKS_synteny.

### STRING DB

Protein sequence FASTA files were obtained for the genes in PKS-containing syntenic regions from the union of all four methods (i-ADHoRe+OrthoFinder, i-ADHoRe+MCL, MCScanX+OrthoFinder, MCScanX+MCL) for all PKS-containing syntenic regions from *A. thaliana, S. lycopersicum, O. sativa, Z. mays* and *V. vinifera*. Query sequences with highest identity to STRING proteins were taken as the mapping candidate. Co-expression within syntenic regions were checked by using the STRING DB(105) with the following settings: meaning of network edges=confidence, active interactive sources=Co-expression, minimum required interaction score=medium (0.400), max number of interactors to show: 1st shell=none, 2nd shell=none).

### Gene expression CoNekT database

Raw expression values were downloaded for PKS sequences from the CoNekT database(106) (retrieved April 25, 2019) for the species *Arabidopsis thaliana, Oryza sativa, Selaginella moellendorffii, Solanum lycopersicum, Vitis vinifera* and *Zea mays*. Sampling conditions were categorized into roots/rhizoids, leaves, stem/shoot, fruit/siliques/ear/strobilus/spores, seed, flower, pollen and mean values from raw expression values per category were calculated for each gene. Pearson correlation values were calculated between averaged gene expression values using the cor function within R (v3.5.0). Pearson correlation values were clustered by affinity propagation clustering using apcluster (apcluster package, convits=1000, maxits=10000, nonoise=TRUE seed=1000) in R (v3.5.0).

## Acknowledgements

We would like to thank Stefanie Hartmann for discussion about phylogenetic analysis. We thank Iben Sorensen and Jocelyn Rose for providing the PKS sequences of *Penium margaritaceum*. We would like to thank Camilla Ferrari for help in gathering the genomic data, Nooshin Omranian for helpful discussion about network analysis and Federico Scossa for discussion about phylogenetic analysis. T.N. acknowledges the financial support of the IMPRS-PMPG program. S.P. is supported by VLAIO grant HBC.2016.0556. The work of H.A.M. was supported by National Science Foundation grants IOS-1836824 and MCB-1818040.

## Author contributions

T.N., T.T., H.A.M and A.R.F.designed the research. T.N. performed and analyzed the transcriptome data and co-expression data. T.N., and A.R.F. performed the phylogenetic analysis. T.N. and S.P. performed synteny network analysis. T.N. performed enrichment analysis. T.N. and H.A.M. analyzed enrichment results. T.N. and A.R.F. wrote the manuscript with input from all authors.

