## Supplementary Figures, Tables and Text for "Kingdom-wide analysis of the evolution of the plant type III polyketide synthase superfamily"

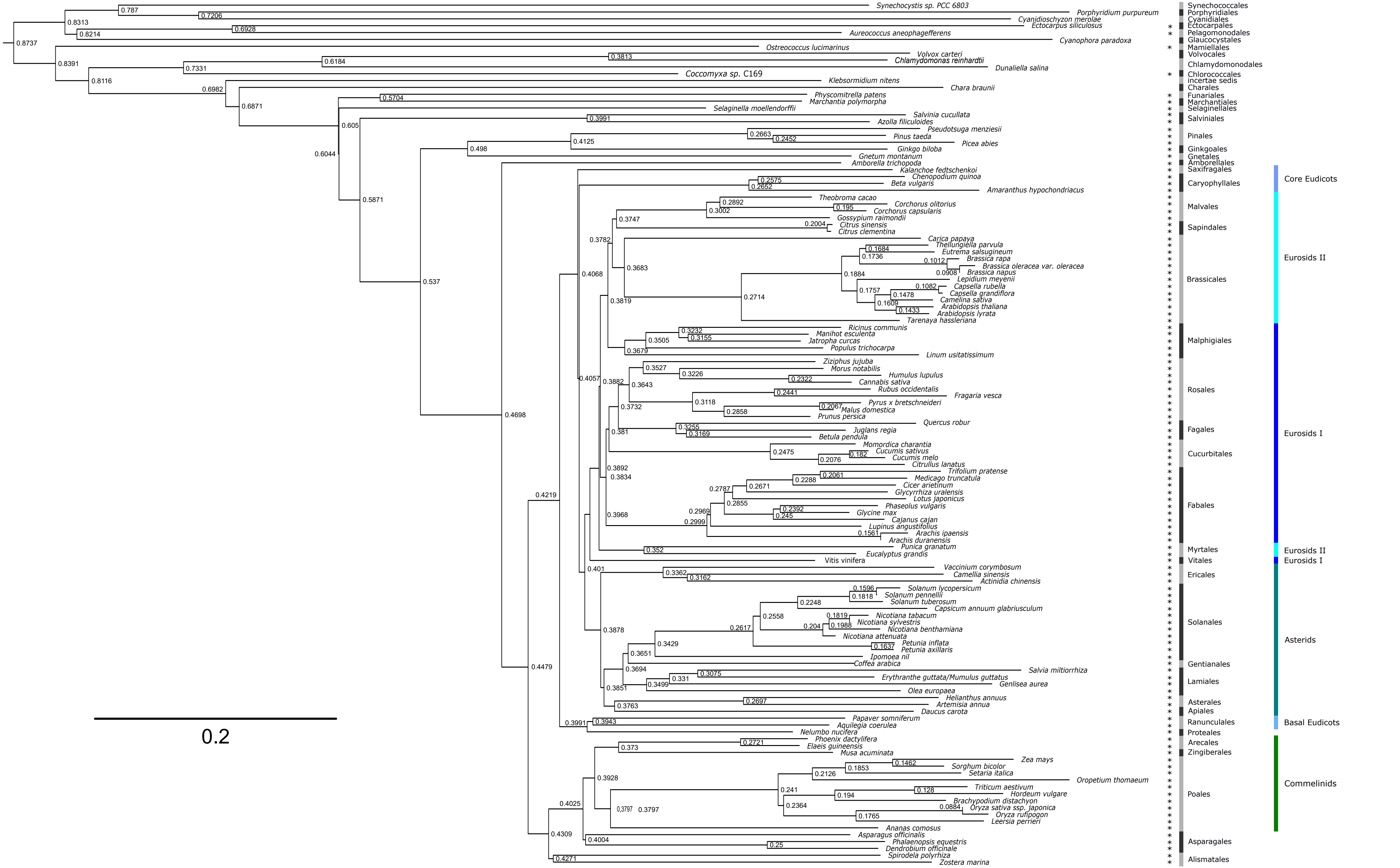

Synechocystis sp. PCC 6803
Porphyridium purpureum
Cyanidioschyzon merolae
Ectocarpus siliculosus
Aureococcus aneophagefferens
Cyanophora paradoxa
Ostreococcus lucimarinus
Volvox carteri
Chlamydomonas reinhardtii
Dunaliella salina
Coccomyxa sp. C169
Klebsormidium nitens
Chara braunii
Physcomitrella patens
Marchantia polymorpha
Selaginella moellendorffii
Salvinia cucullata
Azolla filiculoides
Pseudotsuga menziesii
Pinus taeda
Picea abies
Ginkgo biloba
Gnetum montanum
Amborella trichopoda
Kalanchoe fedtschenkoi
Chenopodium quinoa
Beta vulgaris
Amaranthus hypochondriacus
Theobroma cacao
Corchorus olitorius
Corchorus capsularis
Gossypium raimondii
Citrus sinensis
Citrus clementina
Carica papaya
Thellungiella parvula
Eutrema salsugineum
Brassica rapa
Brassica oleracea var. oleracea
Brassica napus
Lepidium meyenii
Capsella rubella
Capsella grandiflora
Camelina sativa
Arabidopsis thaliana
Arabidopsis lyrata
Tarenaya hassleriana
Ricinus communis
Manihot esculenta
Jatropha curcas
Populus trichocarpa
Linum usitatissimum
Ziziphus jujuba
Morus notabilis
Humulus lupulus
Cannabis sativa
Rubus occidentalis
Fragaria vesca
Pyrus x bretschneideri
Malus domestica
Prunus persica
Quercus robur
Juglans regia
Betula pendula
Momordica charantia
Cucumis sativus
Cucumis melo
Citrullus lanatus
Trifolium pratense
Medicago truncatula
Cicer arietinum
Glycyrrhiza uralensis
Lotus japonicus
Phaseolus vulgaris
Glycine max
Cajanus cajan
Lupinus angustifolius
Arachis ipaensis
Arachis duranensis
Punica granatum
Eucalyptus grandis
Vitis vinifera
Vaccinium corymbosum
Camellia sinensis
Actinidia chinensis
Solanum lycopersicum
Solanum pennellii
Solanum tuberosum
Capsicum annuum glabriusculum
Nicotiana tabacum
Nicotiana sylvestris
Nicotiana benthamiana
Nicotiana attenuata
Petunia inflata
Petunia axillaris
Ipomoea nil
Coffea arabica
Salvia miltiorrhiza
Erythranthe guttata/Mumulus guttatus
Genlisea aurea
Olea europaea
Helianthus annuus
Artemisia annua
Daucus carota
Papaver somniferum
Aquilegia coerulea
Nelumbo nucifera
Phoenix dactylifera
Elaeis guineensis
Musa acuminata
Zea mays
Sorghum bicolor
Setaria italica
Oropetium thomaeum
Triticum aestivum
Hordeum vulgare
Brachypodium distachyon
Oryza sativa ssp. japonica
Oryza rufipogon
Leersia perrieri
Ananas comosus
Asparagus officinalis
Phalaenopsis equestris
Dendrobium officinale
Spirodela polyrhiza
Zostera marina
Synechococcales
Porphyridiales
Cyanidiales
Ectocarpales
Pelagomonodales
Glaucocystales
Mamiellales
Volvocales
Chlamydomonodales
Chlorococcales
incertae sedis
Charales
Funariales
Marchantiales
Selaginellales
Salviniales
Pinales
Ginkgoales
Gnetales
Amborellales
Saxifragales
Caryophyllales
Malvales
Sapindales
Brassicales
Malphigiales
Rosales
Fagales
Cucurbitales
Fabales
Myrtales
Vitales
Ericales
Solanales
Gentianales
Lamiales
Asterales
Apiales
Ranunculales
Proteales
Arecales
Zingiberales
Poales
Asparagales
Alismatales
Core Eudicots
Eurosids II
Eurosids I
Eurosids II
Eurosids I
Asterids
Basal Eudicots
Commelinids
0.2

Supplementary Figure 1: Phylogenetic tree for 126 analyzed species. PKS signatures are present in all vascular plants. The species tree was inferred from all genes (STAG) according to Emms & Kelly (2019)[1]. Branch length represents the average number of substitution per sites across a gene families. Support value for each bipartition in the consensus STAG tree are the proportion of times that the bipartition is seen in each of the individual species tree estimates. Scale represents substitutions per site. The outgroup was defined by midgroup rooting in FigTree. *: PKS signature present in species as given per OrthoFinder and/or MCL analysis.

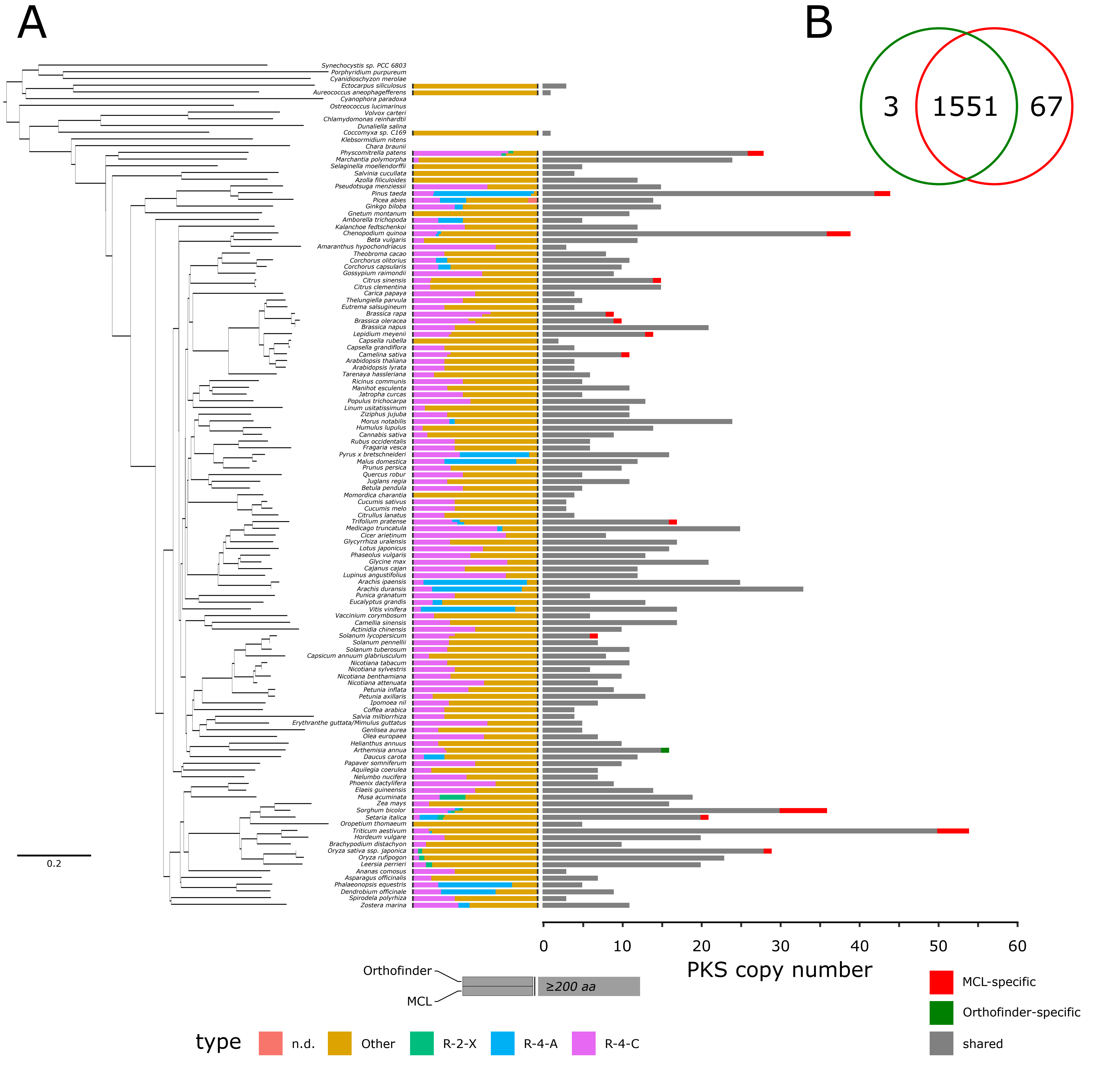
A
B
3
1551
67
0.2
0
10
20
30
40
50
60
PKS copy number
Orthofinder
MCL
≥200 aa
MCL-specific
Orthofinder-specific
shared
type
n.d.
Other
R-2-X
R-4-A
R-4-C

Supplementary Figure 2: Number of PKS genes in analyzed species. A: All analyzed vascular plants showed more than two copies of PKS signatures in their genome. Species with a high number of type III PKS belonged to the clades Fabales, Poacea, gymnosperms, Marchantiales, Sphagnales and Funariales. The species are sorted according to their phylogeny. The type of the PKS according to pPAP classification is displayed in the middle part of the panel (all signatures with length ≤ 200 amino acids removed) on a relative scale and the number of PKS signatures within the genome is displayed in the right panel for OrthoFinder and MCL separately. B: Number of PKS genes according to OrthoFinder (in total 1,554 signatures) and MCL (1,618 signatures). 1,551 genes are shared between the two detection algorithms.

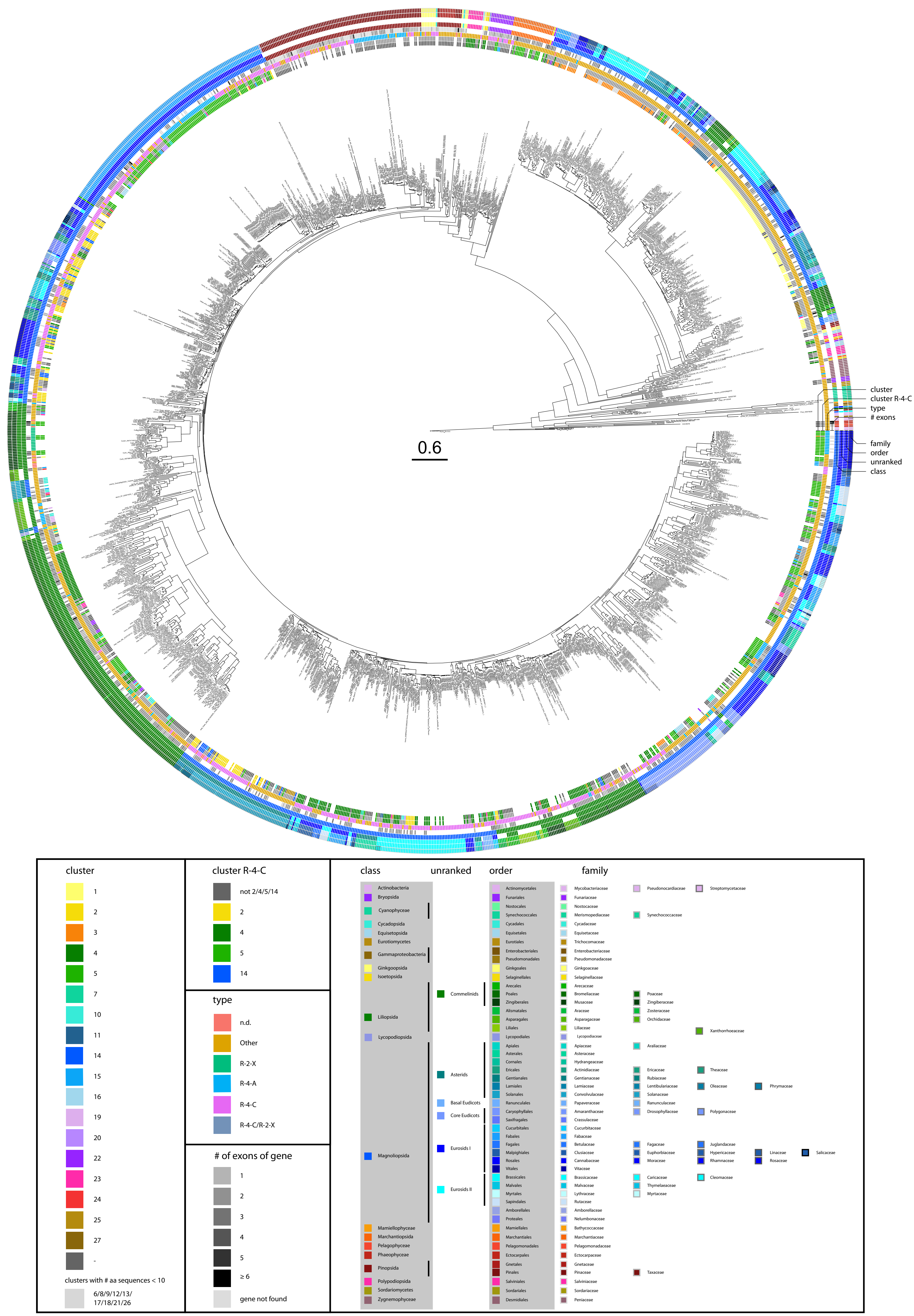

0.6
cluster
cluster R-4-C
type
### exons
family
order
unranked
class
cluster
1
2
3
4
5
7
10
11
14
15
16
19
20
22
23
24
25
27
-
clusters with # aa sequences < 10
6/8/9/12/13/
17/18/21/26
cluster R-4-C
not 2/4/5/14
2
4
5
14
type
n.d.
Other
R-2-X
R-4-A
R-4-C
R-4-C/R-2-X
### of exons of gene
1
2
3
4
5
≥ 6
gene not found
class
Actinobacteria
Bryopsida
Cyanophyceae
Cycadopsida
Equisetopsida
Eurotiomycetes
Gammaproteobacteria
Ginkgoopsida
Isoetopsida
Liliopsida
Lycopodiopsida
Magnoliopsida
Mamiellophyceae
Marchantiopsida
Pelagophyceae
Phaeophyceae
Pinopsida
Polypodiopsida
Sordariomycetes
Zygnemophyceae
unranked
Commelinids
Asterids
Basal Eudicots
Core Eudicots
Eurosids I
Eurosids II
order
Actinomycetales
Funariales
Nostocales
Synechococcales
Cycadales
Equisetales
Eurotiales
Enterobacteriales
Pseudomonadales
Ginkgoales
Selaginellales
Arecales
Poales
Zingiberales
Alismatales
Asparagales
Liliales
Lycopodiales
Apiales
Asterales
Cornales
Ericales
Gentianales
Lamiales
Solanales
Ranunculales
Caryophyllales
Saxifragales
Cucurbitales
Fabales
Fagales
Malpighiales
Rosales
Vitales
Brassicales
Malvales
Myrtales
Sapindales
Amborellales
Proteales
Mamiellales
Marchantiales
Pelagomonadales
Ectocarpales
Gnetales
Pinales
Salviniales
Sordariales
Desmidiales
family
Mycobacteriaceae
Pseudonocardiaceae
Streptomycetaceae
Funariaceae
Nostocaceae
Merismopediaceae
Synechococcaceae
Cycadaceae
Equisetaceae
Trichocomaceae
Enterobacteriaceae
Pseudomonadaceae
Ginkgoaceae
Selaginellaceae
Arecaceae
Bromeliaceae
Poaceae
Musaceae
Zingiberaceae
Araceae
Zosteraceae
Asparagaceae
Orchidaceae
Liliaceae
Xanthorrhoeaceae
Lycopodiaceae
Apiaceae
Araliaceae
Asteraceae
Hydrangeaceae
Actinidiaceae
Ericaceae
Theaceae
Gentianaceae
Rubiaceae
Lamiaceae
Lentibulariaceae
Oleaceae
Phrymaceae
Convolvulaceae
Solanaceae
Papaveraceae
Ranunculaceae
Amaranthaceae
Drosophyllaceae
Polygonaceae
Crassulaceae
Cucurbitaceae
Fabaceae
Betulaceae
Fagaceae
Juglandaceae
Clusiaceae
Euphorbiaceae
Hypericaceae
Linaceae
Salicaceae
Cannabaceae
Moraceae
Rhamnaceae
Rosaceae
Vitaceae
Brassicaceae
Caricaceae
Cleomaceae
Malvaceae
Thymelaeaceae
Lythraceae
Myrtaceae
Rutaceae
Amborellaceae
Nelumbonaceae
Bathycoccaceae
Marchantiaceae
Pelagomonadaceae
Ectocarpaceae
Gnetaceae
Pinaceae
Taxaceae
Salviniaceae
Sordariaceae
Peniaceae

Supplementary Figure 3: Phylogenetic gene tree of type III PKS amino acid sequences with additional information on syntenic cluster membership, type of sequence according to pPAP classification, number of exons of the gene sequence, and taxonomic information on the family, order, unranked taxonomic information and class of the species to which the sequences refers to. Proteome files of 126 species were queried for type III PKS sequences via a HMM profile of known PKS sequences. For species, for which proteome files were not available, PKS genes were selected based on previous annotation. Amino acid sequences were aligned via hmmalign and sites with more than 20% missing values were removed. The gene tree containing 1607 unique sequences were build using RAxML using 1000 bootstrap replications. The tree shows high transfer bootstrap expectation values[2] for all major clades.

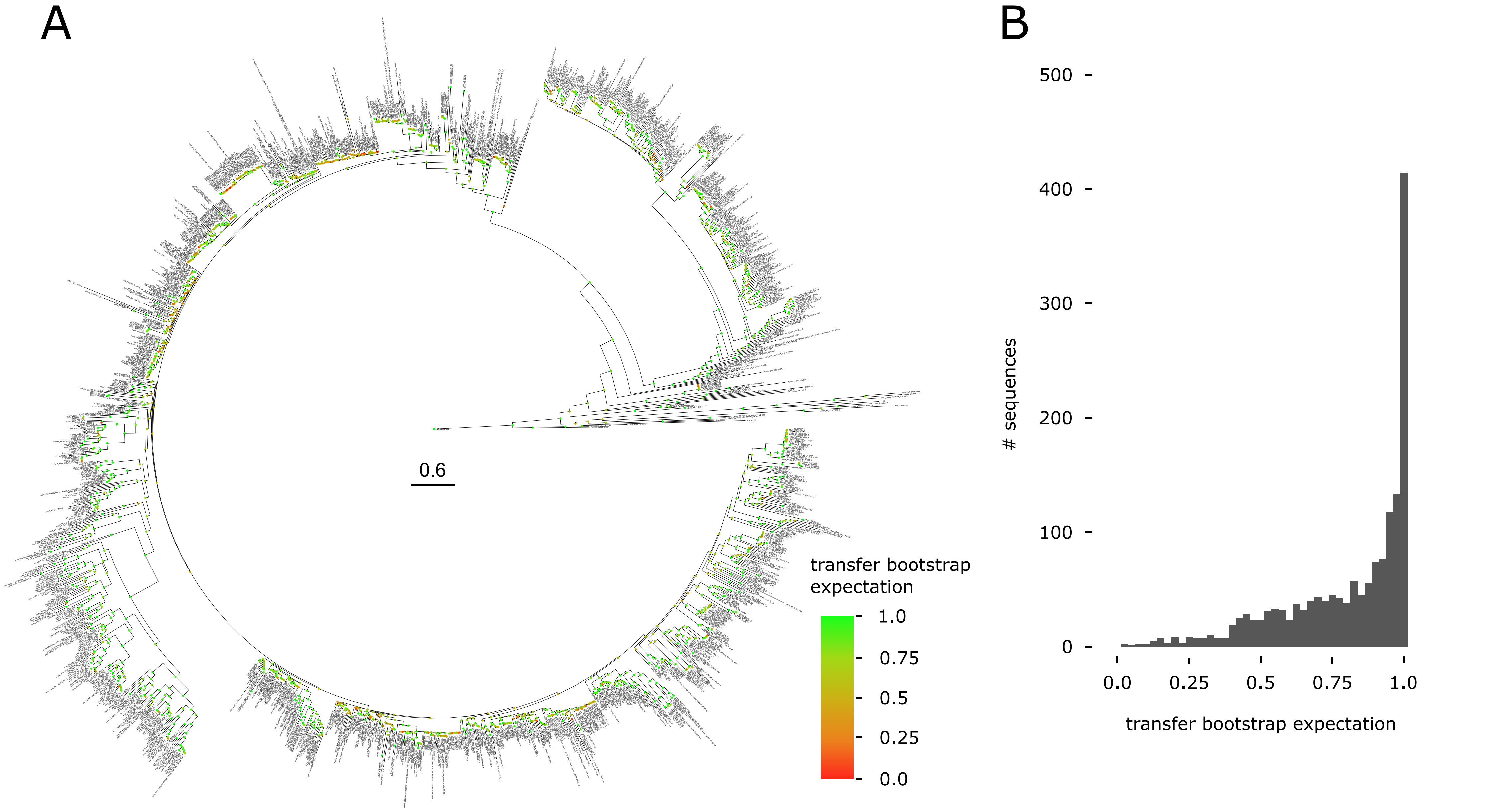
A
0.6
transfer bootstrap expectation
1.0
0.75
0.5
0.25
0.0
B
500
400
300
200
100
0
### sequences
0.0
0.25
0.5
0.75
1.0
transfer bootstrap expectation

Supplementary Figure 4: Transfer bootstrap expectation values for phylogenetic gene tree of type III PKS amino acid sequences. A: Phylogenetic tree with overlaid transfer bootstrap expectation values on nodes. The phylogenetic tree shows high bootstrap values on all major clades. B: Distribution of transfer bootstrap expectation values of the complete tree nodes. Transfer bootstrap expectation values were calculated according to Lemoine et al.[2].

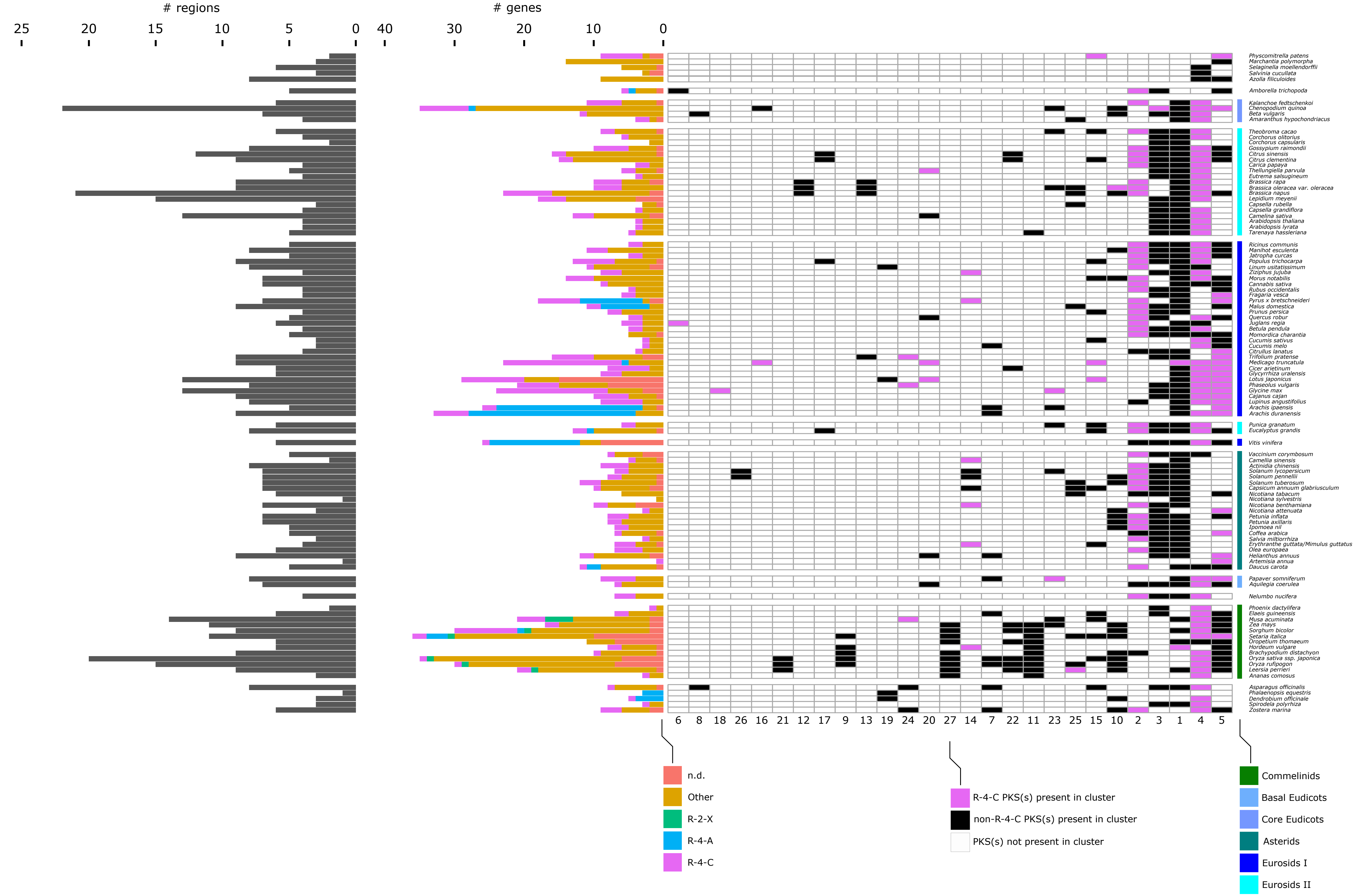

### regions
25 20 15 10 5 0
### genes
40 30 20 10 0
Physcomitrella patens
Marchantia polymorpha
Selaginella moellendorffii
Salvinia cucullata
Azolla filiculoides
Amborella trichopoda
Kalanchoe fedtschenkoi
Chenopodium quinoa
Beta vulgaris
Amaranthus hypochondriacus
Theobroma cacao
Corchorus olitorius
Corchorus capsularis
Gossypium raimondii
Citrus sinensis
Citrus clementina
Carica papaya
Thellungiella parvula
Eutrema salsugineum
Brassica rapa
Brassica oleracea var. oleracea
Brassica napus
Lepidium meyenii
Capsella rubella
Capsella grandiflora
Camelina sativa
Arabidopsis thaliana
Arabidopsis lyrata
Tarenaya hassleriana
Ricinus communis
Manihot esculenta
Jatropha curcas
Populus trichocarpa
Linum usitatissimum
Ziziphus jujuba
Morus notabilis
Cannabis sativa
Rubus occidentalis
Fragaria vesca
Pyrus x bretschneideri
Malus domestica
Prunus persica
Quercus robur
Juglans regia
Betula pendula
Momordica charantia
Cucumis sativus
Cucumis melo
Citrullus lanatus
Trifolium pratense
Medicago truncatula
Cicer arietinum
Glycyrrhiza uralensis
Lotus japonicus
Phaseolus vulgaris
Glycine max
Cajanus cajan
Lupinus angustifolius
Arachis ipaensis
Arachis duranensis
Punica granatum
Eucalyptus grandis
Vitis vinifera
Vaccinium corymbosum
Camellia sinensis
Actinidia chinensis
Solanum lycopersicum
Solanum pennellii
Solanum tuberosum
Capsicum annuum glabriusculum
Nicotiana tabacum
Nicotiana sylvestris
Nicotiana benthamiana
Nicotiana attenuata
Petunia inflata
Petunia axillaris
Ipomoea nil
Coffea arabica
Salvia miltiorrhiza
Erythranthe guttata/Mimulus guttatus
Olea europaea
Helianthus annuus
Artemisia annua
Daucus carota
Papaver somniferum
Aquilegia coerulea
Nelumbo nucifera
Phoenix dactylifera
Elaeis guineensis
Musa acuminata
Zea mays
Sorghum bicolor
Setaria italica
Oropetium thomaeum
Hordeum vulgare
Brachypodium distachyon
Oryza sativa ssp. japonica
Oryza rufipogon
Leersia perrieri
Ananas comosus
Asparagus officinalis
Phalaenopsis equestris
Dendrobium officinale
Spirodela polyrhiza
Zostera marina
6 8 18 26 16 21 12 17 9 13 19 24 20 27 14 7 22 11 23 25 15 10 2 3 1 4 5
n.d.
Other
R-2-X
R-4-A
R-4-C
R-4-C PKS(s) present in cluster
non-R-4-C PKS(s) present in cluster
PKS(s) not present in cluster
Commelinids
Basal Eudicots
Core Eudicots
Asterids
Eurosids I
Eurosids II

Supplementary Figure 5: Distribution of *PKS* genes in syntenic clusters. The panel shows a binary matrix in which it is indicated if one or more PKS or at least one 'R-4-C'-type PKS is present for a certain species given a syntenic cluster. The Clusters 2, 4, 5 and 14 are 'R-4-C'-enriched syntenic clusters. The clusters 1, 3, 11 and 27 contain *LAP5* and *LAP6* homologs. The clusters 11 and 27 show specificity for Commelinid species. The order of species is according to their phylogeny. On the left, the number of syntenic genes and the number of syntenic regions per species are displayed.

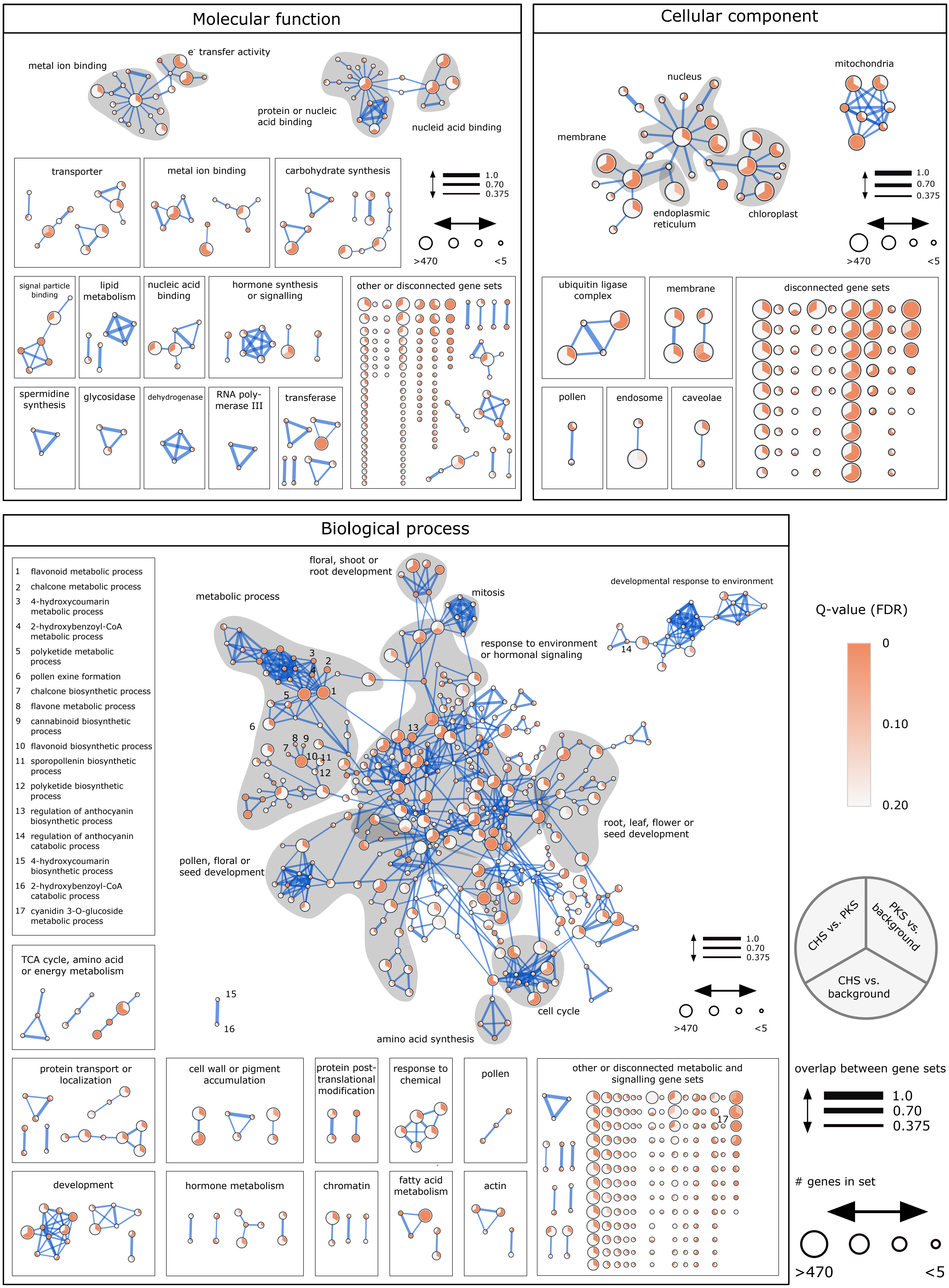

Molecular function
metal ion binding
e- transfer activity
protein or nucleic acid binding
nucleid acid binding
transporter
metal ion binding
carbohydrate synthesis
1.0
0.70
0.375
>470
<5
signal particle binding
lipid metabolism
nucleic acid binding
hormone synthesis or signalling
other or disconnected gene sets
spermidine synthesis
glycosidase
dehydrogenase
RNA poly-merase III
transferase
Cellular component
nucleus
mitochondria
membrane
endoplasmic reticulum
chloroplast
1.0
0.70
0.375
>470
<5
ubiquitin ligase complex
membrane
disconnected gene sets
pollen
endosome
caveolae
Biological process
1 flavonoid metabolic process
2 chalcone metabolic process
3 4-hydroxycoumarin metabolic process
4 2-hydroxybenzoyl-CoA metabolic process
5 polyketide metabolic process
6 pollen exine formation
7 chalcone biosynthetic process
8 flavone metabolic process
9 cannabinoid biosynthetic process
10 flavonoid biosynthetic process
11 sporopollenin biosynthetic process
12 polyketide biosynthetic process
13 regulation of anthocyanin biosynthetic process
14 regulation of anthocyanin catabolic process
15 4-hydroxycoumarin biosynthetic process
16 2-hydroxybenzoyl-CoA catabolic process
17 cyanidin 3-O-glucoside metabolic process
floral, shoot or root development
metabolic process
mitosis
developmental response to environment
response to environment or hormonal signaling
root, leaf, flower or seed development
pollen, floral or seed development
cell cycle
amino acid synthesis
1.0
0.70
0.375
>470
<5
TCA cycle, amino acid or energy metabolism
protein transport or localization
cell wall or pigment accumulation
protein post-translational modification
response to chemical
pollen
other or disconnected metabolic and signalling gene sets
development
hormone metabolism
chromatin
fatty acid metabolism
actin
Q-value (FDR)
0
0.10
0.20
CHS vs. PKS
PKS vs. background
CHS vs. background
overlap between gene sets
1.0
0.70
0.375
### genes in set
>470
<5

Supplementary Figure 6: Gene Ontology enrichment of syntenic regions where PKS genes were included. Three enrichment sets were compared: A) Genes of PKS-containing syntenic regions were checked for enrichment against background (all genes in syntenic regions), B) Genes of syntenic regions in CHS-enriched clusters 2, 4, 5 and 14 against background, C) Genes of syntenic regions in CHS-enriched clusters 2, 4, 5 and 14 against syntenic genes of PKS-containing syntenic regions of all species. The syntenic regions were enriched to a total of 517 (biological process), 230 (molecular function) and 105 (cellular compartment) significant terms (FDR-corrected q-value < 0.05). Enriched terms were categorized into higher categories. Many enriched terms in the category 'Biological process' can be linked to flavonoid-related processes ('leaf, root, pollen, floral or seed development', 'response to environment and hormonal signaling'). The size of the vertex corresponds to the number of gene with the same GO term. Terms with a FDR-corrected q-value of <0.2 are displayed. Edges correspond to the similarity between terms based on their gene set overlap (50% Jaccard similarity and 50% overlap between terms with a cutoff of 0.375).

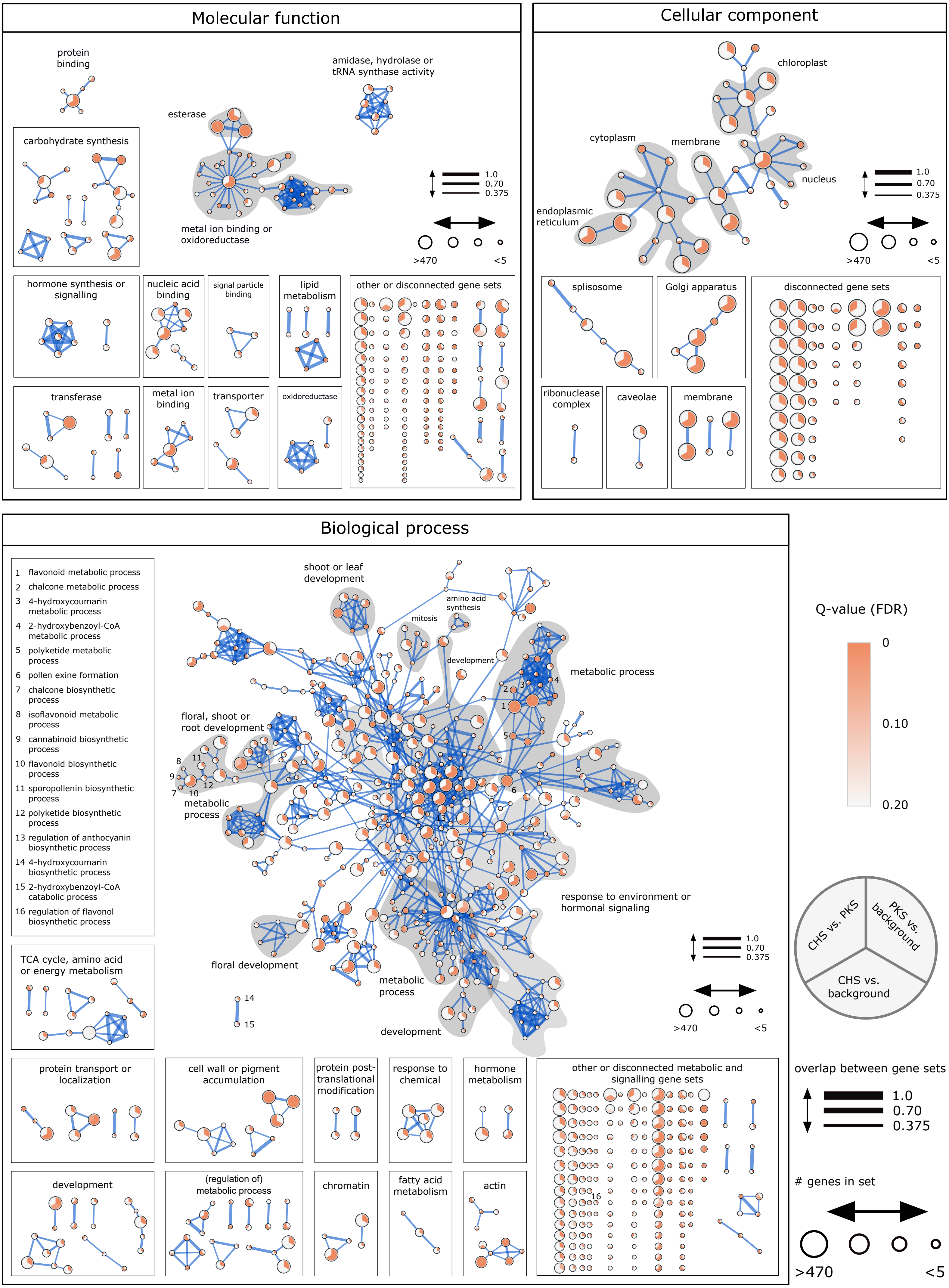

Molecular function
protein binding
amidase, hydrolase or tRNA synthase activity
esterase
carbohydrate synthesis
metal ion binding or oxidoreductase
1.0
0.70
0.375
>470
<5
hormone synthesis or signalling
nucleic acid binding
signal particle binding
lipid metabolism
other or disconnected gene sets
transferase
metal ion binding
transporter
oxidoreductase
Cellular component
chloroplast
cytoplasm
membrane
nucleus
endoplasmic reticulum
1.0
0.70
0.375
>470
<5
splisosome
Golgi apparatus
disconnected gene sets
ribonuclease complex
caveolae
membrane
Biological process
1 flavonoid metabolic process
2 chalcone metabolic process
3 4-hydroxycoumarin metabolic process
4 2-hydroxybenzoyl-CoA metabolic process
5 polyketide metabolic process
6 pollen exine formation
7 chalcone biosynthetic process
8 isoflavonoid metabolic process
9 cannabinoid biosynthetic process
10 flavonoid biosynthetic process
11 sporopollenin biosynthetic process
12 polyketide biosynthetic process
13 regulation of anthocyanin biosynthetic process
14 4-hydroxycoumarin biosynthetic process
15 2-hydroxybenzoyl-CoA catabolic process
16 regulation of flavonol biosynthetic process
shoot or leaf development
amino acid synthesis
mitosis
development
metabolic process
floral, shoot or root development
metabolic process
response to environment or hormonal signaling
floral development
metabolic process
development
1.0
0.70
0.375
>470
<5
TCA cycle, amino acid or energy metabolism
protein transport or localization
cell wall or pigment accumulation
protein post-translational modification
response to chemical
hormone metabolism
other or disconnected metabolic and signalling gene sets
development
(regulation of) metabolic process
chromatin
fatty acid metabolism
actin
Q-value (FDR)
0
0.10
0.20
CHS vs. PKS
PKS vs. background
CHS vs. background
overlap between gene sets
1.0
0.70
0.375
### genes in set
>470
<5

Supplementary Figure 7: Gene Ontology enrichment of syntenic regions where PKS genes were included. Three enrichment sets were compared: A) Genes of PKS-containing syntenic regions were checked for enrichment against background (all genes in data set), B) Genes of syntenic regions in CHS-enriched clusters 2, 3, 5 and 14 against background, C) Genes of syntenic regions in CHS-enriched clusters 2, 3, 5 and 14 against syntenic genes of PKS-containing syntenic regions of all species. The syntenic regions were enriched to a total of 626 (biological process), 232 (molecular function) and 109 (cellular compartment) significant terms (FDR-corrected q-value < 0.05). Enriched terms were categorized into higher categories. Many enriched terms in the category 'Biological process' can be linked to flavonoid-related processes ('leaf, root, pollen, floral or seed development', 'response to environment or hormonal signaling'). The size of the vertex corresponds to the number of gene with the same GO term. Terms with a FDR-corrected q-value of <0.2 are displayed. Edges correspond to the similarity between terms based on their gene set overlap (50% Jaccard similarity and 50% overlap between terms with a cutoff of 0.375).

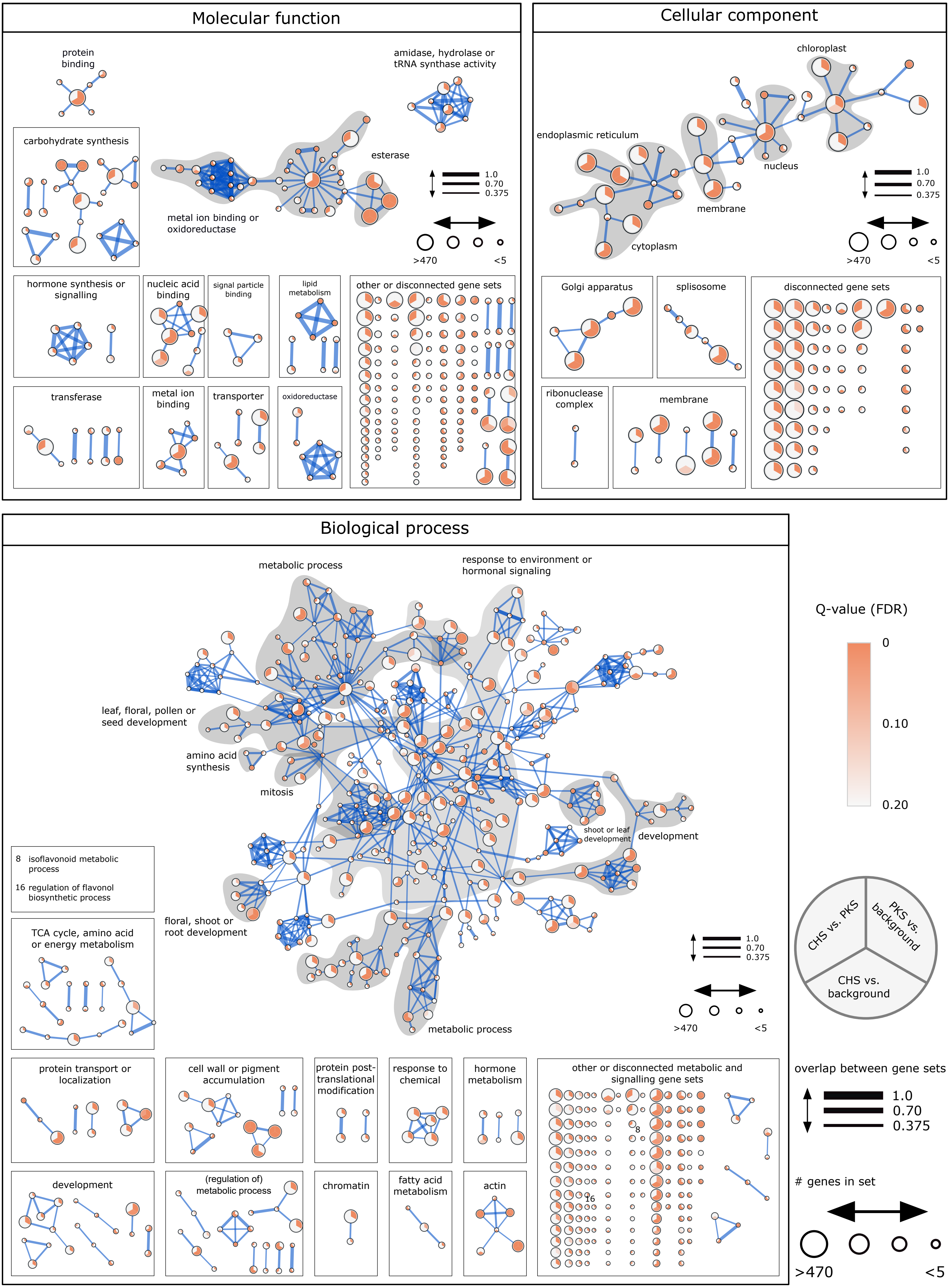
Molecular function
protein binding
amidase, hydrolase or tRNA synthase activity
carbohydrate synthesis
esterase
metal ion binding or oxidoreductase
1.0
0.70
0.375
>470
<5
hormone synthesis or signalling
nucleic acid binding
signal particle binding
lipid metabolism
other or disconnected gene sets
transferase
metal ion binding
transporter
oxidoreductase
Cellular component
chloroplast
endoplasmic reticulum
nucleus
membrane
cytoplasm
Golgi apparatus
splisosome
disconnected gene sets
ribonuclease complex
membrane
Biological process
metabolic process
response to environment or hormonal signaling
leaf, floral, pollen or seed development
amino acid synthesis
mitosis
shoot or leaf development
development
8 isoflavonoid metabolic process
16 regulation of flavonol biosynthetic process
TCA cycle, amino acid or energy metabolism
floral, shoot or root development
metabolic process
Q-value (FDR)
0
0.10
0.20
CHS vs. PKS
PKS vs. background
CHS vs. background
protein transport or localization
cell wall or pigment accumulation
protein post-translational modification
response to chemical
hormone metabolism
other or disconnected metabolic and signalling gene sets
overlap between gene sets
### genes in set
development
(regulation of) metabolic process
chromatin
fatty acid metabolism
actin

Supplementary Figure 8: Gene Ontology enrichment of syntenic regions where *PKS* genes were excluded. *PKS* genes were removed prior to conducting enrichment analysis. Three enrichment sets were compared: A) Genes of *PKS*-containing syntenic regions were checked for enrichment against background (all genes in data set), B) Genes of syntenic regions in *CHS*-enriched clusters 2, 3, 5 and 14 against background, C) Genes of syntenic regions in *CHS*-enriched clusters 2, 3, 5 and 14 against syntenic genes of PKS-containing syntenic regions of all species. The syntenic regions were enriched to a total of 553 (biological process), 224 (molecular function) and 107 (cellular compartment) significant terms (FDR-corrected q-value < 0.05). Enriched terms were categorized into higher categories. Many enriched terms in the category 'Biological process' can be linked to flavonoid-related processes ('leaf, root, pollen, floral or seed development', 'response to environment or hormonal signaling'). The size of the vertex corresponds to the number of gene with the same GO term. Terms with a FDR-corrected q-value of <0.2 are displayed. Edges correspond to the similarity between terms based on their gene set overlap (50% Jaccard similarity and 50% overlap between terms with a cutoff of 0.375).

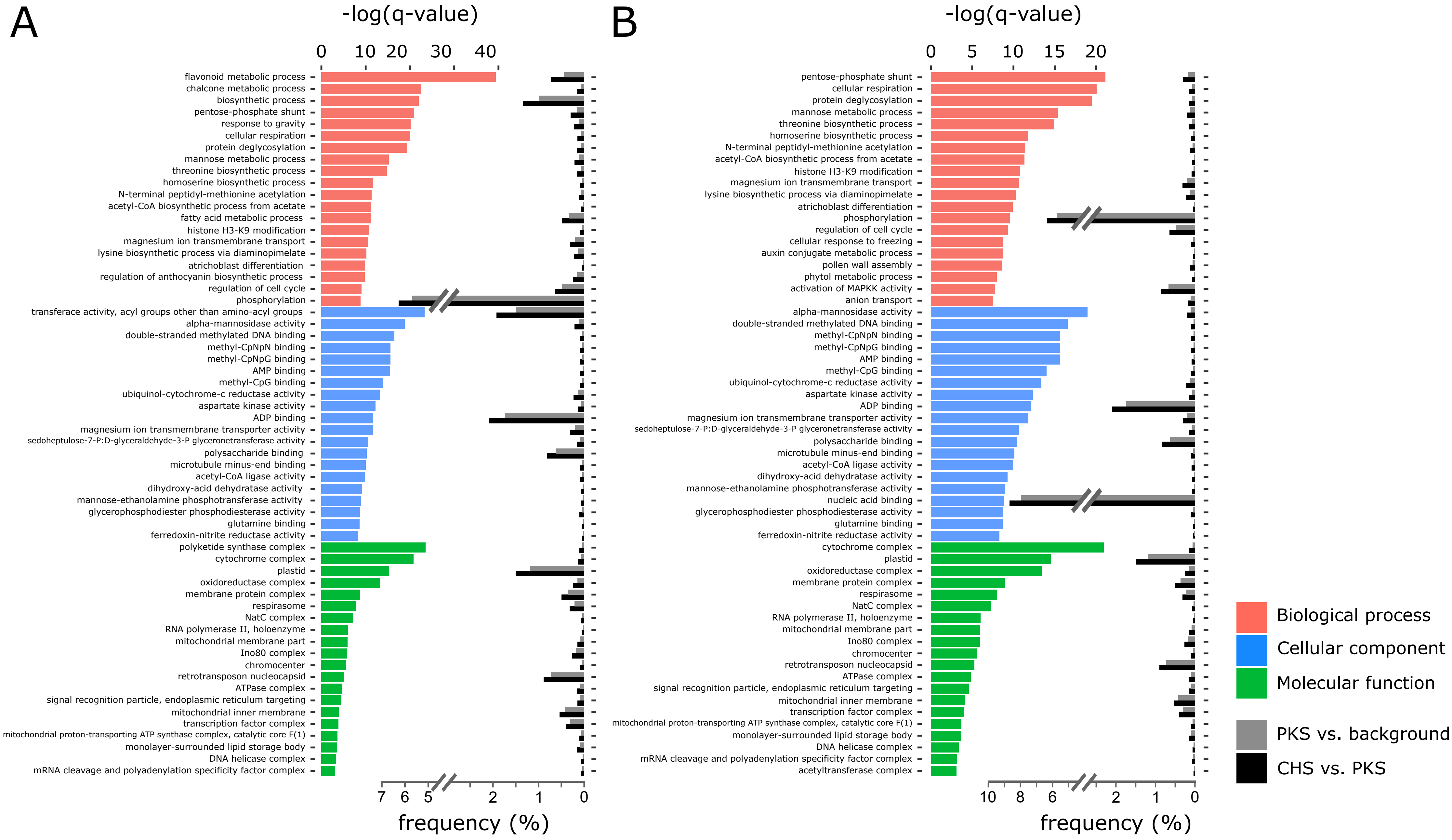
A
B
-log(q-value)
0 10 20 30 40
0 5 10 15 20
flavonoid metabolic process
chalcone metabolic process
biosynthetic process
pentose-phosphate shunt
response to gravity
cellular respiration
protein deglycosylation
mannose metabolic process
threonine biosynthetic process
homoserine biosynthetic process
N-terminal peptidyl-methionine acetylation
acetyl-CoA biosynthetic process from acetate
fatty acid metabolic process
histone H3-K9 modification
magnesium ion transmembrane transport
lysine biosynthetic process via diaminopimelate
atrichoblast differentiation
regulation of anthocyanin biosynthetic process
regulation of cell cycle
phosphorylation
transferace activity, acyl groups other than amino-acyl groups
alpha-mannosidase activity
double-stranded methylated DNA binding
methyl-CpNpN binding
methyl-CpNpG binding
AMP binding
methyl-CpG binding
ubiquinol-cytochrome-c reductase activity
aspartate kinase activity
ADP binding
magnesium ion transmembrane transporter activity
sedoheptulose-7-P:D-glyceraldehyde-3-P glyceronetransferase activity
polysaccharide binding
microtubule minus-end binding
acetyl-CoA ligase activity
dihydroxy-acid dehydratase activity
mannose-ethanolamine phosphotransferase activity
glycerophosphodiester phosphodiesterase activity
glutamine binding
ferredoxin-nitrite reductase activity
polyketide synthase complex
cytochrome complex
plastid
oxidoreductase complex
membrane protein complex
respirasome
NatC complex
RNA polymerase II, holoenzyme
mitochondrial membrane part
Ino80 complex
chromocenter
retrotransposon nucleocapsid
ATPase complex
signal recognition particle, endoplasmic reticulum targeting
mitochondrial inner membrane
transcription factor complex
mitochondrial proton-transporting ATP synthase complex, catalytic core F(1)
monolayer-surrounded lipid storage body
DNA helicase complex
mRNA cleavage and polyadenylation specificity factor complex
7 6 5 2 1 0
frequency (%)
pentose-phosphate shunt
cellular respiration
protein deglycosylation
mannose metabolic process
threonine biosynthetic process
homoserine biosynthetic process
N-terminal peptidyl-methionine acetylation
acetyl-CoA biosynthetic process from acetate
histone H3-K9 modification
magnesium ion transmembrane transport
lysine biosynthetic process via diaminopimelate
atrichoblast differentiation
phosphorylation
regulation of cell cycle
cellular response to freezing
auxin conjugate metabolic process
pollen wall assembly
phytol metabolic process
activation of MAPKK activity
anion transport
alpha-mannosidase activity
double-stranded methylated DNA binding
methyl-CpNpN binding
methyl-CpNpG binding
AMP binding
methyl-CpG binding
ubiquinol-cytochrome-c reductase activity
aspartate kinase activity
ADP binding
magnesium ion transmembrane transporter activity
sedoheptulose-7-P:D-glyceraldehyde-3-P glyceronetransferase activity
polysaccharide binding
microtubule minus-end binding
acetyl-CoA ligase activity
dihydroxy-acid dehydratase activity
mannose-ethanolamine phosphotransferase activity
nucleic acid binding
glycerophosphodiester phosphodiesterase activity
glutamine binding
ferredoxin-nitrite reductase activity
cytochrome complex
plastid
oxidoreductase complex
membrane protein complex
respirasome
NatC complex
RNA polymerase II, holoenzyme
mitochondrial membrane part
Ino80 complex
chromocenter
retrotransposon nucleocapsid
ATPase complex
signal recognition particle, endoplasmic reticulum targeting
mitochondrial inner membrane
transcription factor complex
mitochondrial proton-transporting ATP synthase complex, catalytic core F(1)
monolayer-surrounded lipid storage body
DNA helicase complex
mRNA cleavage and polyadenylation specificity factor complex
acetyltransferase complex
10 8 6 2 1 0
frequency (%)
Biological process
Cellular component
Molecular function
PKS vs. background
CHS vs. PKS

Supplementary Figure 9: Enriched terms for syntenic genes of CHS-enriched clusters vs. syntenic genes of PKS-containing syntenic regions of all species. Top 20 enriched terms with lowest q-value for 'Biological process', 'Molecular function' and 'Cellular compartment' are shown together with their frequencies in the data sets. A: Data set including *PKS* genes. In total 153 terms for 'Biological process', 84 terms for 'Molecular function' and 34 terms 'Cellular compartment'. B: Data set where *PKS* genes were excluded prior to the analysis. In total 144 terms for 'Biological process', 83 terms for 'Molecular function' and 36 terms for 'Cellular compartment'. Enrichment based on one-sided Fisher test with FDR correction.

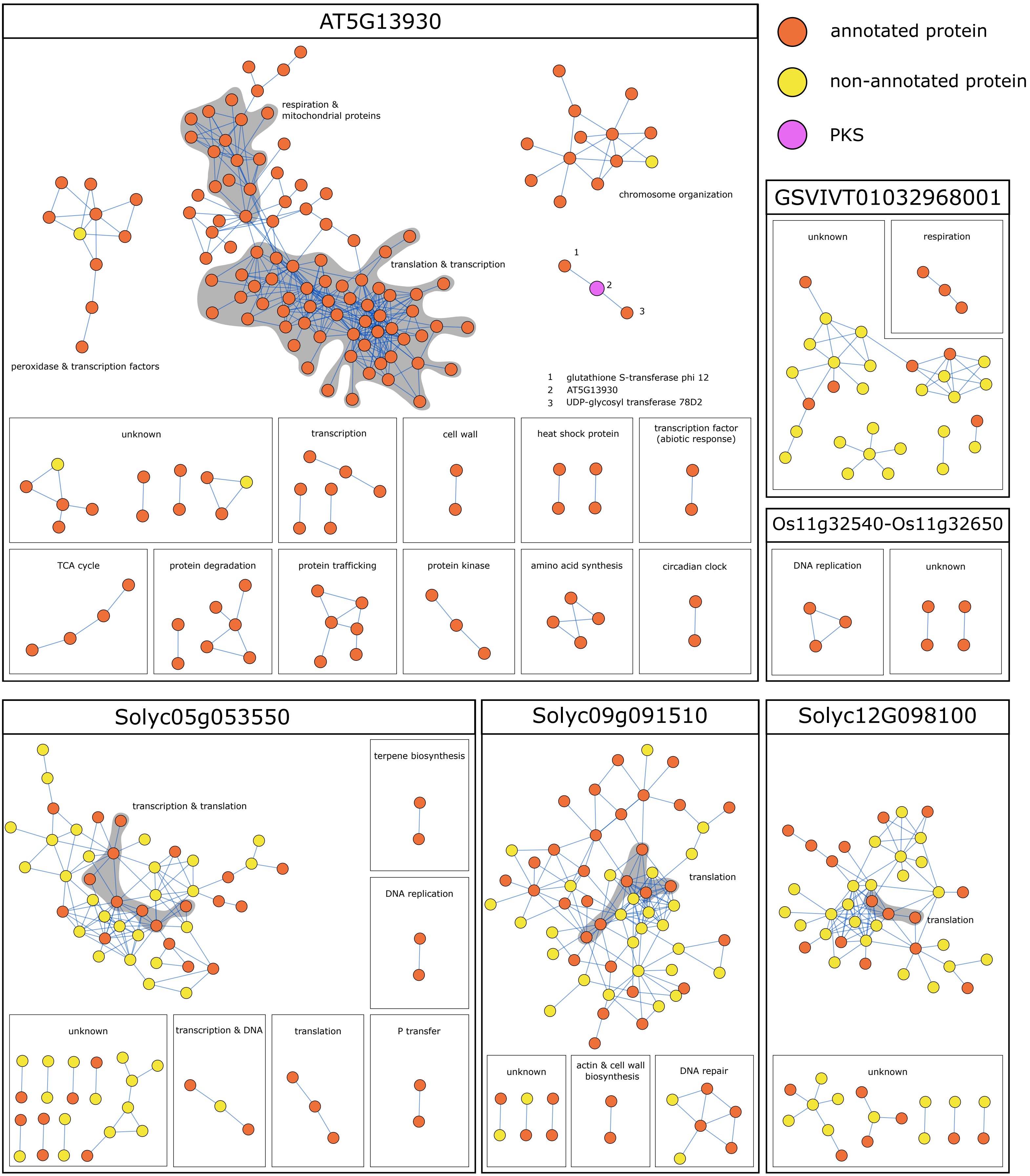

AT5G13930
annotated protein
non-annotated protein
PKS
respiration &
mitochondrial proteins
chromosome organization
translation & transcription
peroxidase & transcription factors
1 glutathione S-transferase phi 12
2 AT5G13930
3 UDP-glycosyl transferase 78D2
unknown
transcription
cell wall
heat shock protein
transcription factor
(abiotic response)
TCA cycle
protein degradation
protein trafficking
protein kinase
amino acid synthesis
circadian clock
GSVIVT01032968001
unknown
respiration
Os11g32540-Os11g32650
DNA replication
unknown
Solyc05g053550
transcription & translation
terpene biosynthesis
DNA replication
unknown
transcription & DNA
translation
P transfer
Solyc09g091510
translation
unknown
actin & cell wall
biosynthesis
DNA repair
Solyc12G098100
translation
unknown

Supplementary Figure 10: Co-expression network of syntenic regions in *Arabidopsis thaliana*, *Solanum lycopersicum*, *Vitis vinifera* and *Zea mays*. Co-expression and associated annotation was inferred from StringDB (meaning of network edges: confidence; minimum required interaction score: medium, 0.400). Displayed are PKS from the ‘R-4-C’ type from the above species in the cluster 2, 4 and 14. Region Os11g32540-Os11g32650 contains the *PKS* genes: LOC_Os11g32540, LOC_Os11g32550, LOC_Os11g32580, LOC_Os11g32610, LOC_Os11g32620 and LOC_Os11g32650. The co-expression of the syntenic region containing the tandem Zm00001d007400 and Zm00001d007403 and the region containing the tandem PKS genes Zm00001d052673, Zm00001d052675 and Zm00001d052676 from *Zea mays* (no co-expression between the PKS genes and other genes in the syntenic region) is not displayed. The analysis was done for all PKS-containing regions from *A. thaliana*, *S. lycopersium*, *O. sativa*, *V. vinifera*, *Z. mays.* The analysis revealed that there is no co-expression of *PKS* genes with other genes in PKS-containing syntenic regions.

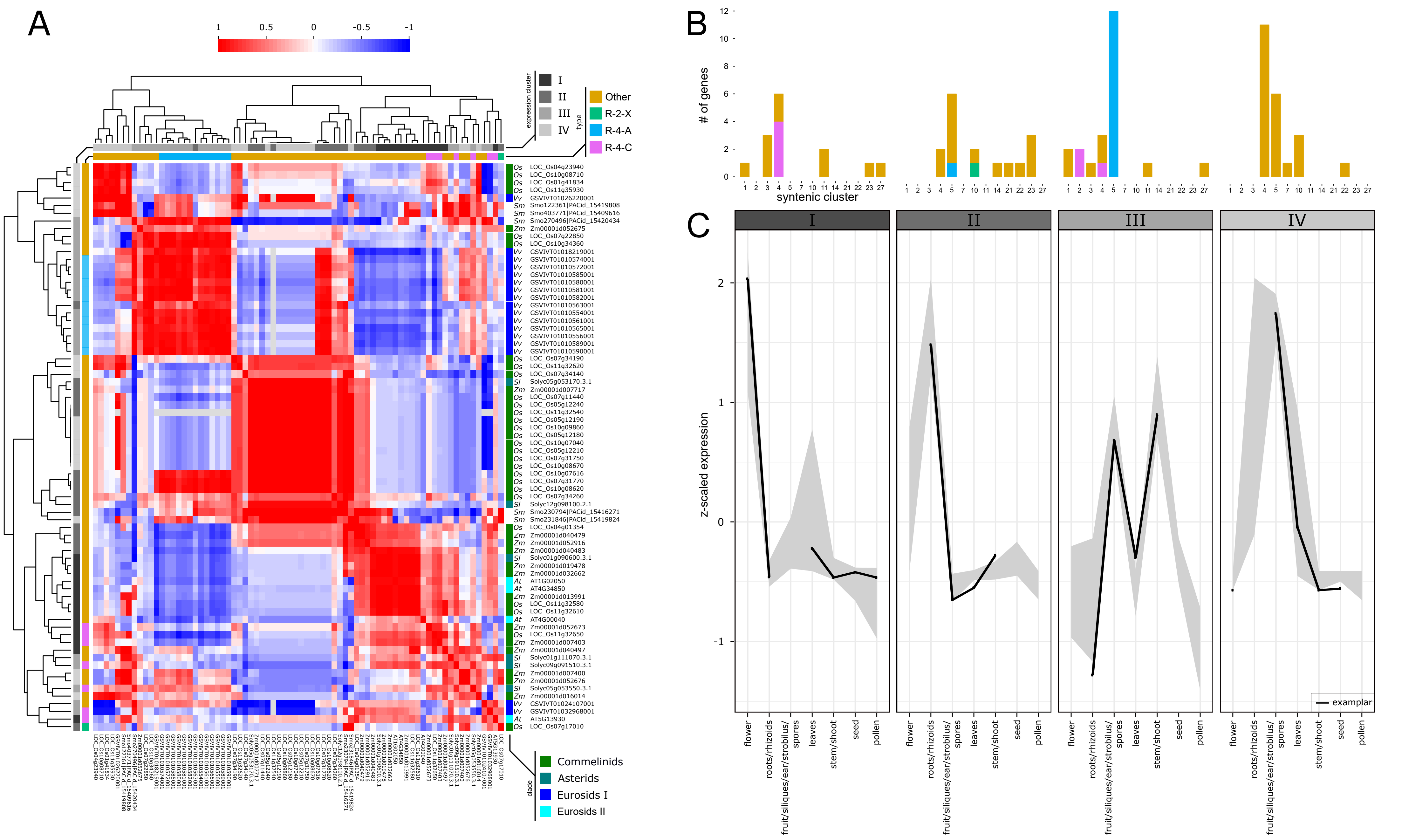

A
1
0.5
0
-0.5
-1
expression cluster
I
II
III
IV
type
Other
R-2-X
R-4-A
R-4-C
Os LOC_Os04g23940
Os LOC_Os10g08710
Os LOC_Os01g41834
Os LOC_Os11g35930
Vv GSVIVT01026220001
Sm Smo122361|PACid_15419808
Sm Smo403771|PACid_15409616
Sm Smo270496|PACid_15420434
Zm Zm00001d052675
Os LOC_Os07g22850
Os LOC_Os10g34360
Vv GSVIVT01018219001
Vv GSVIVT01010574001
Vv GSVIVT01010572001
Vv GSVIVT01010585001
Vv GSVIVT01010580001
Vv GSVIVT01010581001
Vv GSVIVT01010582001
Vv GSVIVT01010563001
Vv GSVIVT01010554001
Vv GSVIVT01010561001
Vv GSVIVT01010565001
Vv GSVIVT01010556001
Vv GSVIVT01010589001
Vv GSVIVT01010590001
Os LOC_Os07g34190
Os LOC_Os11g32620
Os LOC_Os07g34140
Sl Solyc05g053170.3.1
Zm Zm00001d007717
Os LOC_Os07g11440
Os LOC_Os05g12240
Os LOC_Os11g32540
Os LOC_Os05g12190
Os LOC_Os10g09860
Os LOC_Os05g12180
Os LOC_Os10g07040
Os LOC_Os05g12210
Os LOC_Os07g31750
Os LOC_Os10g08670
Os LOC_Os10g07616
Os LOC_Os07g31770
Os LOC_Os10g08620
Os LOC_Os07g34260
Sl Solyc12g098100.2.1
Sm Smo230794|PACid_15416271
Sm Smo231846|PACid_15419824
Os LOC_Os04g01354
Zm Zm00001d040479
Zm Zm00001d052916
Zm Zm00001d040483
Sl Solyc01g090600.3.1
Zm Zm00001d019478
Zm Zm00001d032662
At AT1G02050
At AT4G34850
Zm Zm00001d013991
Os LOC_Os11g32580
Os LOC_Os11g32610
At AT4G00040
Zm Zm00001d052673
Os LOC_Os11g32650
Zm Zm00001d007403
Zm Zm00001d040497
Sl Solyc01g111070.3.1
Sl Solyc09g091510.3.1
Zm Zm00001d007400
Zm Zm00001d052676
Sl Solyc05g053550.3.1
Zm Zm00001d016014
Vv GSVIVT01024107001
Vv GSVIVT01032968001
At AT5G13930
Os LOC_Os07g17010
clade
Commelinids
Asterids
Eurosids I
Eurosids II
B
### of genes
syntenic cluster
C
I
II
III
IV
z-scaled expression
flower
roots/rhizoids
fruit/siliques/ear/strobilus/spores
leaves
stem/shoot
seed
pollen
examplar

Supplementary Figure 11: Gene expression of *PKS* genes of *Arabidopsis thaliana*, *Selaginella moellendorffii*, *Solanum lycopersicum*, *Vitis vinifera* and *Zea mays*. A: Expression values were taken from the CoNekT database and averaged for the respective tissue. Pearson correlation scores were calculated and clusters defined by affinity propagation clustering (clusters I-IV). Heatmap shows Pearson correlation scores between the different transcripts. B: Network syntenic cluster membership for expression clusters I-IV as defined in A. The color values correspond to the PKS classification by pPAP. C: Expression profiles of clusters I-IV as defined in A. Displayed are the 25% and 75% quantile of expression values together with the exemplar given by affinity propagation clustering. The analysis showed that the syntenic cluster membership does not dictate the expression pattern of the PKS gene, suggesting that the evolution of gene expression is uncoupled from the cluster membership.

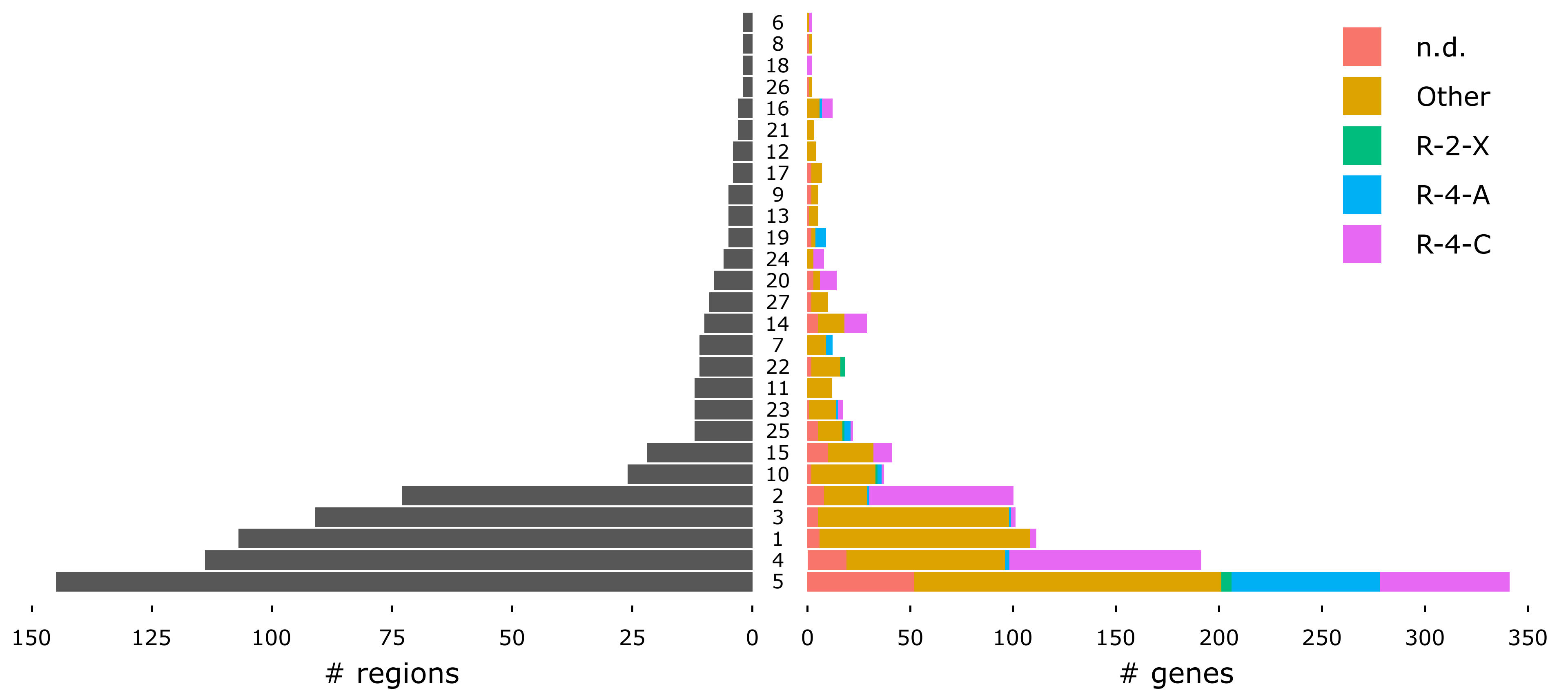

n.d.
Other
R-2-X
R-4-A
R-4-C
6
8
18
26
16
21
12
17
9
13
19
24
20
27
14
7
22
11
23
25
15
10
2
3
1
4
5
150
125
100
75
50
25
0
0
50
100
150
200
250
300
350
### regions
### genes

Supplementary Figure 12: Number of regions and genes per cluster in the syntenic network. The clusters 2, 4, 5 and 14 are enriched for the ‘R-4-C’ function referring to chalcone synthase. The clusters 1, 3, 11 and 27 contain ‘Other’-type *PKS* genes corresponding to LAP5 and LAP6 function. For the pPAP classification of PKS genes refer to the legend.

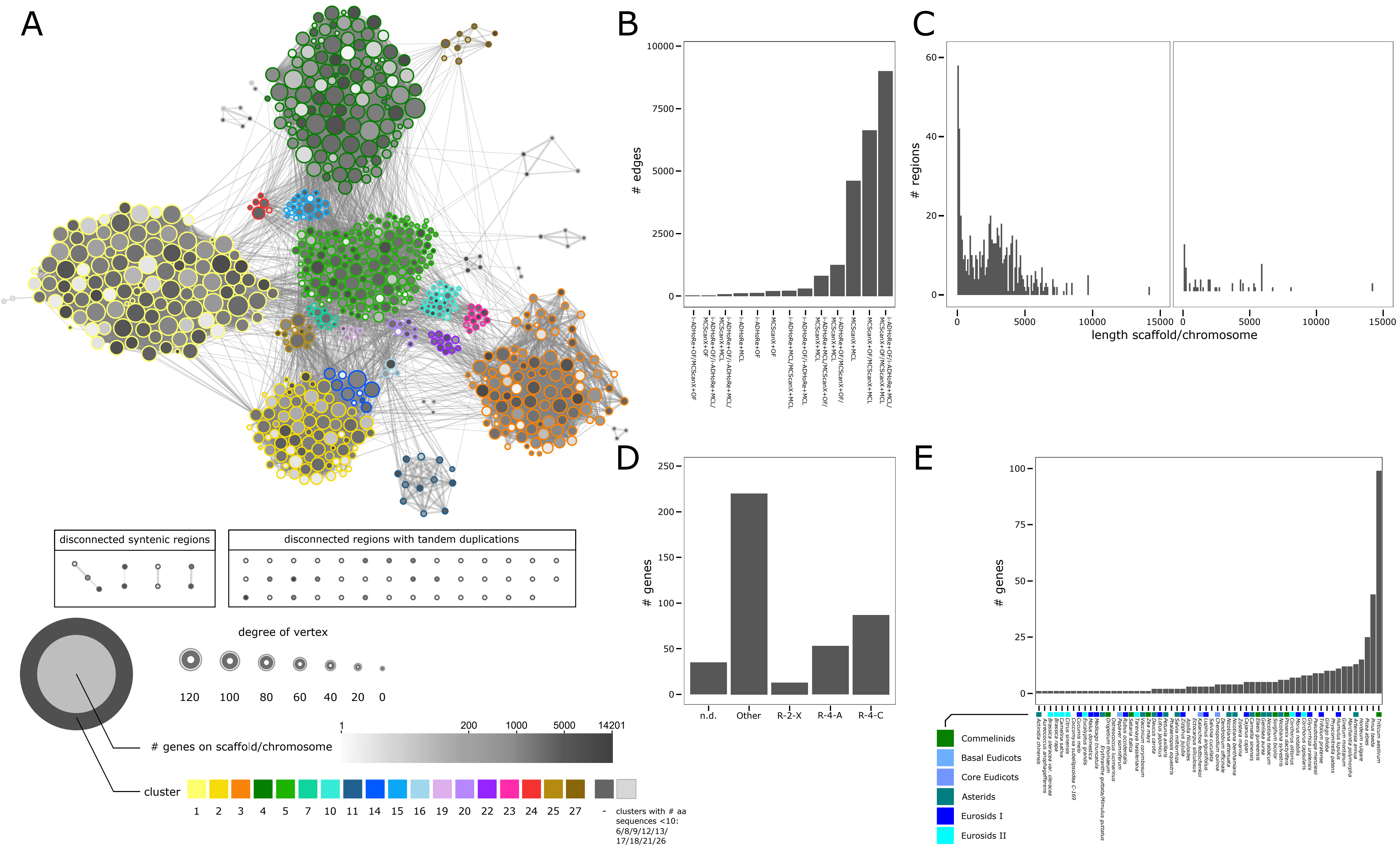

A
disconnected syntenic regions
disconnected regions with tandem duplications
degree of vertex
120
100
80
60
40
20
0
### genes on scaffold/chromosome
1
200
1000
5000
14201
cluster
1
2
3
4
5
7
10
11
14
15
16
19
20
22
23
24
25
27
-
clusters with # aa sequences <10: 6/8/9/12/13/17/18/21/26
B
### edges
10000
7500
5000
2500
0
i-ADHoRe+OF/MCScanX+OF
i-ADHoRe+OF/i-ADHoRe+MCL/MCScanX+OF
i-ADHoRe+OF/i-ADHoRe+MCL/MCScanX+MCL
i-ADHoRe+MCL
i-ADHoRe+OF
MCScanX+OF
i-ADHoRe+MCL/MCScanX+MCL
i-ADHoRe+OF/i-ADHoRe+MCL
i-ADHoRe+MCL/MCScanX+OF/MCScanX+MCL
i-ADHoRe+OF/MCScanX+OF/MCScanX+MCL
MCScanX+MCL
MCScanX+OF/MCScanX+MCL
i-ADHoRe+OF/i-ADHoRe+MCL/MCScanX+OF/MCScanX+MCL
C
### regions
60
40
20
0
0
5000
10000
15000
length scaffold/chromosome
D
### genes
250
200
150
100
50
0
n.d.
Other
R-2-X
R-4-A
R-4-C
E
### genes
100
75
50
25
0
Commelinids
Basal Eudicots
Core Eudicots
Asterids
Eurosids I
Eurosids II

Supplementary Figure 13: Quality of synteny network. A: Synteny network as shown in Fig. 2 with information on number of genes of scaffold/chromosome. B: Barplot giving information on which method the edges within network are supported. By highest number, edges are supported by all four methods, the second highest number of edges is supported by MCScanX+OrthoFinder and MCScanX+MCL. OF: OrthoFinder. C: Barplot showing how many regions have a certain length of scaffold/chromosome, right: singletons (nodes that do not connect to other nodes and do not contain tandem duplications), left: network, disconnected syntenic regions and disconnected regions with tandem duplications. D: Barplot giving information on how many genes are per type of pPAP-classified PKS for singletons. E: Barplot giving information on how many singleton regions exist per species.

### Supplementary Tables

Supplementary Table S1: Number of type III polyketide synthases for selected species. Shown are clades with species that have 15 or more copies of type III PKS sequences with length ≥ 200 amino acids together with all species from the same order (shown in brackets):

| **clade** | **species** | **# PKS (Orthofinder/MCL)** | # PKS ≥ 200 amino acids (Orthofinder/MCL) |
|---|---|---|---|
| Asterales | *Artemisia annua* | 16/15 | 16/15 |
| Ericales | *Camellia sinensis* | 18/18 | 17/17 |
| Fabales | *Arachis duranensis* | 33/33 | 33/33 |
| | *Arachis ipaensis* | 26/25 | 25/25 |
| | *(Cajanus cajan)* | 15/15 | 12/12 |
| | *Glycine max* | 24/24 | 21/21 |
| | *Glycyrrhiza uralensis* | 19/19 | 17/17 |
| | *Lotus japonicus* | 35/35 | 16/16 |
| | *Medicago truncatula* | 25/25 | 25/25 |
| | *Phaseolus vulgaris* | 21/21 | 13/13 |
| | *Trifolium pratense* | 23/24 | 16/17 |
| Funariales | *Physcomitrella patens* | 30/38 | 26/28 |
| Ginkgoales | *Ginkgo biloba* | 16/16 | 15/15 |
| Gnetales | (*Gnetum montanum*) | 20/22 | 11/11 |
| Marchantiales | *Marchantia polymorpha* | 25/26 | 24/24 |
| Pinales | (*Picea abies*) | 27/27 | 14/14 |
| | *Pinus taeda* | 42/44 | 42/44 |
| | *Pseudotsuga menziesii* | 18/18 | 15/15 |
| Poales,<br>BEP clade of Poaceae | (*Brachypodium distachyon*) | 10/11 | 10/10 |
| | *Hordeum vulgare* | 23/24 | 20/20 |
| | *Leersia perrieri* | 21/21 | 20/20 |
| | *Oryza rufipogon* | 27/30 | 23/23 |
| | *Oryza sativa* | 30/35 | 28/29 |
| | *Triticum aestivum* | 82/99 | 50/54 |
| Poales,<br>PACMAD clade of Poaceae | *Setaria italica* | 26/27 | 20/21 |
| | *Sorghum bicolor* | 31/37 | 30/36 |
| | *Zea mays* | 18/18 | 16/16 |
| Sapindales | *Citrus clementina* | 15/15 | 15/15 |
| | *Citrus sinensis* | 16/17 | 14/15 |
| Vitales | *Vitis vinifera* | 26/26 | 17/17 |
| Zingiberales | *Musa acuminata* | 21/21 | 19/19 |

Supplementary Table S2: Species not represented in the synteny network due to missing type III *PKS* genes, taxonomic distance and/or low quality genome assemblies. The obtained synteny network (Fig. 2) contained syntenic regions of 105 of the initial 126 species; nine species did not contain type III PKS sequences, six species did not show synteny/tandem duplications and six showed only tandem duplications and no synteny due to lower quality genome assemblies.

| | species | division | class |
|---|---|---|---|
| no type III PKS sequences | *Chara braunii* | Charophyta | Charophyceae |
| | *Chlamydomonas reinhardtii* | Chlorophyta | Chlorophyceae |
| | *Cyanidioschyzon merolae* | Rhodophyta | Cyanidiophyceae |
| | *Cyanophora paradoxa* | Glaucophyta | Glaucophyceae |
| | *Dunaliella salina* | Chlorophyta | Chlorophyceae |
| | *Klebsormidium nitens* | Charophyta | Klebsormidiophyceae |
| | *Porphyridium purpureum* | Rhodophyta | Porphyridiophyceae |
| | *Synechocystis sp.* PCC 6803 | Cyanophyta | Cyanophyceae |
| | *Volvox carteri* | Chlorophyta | Chlorophyceae |
| no synteny/tandem duplications | *Aureococcus aneophagefferens* | Ochrophyta | Pelagophyceae |
| | *Ectocarpus siliculosus* | Ochrophyta | Phaeophyceae |
| | *Coccomyxa sp.* C169 | Chlorophyta | Trebouxiophyceae |
| | *Genlisea aurea* | Magnoliophyta | Magnoliopsida |
| | *Pinus taeda* | Pinophyta | Pinopsida |
| | *Ostreococcus lucimarinus* | Chlorophyta | Mamiellophyceae |
| only tandem duplications | *Ginkgo biloba* | Ginkgophyta | Ginkgoopsida |
| | *Gnetum montanum* | Gnetophyta | Gnetopsida |
| | *Humulus lupulus* | Magnoliophyta | Magnoliopsida |
| | *Picea abies* | Pinophyta | Pinopsida |
| | *Pseudotsuga menziessii* | Pinophyta | Pinopsida |
| | *Triticum aestivum* | Magnoliophyta | Liliopsida |

Supplementary Table S3: Number of PKS genes in the network for species belonging to the Asterids. Syntenic cluster 4 does not contain PKS-containing genomic regions of species belonging to Asterids (except *Daucus carota* and *Vaccinium corymbosum*). The table shows that for most species of the Asterids, the assembly quality is large (low number of non-connecting genes for most of the species), indicating that the depletion of Asterids in syntenic cluster 4 is most likely not an artefact of low assembly quality:

| species | # of PKS | # of genes in network | # of non-connecting single genes | # of non-connecting tandem-duplicated genes |
|---|---|---|---|---|
| *Actinidia chinensis* | 10 | 9 | 1 | 0 |
| *Artemisia annua* | 16 | 1 | 15 | 2 |
| *Camellia sinensis* | 18 | 5 | 5 | 8 |
| *Capsicum annuum glabriusculum* | 10 | 10 | - | - |
| *Coffea arabica* | 7 | 7 | - | - |
| *Daucus carota* | 14 | 12 | 2 | - |
| *Erythranthe guttata/ Mimulus guttatus* | 8 | 7 | 1 | - |
| *Helianthus annuus* | 12 | 12 | - | - |
| *Ipomoea nil* | 7 | 7 | - | - |
| *Petunia axillaris* | 16 | 8 | 3 | 5 |
| *Petunia inflata* | 9 | 8 | 1 | - |
| *Nicotiana attenuata* | 7 | 3 | 4 | - |
| *Nicotiana benthamiana* | 14 | 10 | 4 | - |
| *Nicotiana sylvestris* | 7 | 1 | 6 | - |
| *Nicotiana tabacum* | 11 | 6 | 5 | - |
| *Olea europaea* | 7 | 7 | - | - |
| *Salvia miltiorrhiza* | 5 | 3 | 2 | |
| *Solanum lycopersicum* | 7 | 7 | - | - |
| *Solanum pennellii* | 8 | 8 | - | - |
| Solanum tuberosum | 12 | 12 | - | - |
| *Vaccinium corymbosum* | 9 | 8 | 1 | - |

Supplementary Table S4: Sequences in LAP5/LAP6 ortholog-specific clade. Given are representatives of Angiosperms sequences in the clade (sequences from *Arabidopsis thaliana*, *Solanum lycopersicum*, *Oryza sativa*, *Zea mays*) and all present gymnosperm and Bryophyta, Marchantiophyta and Pteridophyta. Amino acid sequences from gymnosperm, Bryophyta, Marchantiophyta and Pteridophyta were blasted against sequences of *A. thaliana* and query cover, identity and E-value was retrieved to the closest orthologue given by blastp:

| Gene | cluster | type | ortholog | query cover [%] | identity [%] | E-value |
|---|---|---|---|---|---|---|
| arath_AT1G02050 | 15 | Other | LAP6 [3,4] | | | |
| arath_AT4G00040 | 14 | Other | | | | |
| solyc_Solyc01g090600.3.1 | 13 | | *At*LAP6 [5] | | | |
| zeama_Zm00001d032662 | 18 | Other | *At*LAP6 and *Os*LAP6 [3] | | | |
| zeama_Zm00001d013991 | 18 | Other | *At*LAP6 and *Os*LAP6 [3] | | | |
| orysa_LOC_Os10g34360 | 18 | Other | CHSL1 (linked to immature panicle) [3] | | | |
| arath_AT4G34850 | 9 | Other | LAP5 [3,4] | | | |
| solyc_Solyc01g111070.3.1 | 9 | Other | *At*LAP5 [5] | | | |
| zeama_Zm00001d019478 | 19 | Other | *At*LAP5 and OsLAP5 [3] | | | |
| orysa_LOC_Os07g22850 | 19 | Other | CHSL2 (linked to immature panicle) [6] | | | |
| picab_MA_619670g0010 | - | nd | *At*LAP5 | 95.0 | 64.94 | 5e-67 |
| picab_MA_4931944g0010 | - | nd | *At*LAP6 | 95.0 | 86.05 | 8e-45 |
| picab_MA_1748261g0010 | - | nd | *At*LAP6 | 98.0 | 63.73 | 9e-38 |
| pseme_PSME_00004131-RA | - | Other | *At*LAP5 | 96.0 | 67.72 | 0.0 |
| pinpi_PPI00030756 | not incl. | Other | *At*LAP5 | 100 | 67.93 | 0.0 |
| ginbi_Gb_02579 | - | Other | *At*LAP5 | 96.0 | 71.84 | 0.0 |
| gnemo_TnS000356079t05 | - | Other | *At*LAP6 | 81 | 66.46 | 5e-152 |
| equgi_cds.Locus_29021_ Transcript_1_1_m.39955 | not incl. | Other | *At*LAP5 | 100 | 60.33 | 8e-161 |
| equgi_cds.Locus_25846_Transcript_2_9_m.38839/ equgi_cds.Locus_25846_Transcript_9_9_m.38842 | not incl. | nd/nd | *At*LAP5 | 94.0 | 60.24 | 2e-69 |
| sphfa_Sphfalx0085s0032.1 | 4 | Other | *At*LAP5 | 85.0 | 56.03 | 3e-153 |
| sphfa_Sphfalx0160s0011.1 | - | Other | *At*LAP5 | 90.0 | 58.75 | 7e-161 |
| marpo_Mapoly0014s0122.1 | - | Other | *At*LAP5 | 94.0 | 57.32 | 9e-162 |
| marpo_Mapoly0020s0082.1 | - | Other | *At*CHS=AT5G13930 | 82.0 | 46.42 | 3e-117 |
| phypa_Pp3c18_21820V3.1.p | - | nd | - | - | - | - |
| phypa_Pp3c2_32960V3.1.p | - | Other | *At*LAP5 | 90.0 | 58.12 | 3e-160 |
| selmo_Smo231846\|PACid_15419824 | 4 | Other | *At*CHS=AT5G13930 | 91.0 | 46.00 | 1e-65 |
| selmo_Smo122361\|PACid_15419808 | - | Other | *At*LAP5 | 95.0 | 59.06 | 1e-163 |

| | | | | | | |
|---|---|---|---|---|---|---|
| equgi_cds.Locus_10049_Transcript_1_1_m.20667 | not incl. | Other | *At*LAP6 | 94.0 | 47.18 | 2e-106 |
| equgi_cds.Locus_462_Transcript_3_4_m.1092 | not incl. | Other | *At*LAP5 | 83.0 | 44.5 | 4e-66 |
| equgi_cds.Locus_5106_Transcript_1_2_m.11120/ equgi_cds.Locus_5106_Transcript_2_2_m.11121 | not incl. | Other | *At*CHS=AT5G13930 | 73.0 | 50.00 | 9e-74 |
| equgi_cds.Locus_1312_Transcript_1_1_m.2882 | not incl. | Other | *At*CHS=AT5G13930 | 88.0 | 45.99 | 6e-123 |

Supplementary Table S5: GO enrichment analysis of syntenic regions against the background 'genes of syntenic regions'. Given are enriched terms for 'biological process' that are potentially linked to flavonoid metabolism. Enrichment tests were done for the following cases: genes in PKS syntenic regions against background (genes of syntenic regions, PKS-BG), genes in CHS-enriched syntenic regions against background (CHS-BG) and genes in CHS-enriched syntenic regions against syntenic genes in PKS syntenic regions (CHS-PKS). PKS genes were deleted from the syntenic regions prior to performing the enrichment analysis. p-values were adjusted for multiple testing by the Benjamini Hochberg method (q-values: . < 0.1, * ≤ 0.05, ** ≤ 0.01, *** ≤ 0.001, **** ≤ 0.0001). ABA: abscisic acid, JA: jasmonic acid, SA: salicylic acid, BR: brassinosteroids, GA: gibberellins.

| description | ID | PKS-BG | CHS-BG | CHS-PKS | clade specificity | reference |
|---|---|---|---|---|---|---|
| *direct and indirect effects of polyketides on biological processes* | | | | | | |
| actin cortical patch assembly | BP:0000147 | ** (< 0.007) | | | | [7] (human), [8] (in vitro), [9] (correlation) |
| actin cytoskeletal reorganization | BP:0031532 | (0.157) | | | | [7] (human), [8] (in vitro), [9] (correlation) |
| actin filament depolymerization | BP:0030042 | **** (< 6e-14) | | | | [7] (human), [8] (in vitro), [9] (correlation) |
| actin filament polymerization | BP:0030041 | **** (< 1e-18) | **** (< 8e-8) | | | [7] (human), [8] (in vitro), [9] (correlation) |
| anther development | BP:0048653 | **** (< 3e-6) | ** (<0.004) | | | [3,4,10-13] |
| auxin biosynthetic process | BP:0009851 | (0.121) | | | | [14-16] |
| auxin conjugate metabolic process | BP:0010249 | **** (< 4e-12) | **** (< 3e-24) | **** (< 3e-9) | Brassicales | [14-16] |
| auxin mediated signaling pathway involved in phyllotactic patterning | BP:0060774 | | | ** (< 0.005) | eudicots | [14-16] |
| auxin metabolic process | BP:0009850 | * (0.0474) | | | eudicots | [14-16] |
| basipetal auxin transport | BP:0010540 | | *** (< 8e-4) | . (0.0869) | | [14-16] |
| cell development | BP:0048468 | . (0.0634) | | | | [15,17] |
| cell differentiation | BP:0030154 | **** (< 2e-6) | **** (< 2e-6) | | | [15,17] |
| cell fate commitment | BP:0045165 | | *** (< 5e-4) | ** (< 0.005) | eudicots | [15,17] |
| cell fate speciation | BP:0001708 | | *** (< 4e-4) | ** (< 0.004) | | [15,17] |
| cell growth | BP:0016049 | . (0.0943) | | | | [15,17] |
| cell morphogenesis | BP:0000902 | **** (<2e-6) | | | | [15,17] |
| cell plate assembly | BP:0000919 | | | ** (< 0.002) | | |
| cell plate formation involved in | BP:0009920 | **** | ** | | | |

| | | | | | | |
|---|---|---|---|---|---|---|
| plant-type cell wall biogenesis | | (< 9e-17) | (0.005) | | | |
| cell population proliferation | BP:0008283 | . (0.0933) | | | | 14-16 |
| cell septum assembly | BP:0090529 | **** (< 9e-12) | | | eudicots | |
| cell tip growth | BP:0009932 | | **** (< 2e-5) | **** (< 8e-7) | 21 | 16,18 |
| cell wall modification | BP:0042545 | *** (7e-4) | | | | 17,19,20 |
| cellular response to auxin stimulus | BP:0071365 | | | * (0.0183) | | 14-16 |
| cellular response to freezing | BP:0071497 | ** (< 0.002) | **** (< 7e-15) | **** (< 3e-9) | | 21,22 |
| cellular response to hydrogen peroxide | BP:0070301 | **** (< 6e-7) | | | | 23 |
| cellular response to hypoxia | BP:0071456 | **** (< 5e-21) | | | | 24 (human) |
| cellular response to jasmonic acid stimulus | BP:0071395 | **** (< 3e-8) | | | | 25-27 |
| cellular response to sulphur starvation | BP:0010438 | **** (< 8e-9) | **** (< 3e-17) | **** (< 5e-6) | | 28,29 |
| cellular response to water deprivation | BP:0042631 | **** (< 6e-5) | | | | 30,31 |
| cuticle hydrocarbon biosynthetic process | BP:0006723 | | | *** (< 2e-4) | Eurosids I + Eurosids II | 32-34 |
| cuticle pattern formation | BP:0035017 | | ** (0.0016) | *** (< 8e-4) | eudicots | 32-34 |
| cytokinin metabolic process | BP:0009690 | | | * (0.0229) | eudicots | 35 |
| defense response | BP:0006952 | ** (0.0033) | | | | 36-44 |
| defense response to fungus, incompatible interaction | BP:0009817 | **** (< 5e-9) | | | | 37-40 |
| defense response to insect | BP:0002213 | **** (< 7e-7) | **** (< 8e-6) | | | 41-43 |
| defense response to oomycetes | BP:0002229 | | ** (0.0043) | **** (5e-6) | | 36,44 |
| developmental growth involved in morphogenesis | BP:0060560 | *** (< 2e-4) | | | | 6,15,17,45 |
| embryo development | BP:0009790 | ** (0.0075) | * (0.0283) | | | 46,47 |
| embryo development ending in seed dormancy | BP:0009793 | **** (< 4e-6) | | | | 46,47 |
| embryo sac development | BP:0009553 | | **** (< 5e-9) | **** (< 4e-8) | 40 | 48 |
| ethylene-activated signaling pathway | BP:000987 | **** (< 3e-16) | **** (< 9e-5) | | | 49 |

| | | | | | | |
|---|---|---|---|---|---|---|
| flower development | BP:0009908 | **** (< 8e-6) | | | | [3,4,10-13] |
| guard cell fate commitment | BP:0010377 | | * (0.0315) | | eudicots | [15] |
| hormone metabolic process | BP:0042445 | . (0.0615) | | | | see related terms to auxin, ABA, JA, SA, ethylene, BR, GA |
| hormone-mediated signaling pathway | BP:0009755 | **** (< 5e28) | **** (< 5e-5) | | | see related terms to auxin, ABA, JA, SA, ethylene, BR, GA |
| hyperosmotic response | BP:0006972 | ** (0.0024) | | | | [50] |
| jasmonic acid and ethylene-dependent systemic resistance | BP:0009861 | *** (< 2e-4) | | | | see related terms to JA and ethylene |
| jasmonic acid biosynthetic process | BP:0009695 | ** (0.0010) | . (0.0679) | | | [25-27] |
| jasmonic acid metabolic process | BP:0009694 | * (0.0304) | | | | [25-27] |
| lateral root development | BP:0048527 | **** (< 5e-6) | (0.1756) | | | [51-54] |
| lateral root morphogenesis | BP:0010102 | | *** (< 9e-4) | | Eurosids I + Eurosids II | [51-54] |
| leaf development | BP:0048366 | . (0.0557) | | | | [15] |
| leaf senescence | BP:0010150 | * (0.0389) | | | | [55,56] |
| lipid storage | BP:0019915 | * (0.0418) | | | | [57,58] |
| lipid transport | BP:0006869 | * (0.0205) | | | | [57,58] |
| maintenance of seed dormancy by abscisic acid | BP:0098755 | *** (< 4e-4) | **** (< 2e-11) | *** (< 2e-4) | eudicots | [46,47] |
| multicellular organism development | BP:0007275 | **** (< 7e-9) | | | | see related terms to auxin, ABA, JA, SA, ethylene, BR, GA |
| negative gravitropism | BP:0009959 | ** (0.0045) | | | | [52] |
| negative regulation of abscisic acid-activated signaling pathway | BP:0009788 | | | * (0.0112) | | [25,49,59,60] |
| negative regulation of brassinosteroid mediated signaling pathway | BP:1900458 | **** (< 5e-7) | | | 60 | [61,62] |
| negative regulation of cell division | BP:1900458 | | **** (< 4e-6) | **** (< 5e-6) | | [15,17] |
| negative regulation of flower development | | | | * (0.0425) | | [3,4,10-13] |
| negative regulation of gibberellin biosynthetic process | BP:0010373 | | ** (0.0017) | (0.1177) | Eurosids I + Eurosids II | [25,63,64] |

| | | | | | | |
|---|---|---|---|---|---|---|
| negative regulation of lateral root development | BP:1901332 | | **** (< 7e-9) | **** (< 5e-7) | | 51-54 |
| negative regulation of response to water deprivation | BP:0080148 | (0.2066) | *** (< 2e-4) | | | 30,31 |
| negative regulation of seed germination | BP:0010187 | | . (0.0523) | | | 46,47,65 |
| negative regulation of seed maturation | BP:2000692 | | **** (<3e-5) | *** (< 2e-4) | Eurosids II | 46,47,65 |
| petal formation | BP:0048451 | | (0.1765) | * (0.0310) | | 66,67 |
| pigment accumulation in response to UV light | BP:0043478 | **** (< 4e-5) | | | | 68,69 |
| pigment accumulation in tissues | BP:0043480 | **** (< 4e-5) | | | | 68,69 |
| plant epidermal cell differentiation | BP:0043480 | . (0.0592) | **** (< 2e-7) | ** (0.0048) | | 15 |
| plant epidermis morphogenesis | BP:0090627 | | ** (0.0020) | *** (< 2e-4) | | 15 |
| plant-type cell wall cellulose biosynthetic process | BP:0052324 | *** (< 3e-4) | | | | 19,20 |
| plant-type cell wall modification | BP:0009827 | ** (0.0048) | | | | 19,20 |
| plant-type cell wall organization or biogenesis | BP:0071669 | **** (< 4e-6) | | | | 17,19,20 |
| plant-type primary cell wall biogenesis | BP:0009833 | ** (0.0012) | | | | 17,19,20 |
| polarity specification of adaxial/abaxial axis | BP:0009944 | | | *** (< 3e-4) | eudicots | 14-16 |
| pollen development | BP:0009555 | . (0.0688) | | | | 3,4,10-12,70,71 |
| pollen germination | BP:0009846 | **** (< 2e-8) | **** (< 7e-16) | **** (< 2e-5) | | 70,71 |
| pollen maturation | BP:0010152 | . (0.0607) | | | 80 | 3,4,10-12 |
| pollen tube growth | BP:0009860 | **** (< 3e-8) | | | | 71,72 |
| pollen wall assembly | BP:0010208 | | ** (0.0044) | **** (< 3e-9) | | 3,4,10,12 |
| positive regulation of abscisic acid biosynthetic process | BP:0010116 | | *** (< 4e-4) | ** (0.0091) | eudicots | 25,49,59,60 |
| positive regulation of cell differentiation | BP:0045597 | . (0.0522) | ** (0.0066) | | | 15,17 |
| positive regulation of cell division | BP:0051781 | **** (< 3e-10) | ** (0.0038) | | | 15,17 |
| positive regulation of cell population proliferation | BP:0008284 | | | * (0.0405) | | 14-16 |
| positive regulation of defense | BP:1902290 | | ** | | eudicots | 36,44 |

| | | | | | | |
|---|---|---|---|---|---|---|
| response to oomycetes | | | (0.0036) | | | |
| positive regulation of defense response to virus by host | BP:0002230 | | *** (< 2e-4) | ** (0.0046) | eudicots | 73 |
| positive regulation of gibberellin biosynthetic process | BP:0010372 | | * (0.0243) | | Eurosids I + Eurosids II | 25,63,64 |
| positive regulation of leaf senescence | BP:1900057 | **** (< 4e-14) | **** (< 1e-6) | | | 55,56 |
| positive regulation of plant epidermal cell differentiation | BP:1903890 | **** (< 3e-5) | | | | 15 |
| positive regulation of seed maturation | BP:2000693 | . (0.0635) | | | | 46,47,65 |
| primary root development | BP:0080022 | **** (< 2e-5) | | | | 74,75 |
| regulation of abscisic acid12-activated signaling pathway | BP:0009787 | **** (< 6e-6) | | | | 25,49,59,60 |
| regulation of auxin mediated signaling pathway | BP:0010928 | **** (< 3e-5) | | | | 14-16 |
| regulation of auxin polar transport | BP:2000012 | **** (< 2e-18) | **** (< 3e-9) | | | 14,15 |
| regulation of brassinosteroid biosynthetic process | BP:0010422 | . (0.0851) | | | eudicots | 61,62 |
| regulation of cellular response to heat | BP:1900034 | | * (0.0239) | | eudicots | 76 |
| regulation of cellular response to stress | BP:0080135 | (0.1539) | * (0.0383) | | 100 | see biotic stress-, heat-, cold-, drought-, salt-, light-, UV-related terms |
| regulation of flower development | BP:0009909 | **** (< 8e-6) | **** (< 2e-10) | * (0.0418) | | 3,4,10-13 |
| regulation of growth | BP:0040008 | **** (< 9e-27) | **** (< 2e-4) | | | 6,45 |
| regulation of pollen tube growth | BP:0080092 | | *** (< 4e-4) | | | 71,72 |
| regulation of response to biotic stimulus | BP:0002831 | | ** (0.0024) | **** (< 3e-5) | eudicots | 41-43 |
| regulation of response to external stimulus | BP:0032101 | | * (0.0104) | **** (< 3e-5) | eudicots | see biotic stress-, heat-, cold-, drought-, salt-, light-, UV-related terms |
| regulation of response to osmotic stress | BP:0047484 | | | . (0.0514) | | 50 |
| regulation of response to salt stress | BP:1901000 | * (0.0423) | | | | 77-79 |
| regulation of root development | BP:2000280 | . (0.0946) | | | | 74,75 |
| regulation of stomatal closure | BP:0090333 | ** (0.0070) | | | | 15 |
| regulation of sulfur utilization | BP:0006792 | ** (0.0010) | **** (< 2e-8) | * (0.0342) | eudicots | 28,29 |

| | | | | | | |
|---|---|---|---|---|---|---|
| response to abiotic stimulus | BP:0009628 | **** (< 7e-13) | | | | see heat-, cold-, drought-, salt-, light-, UV-related terms |
| response to abscisic acid | BP:0009737 | **** (< 8e-9) | | | | 25,49,59,60 |
| response to auxin | BP:0009733 | **** (<7e-102) | * (0.0133) | | | 14-16 |
| response to cold | BP:0009409 | **** (< 2e-8) | | | | 21,22 |
| response to extracellular stimulus | BP:0009991 | **** (< 3e-5) | | | | see biotic stress-, heat-, cold-, drought-, salt-, light-, UV-related terms |
| response to heat | BP:0009408 | **** (< 7e-5) | **** (< 7e-10) | ** (0.0047) | | 76 |
| response to hydroperoxide | BP:0033194 | **** (< 6e-7) | . (0.0515) | | | 23 |
| response to jasmonic acid | BP:0009753 | **** (< 3e-14) | | | | 25-27 |
| response to karrikin | BP:0080167 | **** (< 6e-17) | | | | 80,81 |
| response to low fluence red light stimulus | BP:0010202 | **** (< 4e-5) | | | eudicots | 82 |
| response to low humidity | BP:0090547 | *** (< 3e-4) | **** (< 6e-12) | **** (< 5e-5) | eudicots | 30,31 |
| response to molecule of oomycetes origin | BP:0002240 | * (0.0439) | | | 122 | 36,44 |
| response to oomycetes | BP:0002239 | . (0.0768) | | | | 36,44 |
| response to oxidative stress | BP:0006979 | *** (< 2e-4) | | | | 31,72 |
| response to salicylic acid | BP:0009751 | **** (< 2e-13) | (0.1231) | | | 83-85 |
| response to salt | BP:1902074 | | * (0.0306) | | | 77-79 |
| response to salt stress | BP:0009651 | **** (< 2e-7) | ** (0.0016) | | | 77-79 |
| response to water deprivation | BP:0009414 | **** (< 7e-8) | *** (< 2e-4) | | | 30,31 |
| response to wounding | BP:0009611 | **** (< 1e-6) | | | | 85 |
| root development | BP:0048364 | * (0.0101) | | | | 74,75 |
| root hair elongation | BP:0048767 | **** (< 8e-8) | | | | 74,75 |
| salicylic acid metabolic process | BP:0009696 | * (0.0345) | | | | 83-85 |
| seed germination | BP:0009845 | **** (< 2e-7) | ** (0.0024) | | | 46,47,65 |

| | | | | | | |
|---|---|---|---|---|---|---|
| shoot system morphogenesis | BP:0010016 | * (0.0345) | ** (0.0011) | | eudicots | 14,15,86 |
| stigma development | BP:0048480 | | | . (0.0637) | eudicots | 3,4,10-13 |
| stomatal complex development | BP:0010374 | **** (< 2e-6) | | | | 15 |
| stomatal movement | BP:0010118 | ** (0.0057) | | | | 15 |
| trichome morphogenesis | BP:0010090 | ** (0.0061) | | | | 15 |
| wax biosynthetic process | BP:0010025 | * (0.0133) | | | 139 | 32-34 |
| *upstream or downstream processes of polyketide biosynthetic pathway* | | | | | | |
| acetate metabolic process | BP:0006083 | * (0.0380) | **** (< 6e-10) | **** (< 2e-5) | | |
| acetyl-CoA biosynthetic process from acetate | BP:0019427 | **** (< 1e-12) | **** (< 4e-27) | **** (< 5e-12) | | |
| cinnamic acid ester metabolic process | BP:0009801 | * (0.0434) | | | Asterids + Eurosids II | |
| cyanidin 3-O-glycoside metabolic process | BP:1901038 | | . (0.0860) | | Eurosids II + Core Eudicots | |
| fatty acid metabolic process | BP:0006631 | | ** (0.0074) | ** (0.0044) | | |
| flavone metabolic process | BP:0051552 | | * (0.0159) | ** (0.0052) | | |
| flavonoid biosynthetic process | BP:0072388 | | | . (0.0567) | Brassicales | |
| flavonoid biosynthetic process | BP:0009813 | | | **** (< 4e-8) | | |
| glucose metabolic process | BP:0006006 | | *** (< 4e-4) | **** (< 2e-7) | | |
| glycoside catabolic process | BP:0016139 | | * (0.0251) | . (0.0790) | eudicots | |
| negative regulation of secondary metabolite biosynthetic process | BP:1900377 | **** (< 7e-7) | | | | |
| positive regulation of secondary metabolite biosynthetic process | BP:1900378 | | | * (0.0271) | | |
| regulation of anthocyanin catabolic process | BP:1900000 | | *** (< 5e-4) | *** (< 8e-4) | Brassicales | |
| regulation of flavonol biosynthetic process | BP:1900384 | ** (0.0015) | * (0.0217) | | eudicots | |
| regulation of syringal lignin biosynthetic process | BP:1901428 | | | ** (0.0067) | | |
| shikimate biosynthetic process | BP:0033587 | | *** (< 3e-4) | *** (< 2e-4) | | |

Table S6: GO enrichment analysis of syntenic regions against the background 'all genes'. Given are enriched terms for 'biological process' that are potentially linked to flavonoid metabolism. Enrichment tests were done for the following cases: genes in PKS syntenic regions against background (all genes, PKS-BG), genes in CHS-enriched syntenic regions against background (CHS-BG) and genes in CHS-enriched syntenic regions against all genes in PKS syntenic regions (CHS-PKS). PKS genes were deleted from the syntenic regions prior to performing the enrichment analysis. p-values were adjusted for multiple testing by the Benjamini Hochberg method (q-values: . < 0.1, * ≤ 0.05, ** ≤ 0.01, *** ≤ 0.001, **** ≤ 0.0001). ABA: abscisic acid, JA: jasmonic acid, SA: salicylic acid, BR: brassinosteroids, GA: gibberellins.

| description | ID | PKS-BG | CHS-BG | CHS-PKS | clade specificity | reference |
|---|---|---|---|---|---|---|
| *direct and indirect effects of polyketides on biological processes* | | | | | | |
| actin cortical patch assembly | BP:0000147 | . (0.0513) | **** (< 4e-5) | | | [7] (human), [8] (in vitro), [9] (correlation) |
| actin cytoskeletal reorganization | BP:0031532 | * (0.0192) | | | | [7] (human), [8] (in vitro), [9] (correlation) |
| actin filament depolymerization | BP:0030042 | **** (< 5e-16) | **** (< 2e-6) | | | [7] (human), [8] (in vitro), [9] (correlation) |
| actin filament polymerization | BP:0030041 | **** (< 1e-17) | **** (< 3e-21) | *** (< 4e-4) | | [7] (human), [8] (in vitro), [9] (correlation) |
| anther development | BP:0048653 | **** (< 8e-8) | **** (< 7e-6) | | | [3,4,10-13] |
| auxin biosynthetic process | BP:0009851 | * (0.0103) | | | | [14-16] |
| auxin conjugate metabolic process | BP:0010249 | **** (< 8e-13) | | | Brassicales | [14-16] |
| auxin metabolic process | BP:0009850 | ** (0.0026) | | | eudicots | [14-16] |
| brassinosteroid mediated signaling pathway | BP:0009742 | *** (< 3e-4) | | | | [61,62] |
| cell development | BP:0048468 | * (0.0161) | | | | [15,17] |
| cell differentiation | BP:0030154 | **** (< 7e-13) | | | | [15,17] |
| cell growth | BP:0016049 | ** (0.0019) | | | | [15,17] |
| cell morphogenesis | BP:0000902 | **** (< 4e-7) | | | | [15,17] |
| cell plate formation involved in plant-type cell wall biogenesis | BP:0009920 | **** (< 2e-18) | **** (< 7e-9) | | | |
| cell population proliferation | BP:0008283 | * (0.0449) | | | | [14-16] |
| cell septum assembly | BP:0090529 | **** (< 4e-11) | | | eudicots | |
| cell wall biogenesis | BP:0042546 | ** (0.0015) | | | | [17,19,20] |
| cell wall modification | BP:0042545 | **** | **** | **** | | [17,19,20] |

| | | | | | | |
|---|---|---|---|---|---|---|
| | | (< 2e-6) | (< 6e-18) | (< 5e-9) | | |
| cell wall polysaccharide biosynthetic process | BP:0070592 | . (0.0721) | | | | [17,19,20] |
| cellular response to abscisic acid stimulus | BP:0071215 | ** (0.0016) | | | | [25,46,47,49,59,60] |
| cellular response to cold | BP:0070417 | | ** (0.0068) | * (0.0473) | | [21,22] |
| cellular response to freezing | BP:0071497 | **** (< 5e-5) | | | | [21,22] |
| cellular response to hormone stimulus | BP:0032870 | * (0.0420) | | | | see related terms to auxin, ABA, JA, SA, ethylene, BR, GA |
| cellular response to hydrogen peroxide | BP:0070301 | **** (< 6e-7) | **** (< 2e-9) | | | [23] |
| cellular response to hypoxia | BP:0071456 | **** (< 3e-27) | | | | [24] (human) |
| cellular response to jasmonic acid stimulus | BP:0071395 | **** (< 4e-10) | ** (0.0060) | | | [25-27] |
| cellular response to salt stress | BP:0071472 | . (0.0527) | | | | [77-79] |
| cellular response to sulphur starvation | BP:0010438 | **** (< 2e-10) | **** (< 3e-13) | * (0.0425) | | [28,29] |
| cellular response to water deprivation | BP:0042631 | **** (< 6e-8) | | | | [30,31] |
| cytokinin transport | BP:0010184 | | . (0.0507) | . (0.0678) | eudicots | [35] |
| cytokinin-activated signalling pathway | BP:0009736 | * (0.0436) | * (0.0313) | | | [35,87] |
| defense response | BP:0006952 | **** (< 3e-9) | ** (0.0023) | | | [36-44] |
| defense response to fungus | BP:0050832 | **** (< 3e-5) | | | | [37-40] |
| defense response to fungus, incompatible interaction | BP:0009817 | **** (< 3e-11) | **** (< 2e-5) | | | [37-40] |
| defense response to insect | BP:0002213 | **** (< 5e-9) | | | | [41-43] |
| detection of external biotic stimulus | BP:0098581 | * (0.0389) | | | | [37-43] |
| detection of fungus | BP:0016046 | **** (< 4e-9) | | | Brassicales | [37-40] |
| developmental growth involved in morphogenesis | BP:0060560 | **** (< 4e-7) | | | | [6,15,17,45] |
| embryo development | BP:0009790 | ** (0.0014) | | | | [46,47] |
| embryo development ending in seed dormancy | BP:0009793 | **** (< 9e-7) | | | | [46,47] |
| embryo sac cellularization | BP:0009558 | | | * | eudicots | [48] |

| | | | | | | |
|---|---|---|---|---|---|---|
| | | | | (0.0139) | | |
| embryo sac development | BP:0009553 | | *** (< 2e-4) | * (0.0140) | | 48 |
| ethylene metabolic process | BP:0009692 | | | **** (< 4e-7) | | 49 |
| ethylene-activated signaling pathway | BP:000987 | **** (< 2e-24) | **** (< 4e-20) | . (0.0833) | | 49 |
| floral organ morphogenesis | BP:0048444 | | * (0.0214) | . (0.0768) | | 3,4,10-13 |
| flower development | BP:0009908 | **** (< 7e-10) | | | | 3,4,10-13 |
| gibberellin in metabolic process | BP:0009685 | | | ** (0.0041) | | 25,63,64 |
| hormone biosynthetic process | BP:0042446 | | *** (< 2e-4) | **** (< 4e-9) | | see related terms to auxin, ABA, JA, SA, ethylene, BR, GA |
| hormone metabolic process | BP:0042445 | | **** (< 3e-5) | (0.1652) | | see related terms to auxin, ABA, JA, SA, ethylene, BR, GA |
| hormone-mediated signaling pathway | BP:0009755 | **** (< 2e-40) | **** (< 3e-7) | | | see related terms to auxin, ABA, JA, SA, ethylene, BR, GA |
| hyperosmotic response | BP:0006972 | *** (< 3e-4) | | | | 50 |
| jasmonic acid and ethylene-dependent systemic resistance | BP:0009861 | **** (< 6e-6) | ** (0.0032) | | | see related terms to JA and ethylene |
| jasmonic acid biosynthetic process | BP:0009695 | *** (< 4e-4) | **** (< 4e-12) | **** (< 3e-5) | | 25-27 |
| jasmonic acid metabolic process | BP:0009694 | ** (0.0012) | (0.1181) | | | 25-27 |
| lateral root development | BP:0048527 | **** (< 2e-8) | . (0.0574) | | | 51-54 |
| lateral root formation | BP:0010311 | * (0.0138) | | | | 51-54 |
| lateral root morphogenesis | BP:0010102 | . (0.0917) | | | Eurosids I + Eurosids II | 51-54 |
| leaf development | BP:0048366 | *** (< 4e-4) | | | | 15 |
| leaf senescence | BP:0010150 | **** (< 2e-5) | | | | 55,56 |
| lipid storage | BP:0019915 | . (0.0625) | | | | 57,58 |
| lipid transport | BP:0006869 | ** (0.0084) | | | | 57,58 |
| maintenance of seed dormancy by abscisic acid | BP:0098755 | *** (< 2e-4) | | | eudicots | 46,47 |
| multicellular organism development | BP:0007275 | **** (< 5e-14) | | | | see related terms to auxin, ABA, JA, SA, ethylene, |

| | | | | | | |
|---|---|---|---|---|---|---|
| | | | | | | BR, GA |
| negative gravitropism | BP:0009959 | ** (0.0015) | . (0.0551) | | | 52 |
| negative regulation of brassinosteroid mediated signaling pathway | BP:1900458 | **** (< 9e-8) | ** (0.0095) | | | 61,62 |
| negative regulation of cell fate commitment | BP:0010454 | | *** (< 3e-4) | | | 15,17 |
| negative regulation of defense response to insect | BP:1900366 | | | . (0.0691) | | 41-43 |
| negative regulation of lateral root development | BP:1901332 | | **** (< 3e-6) | * (0.0164) | | 51-54 |
| negative regulation of response to salt stress | BP:0071156 | | * (0.0304) | | eudicots | 77-79 |
| negative regulation of response to water deprivation | BP:0080148 | * (0.0235) | * (0.0146) | | | 30,31 |
| pigment accumulation in response to UV light | BP:0043478 | **** (< 7e-6) | *** (< 2e-4) | | | 68,69 |
| pigment accumulation in tissues | BP:0043480 | **** (< 7e-6) | *** (< 2e-4) | | | 68,69 |
| plant epidermal cell differentiation | BP:0043480 | ** (0.0012) | * (0.0366) | | | 15 |
| plant-type cell wall assembly | BP:0071668 | | ** (0.0021) | **** (< 3e-5) | eudicots | 17,19,20 |
| plant-type cell wall cellulose biosynthetic process | BP:0052324 | *** (< 2e-5) | | | | 19,20 |
| plant-type cell wall modification | BP:0009827 | **** (< 8e-5) | | | | 19,20 |
| plant-type cell wall organization or biogenesis | BP:0071669 | **** (< 7e-9) | | | | 17,19,20 |
| plant-type primary cell wall biogenesis | BP:0009833 | **** (< 4e-5) | | | | 17,19,20 |
| pollen development | BP:0009555 | * (0.0212) | | | | 3,4,10-12,70,71 |
| pollen germination | BP:0009846 | **** (< 8e-10) | | | | 70,71 |
| pollen maturation | BP:0010152 | ** (0.0039) | | | | 3,4,10-12 |
| pollen tube growth | BP:0009860 | **** (< 2e-11) | | | | 71,72 |
| positive regulation of abscisic acid-activated signaling pathway | BP:0010116 | * (0.0199) | | | eudicots | 25,49,59,60 |
| positive regulation of cell differentiation | BP:0045597 | * (0.0129) | ** (0.0268) | | | 15,17 |
| positive regulation of cell division | BP:0051781 | **** (< 6e-10) | **** (< 9e-19) | **** (< 1e-6) | | 15,17 |
| positive regulation of gibberellin | BP:0010372 | . | | | Eurosids I + | 25,63,64 |

| | | | | | | |
|---|---|---|---|---|---|---|
| biosynthetic process | | (0.0510) | | | Eurosids II | |
| positive regulation of leaf senescence | BP:1900057 | **** (< 2e-17) | **** (< 5e-6) | | | [55,56] |
| positive regulation of plant epidermal cell differentiation | BP:1903890 | *** (< 2e-5) | * (0.0286) | | | [15] |
| positive regulation of seed germination | BP:0010030 | ** (0.0046) | | | | [46,47,65] |
| positive regulation of seed maturation | BP:2000693 | ** (0.0059) | | | | [46,47,65] |
| primary root development | BP:0080022 | **** (< 5e-8) | | | | [74,75] |
| red, far-red light phototransduction | BP:0009585 | | **** (< 3e-5) | ** (0.0043) | | [82] |
| regulation of abscisic acid-activated signaling pathway | BP:0009787 | **** (< 1e-7) | | | | [25,49,59,60] |
| regulation of auxin mediated signaling pathway | BP:0010928 | **** (< 2e-5) | | | | [14-16] |
| regulation of auxin polar transport | BP:2000012 | **** (< 7e-21) | | | | [14,15] |
| regulation of brassinosteroid biosynthetic process | BP:0010422 | . (0.0710) | ** (0.0024) | | eudicots | [61,62] |
| regulation of cellular response to stress | BP:0080135 | * (0.0234) | | | | see biotic stress-, heat-, cold-, drought-, salt-, light-, UV-related terms |
| regulation of defense response to fungus | BP:1900150 | | **** (< 9e-6) | *** (< 7e-4) | | [37-40] |
| regulation of flower development | BP:0009909 | **** (< 1e-11) | | | | [3,4,10-13] |
| regulation of growth | BP:0040008 | **** (< 2e-34) | **** (< 7e-11) | | | [6,45] |
| regulation of leaf senescence | BP:1900055 | | ** (0.0095) | * (0.0195) | | [55,56] |
| regulation of pollen tube growth | BP:0080092 | * (0.0264) | | | | [71,72] |
| regulation of response to salt stress | BP:1901000 | ** (0.0022) | | | | [77-79] |
| regulation of root development | BP:2000280 | * (0.0113) | | | | [74,75] |
| regulation of salicylic acid metabolic process | BP:0010337 | | | ** (0.0019) | | [83-85] |
| regulation of stomatal closure | BP:0090333 | *** (< 2e-4) | | | | [15] |
| regulation of stomatal opening | BP:1902456 | * (0.0252) | | | | [15] |
| regulation of sulfur utilization | BP:0006792 | *** (< 2e-4) | **** (< 4e-13) | **** (< 5e-6) | eudicots | [28,29] |
| response to abiotic stimulus | BP:0009628 | **** | **** | | | see heat-, cold-, drought-, |

| | | | | | | |
|---|---|---|---|---|---|---|
| | | (< 2e-15) | (< 4e-5) | | | salt-, light-, UV-related terms |
| response to abscisic acid | BP:0009737 | **** (< 2e-18) | **** (< 4e-5) | | | [25,49,59,60] |
| response to auxin | BP:0009733 | **** (<5e-127) | . (0.0574) | | | [14-16] |
| response to brassinosteroid | BP:0009741 | | . (0.0969) | | eudicots | [61,62] |
| response to cold | BP:0009409 | **** (< 6e-17) | (0.1162) | | | [21,22] |
| response to ethylene | BP:0009723 | *** (< 5e-4) | | | | [49] |
| response to extracellular stimulus | BP:0009991 | **** (< 2e-5) | | | | see biotic stress-, heat-, cold-, drought-, salt-, light-, UV-related terms |
| response to fungus | BP:0009620 | | **** (< 3e-5) | *** (< 6e-4) | | [37-40] |
| response to gibberellin | BP:0009739 | ** (0.0013) | | | | [25,63,64] |
| response to heat | BP:0009408 | **** (< 2e-8) | | | | [76] |
| response to high fluence blue light stimulus by blue high-fluence system | BP:0055121 | | * (0.0444) | | | [88,89] |
| response to hydroperoxide | BP:0033194 | **** (< 3e-7) | | | | [23] |
| response to jasmonic acid | BP:0009753 | **** (< 7e-23) | *** (< 3e-4) | | | [25-27] |
| response to karrikin | BP:0080167 | **** (< 4e-25) | | | | [80,81] |
| response to light stimulus | BP:0009416 | **** (< 3e-5) | (0.1531) | | | [68,69,82,88,89] |
| response to low fluence red light stimulus | BP:0010202 | **** (< 9e-6) | | | eudicots | [82] |
| response to low humidity | BP:0090547 | **** (< 2e-5) | **** (< 5e-10) | * (0.0334) | eudicots | [30,31] |
| response to molecule of oomycetes origin | BP:0002240 | ** (0.0017) | | | | [36,44] |
| response to oomycetes | BP:0002239 | *** (< 4e-4) | | | | [36,44] |
| response to oxidative stress | BP:0006979 | **** (< 2-6) | | | | [31,72] |
| response to red or far red light | BP:0009639 | | . (0.0643) | | | [82] |
| response to salicylic acid | BP:0009751 | **** (< 6e-23) | **** (< 7e-9) | | | [83-85] |
| response to salt | BP:1902074 | * (0.0283) | | | | [77-79] |

| | | | | | | |
|---|---|---|---|---|---|---|
| response to salt stress | BP:0009651 | ****<br>(< 3e-17) | | | | 77-79 |
| response to water deprivation | BP:0009414 | ****<br>(< 2e-16) | ***<br>(< 2e-4) | | | 30,31 |
| response to wounding | BP:0009611 | ****<br>(< 8e-13) | ****<br>(< 8e-7) | | | 85 |
| root development | BP:0048364 | ****<br>(< 9e-6) | *<br>(0.0210) | | | 74,75 |
| root hair cell development | BP:0080147 | **<br>(0.0073) | | | | 74,75 |
| root hair elongation | BP:0048767 | ****<br>(< 6e-11) | ***<br>(< 1e-3) | | | 74,75 |
| salicylic acid metabolic process | BP:0009696 | ***<br>(< 8e-4) | | | | 83-85 |
| seed coat development | BP:0010214 | *<br>(0.0240) | | | | 46,47,65 |
| seed development | BP:0048316 | *<br>(0.0365) | *<br>(0.0215) | | | 46,47,65 |
| seed dormany process | BP:0010162 | *<br>(0.0426) | | | | 46,47,65 |
| seed germination | BP:0009845 | ****<br>(< 5e-12) | | | | 46,47,65 |
| shoot system morphogenesis | BP:0010016 | ***<br>(< 6e-4) | ****<br>(< 2e-8) | *<br>(0.0407) | eudicots | 14,15,86 |
| stomatal closure | BP:0090332 | | **<br>(0.0078) | **<br>(0.0069) | | 15 |
| stomatal complex development | BP:0010374 | ****<br>(< 2e-8) | | | | 15 |
| stomatal movement | BP:0010118 | ****<br>(< 5e-5) | | | | 15 |
| trichome branching | BP:0010091 | *<br>(0.0499) | | | | 15 |
| trichome morphogenesis | BP:0010090 | ***<br>(< 5e-4) | ****<br>(< 7e-11) | ***<br>(< 2e-4) | | 15 |
| wax biosynthetic process | BP:0010025 | **<br>(0.0028) | | | | 32-34 |
| *upstream or downstream processes of polyketide biosynthetic pathway* | | | | | | |
| acetate metabolic process | BP:0006083 | *<br>(0.0413) | | | | |
| acetyl-CoA biosynthetic process from acetate | BP:0019427 | ****<br>(< 3e-11) | | | | |
| aromatic amino acid family metabolic process | BP:0009072 | | ****<br>(< 8e-5) | ****<br>(< 7e-10) | | |
| cinnamic acid ester metabolic process | BP:0009801 | *<br>(0.0167) | | | Asterids +<br>Eurosids II | |

| | | | | | |
|---|---|---|---|---|---|
| fatty acid beta-oxidation | BP:0006635 | | **** (< 8e-7) | *** (< 2e-4) | |
| isoflavonoid metabolic process | BP:0046287 | | | * (0.0205) | |
| lipid biosynthetic process | BP:0008610 | | **** (< 5e-5) | (0.1641) | |
| negative regulation of secondary metabolite biosynthetic process | BP:1900377 | **** (< 5e-8) | | | |
| positive regulation of fatty acid biosynthetic process | BP:0045723 | | | * (0.0311) | |
| regulation of flavonol biosynthetic process | BP:1900384 | *** (< 4e-4) | | | eudicots |

Table S7: Gene information for enriched GO terms of category ‘Biological process’ directly linking to flavonoid metabolism using the set with the background ‘syntenic regions’. PKS genes were removed prior to the enrichment analysis. The genes are listed according to their enriched processes ‘flavone metabolic process’, ‘flavonoid biosynthetic process’ , ‘regulation of anthocyanin catabolic process’ and ‘regulation of flavonol biosynthetic process’. Amino acid FASTA sequences were blasted against the blastp database.

| | gene | description | Query cover [%] | Identity [%] | E value |
|---|---|---|---|---|---|
| **flavonone metabolic process** | araly_AL6G28030.t1 | UDP-glycosyltransferase 78D3 [Arabidopsis lyrata subsp. lyrata] | 99 | 100 | 0 |
| | araly_AL6G28040.t1 | UDP-glycosyltransferase 78D4 isoform X3 [Arabidopsis lyrata subsp. lyrata] | 99 | 100 | 0 |
| | araly_AL6G28060.t1 | UDP-glycosyltransferase 78D2 [Arabidopsis lyrata subsp. lyrata] | 99 | 100 | 0 |
| | araly_AL6G36120.t1 | 2-oxoglutarate (2OG) and Fe(II)-dependent oxygenase superfamily protein [Arabidopsis thaliana] | 99 | 97.52 | 2e-114 |
| | arath_AT5G17030 | UDP-glucosyl transferase 78D3 [Arabidopsis thaliana] | 99 | 100 | 0 |
| | arath_AT5G17040 | UDP-Glycosyltransferase superfamily protein [Arabidopsis thaliana] | 99 | 100 | 0 |
| | arath_AT5G17050 | UDP-glucosyl transferase 78D2 [Arabidopsis thaliana] | 99 | 100 | 0 |
| | arath_AT5G24530 | 2-oxoglutarate (2OG) and Fe(II)-dependent oxygenase superfamily protein [Arabidopsis thaliana] | 99 | 100 | 0 |
| | brara_Brara.B00690.1.p | PREDICTED: UDP-glycosyltransferase 78D2-like isoform X3 [Brassica rapa] | 99 | 98.03 | 0 |
| | brara_Brara.B03677.1.p | flavanone 3-dioxygenase-like [Brassica oleracea var. oleracea] | 99 | 97.07 | 0 |
| | eutsa_Thhalv10013472m | UDP-glycosyltransferase 78D2 [Eutrema salsugineum] | 99 | 100 | 0 |
| | eutsa_Thhalv10015523m | flavanone 3-dioxygenase 3 [Eutrema salsugineum] | 99 | 95.57 | 0 |
| | glyma_Glyma.06G137000.1.p | DOWNY MILDEW RESISTANCE 6 [Glycine max] | 99 | 100 | 0 |
| | medtr_XP_003602395.1 | DOWNY MILDEW RESISTANCE 6 [Medicago truncatula] | 100 | 100 | 0 |
| | medtr_XP_003625205.1 | cytochrome P450 93A3 [Medicago truncatula] | 100 | 100 | 0 |
| | medtr_XP_003625207.1 | cytochrome P450 93A3 [Medicago truncatula] | 100 | 100 | 0 |
| | orysa_LOC_OS04G01140 | cytochrome P450 93G1-like [Oryza sativa Japonica Group] | 100 | 100 | 0 |
| | selmo_Smo12440\|PACid_15405409 | flavonoid 3'-monooxygenase [Selaginella moellendorffii] | 100 | 100 | 0 |
| | selmo_Smo85404\|PACid_15419886 | flavonoid 3'-monooxygenase [Selaginella moellendorffii] | 99 | 98.80 | 0 |
| | thepa_Tp6g27110 | UDP-glycosyltransferase 78D2 [Eutrema salsugineum] | 99 | 85.93 | 0 |
| **flavonoid biosynthetic process** | araly_AL6G12780.t1 | FAD synthase [Arabidopsis lyrata subsp. lyrata] | 99 | 100 | 0 |
| | araly_AL6G18780.t1 | FAD synthetase 2, chloroplastic [Arabidopsis lyrata subsp. lyrata] | 99 | 100 | 0 |
| | araly_AL6G34590.t1 | FAD synthetase 1, chloroplastic [Arabidopsis lyrata subsp. lyrata] | 99 | 100 | 0 |
| | arath_AT5G23330 | Nucleotidylyl transferase superfamily protein [Arabidopsis thaliana] | 99 | 100 | 0 |
| | arath_AT5G03430 | phosphoadenosine phosphosulfate (PAPS) reductase family protein [Arabidopsis thaliana] | 99 | 100 | 0 |
| | arath_AT5G08340 | Nucleotidylyl transferase superfamily protein [Arabidopsis thaliana] | 99 | 100 | 0 |
| | braol_BO3G002340.1 | FAD synthase-like [Brassica oleracea var. oleracea] | 100 | 100 | 0 |
| | braol_BO3G004870.1 | FAD synthetase 2, chloroplastic [Brassica oleracea var. oleracea] | 100 | 100 | 0 |
| | brara_Brara.B00097.1.p | FAD synthase [Brassica napus] | 99 | 98.21 | 0 |
| | brara_Brara.C00110.1.p | FAD synthase [Brassica napus] | 99 | 99.19 | 0 |
| | camsa_XP_010452817.1 | FAD synthetase 2, chloroplastic isoform X1 [Camelina sativa] | 100 | 100 | 0 |
| | camsa_XP_010456377.2 | FAD synthase-like [Camelina sativa] | 100 | 100 | 0 |
| | camsa_XP_010491465.1 | FAD synthetase 2, chloroplastic-like [Camelina sativa] | 100 | 100 | 0 |
| | camsa_XP_019097972.1 | FAD synthase isoform X1 [Camelina sativa] | 100 | 100 | 0 |
| | eutsa_Thhalv10013303m | FAD synthase [Eutrema salsugineum] | 99 | 98.42 | 0 |
| | eutsa_Thhalv10013886m | FAD synthetase 2, chloroplastic isoform X1 [Eutrema salsugineum] | 99 | 100 | 0 |
| | lepme_evm.model.scaffold98.515 | FAD synthetase 2, chloroplastic [Arabidopsis lyrata subsp. lyrata] | 48 | 82.48 | 1e-130 |
| | thepa_Tp6g34600 | FAD synthetase 2, chloroplastic isoform X1 [Eutrema salsugineum] | 100 | 89.07 | 0 |
| | thepa_Tp6g39310 | FAD synthase-like isoform X1 [Brassica napus] | 99 | 95.58 | 0 |
| | aradu_XP_015952592.2 | zinc finger protein WIP5-like [Arachis duranensis] | 100 | 100 | 0 |
| | aradu_XP_015959151.1 | heme oxygenase 1, chloroplastic [Arachis duranensis] | 100 | 100 | 0 |
| | aradu_XP_015964809.1 | 2-hydroxyisoflavanone dehydratase-like [Arachis duranensis] | 100 | 100 | 0 |
| | aradu_XP_015964810.1 | 2-hydroxyisoflavanone dehydratase [Arachis duranensis] | 100 | 100 | 0 |
| | aradu_XP_015964830.1 | 2-hydroxyisoflavanone dehydratase-like [Arachis duranensis] | 100 | 100 | 0 |
| | araly_AL6G15110.t1 | probable chalcone--flavonone isomerase 3 isoform X2 [Arabidopsis lyrata subsp. lyrata] | 100 | 100 | 2e-148 |
| | araly_AL6G15490.t1 | probable 2-oxoglutarate-dependent dioxygenase At5g05600 [Arabidopsis lyrata subsp. lyrata] | 99 | 100 | 0 |
| | araly_AL6G18330.t1 | flavonoid 3'-monooxygenase [Arabidopsis lyrata subsp. lyrata] | 99 | 100 | 0 |
| | araly_AL6G19370.t1 | flavonol synthase/flavanone 3-hydroxylase [Arabidopsis lyrata subsp. lyrata] | 99 | 100 | 0 |
| | araly_AL6G28030.t1 | UDP-glycosyltransferase 78D3 [Arabidopsis lyrata subsp. lyrata] | 99 | 100 | 0 |

| | | | | | |
|---|---|---|---|---|---|
| flavonoid biosynthetic process | araly_AL6G28040.t1 | UDP-glycosyltransferase 78D4 isoform X3 [Arabidopsis lyrata subsp. lyrata] | 99 | 100 | 0 |
| | araly_AL6G28060.t1 | UDP-glycosyltransferase 78D2 [Arabidopsis lyrata subsp. lyrata] | 99 | 100 | 0 |
| | araly_AL6G29400.t1 | ADP-glucose phosphorylase [Arabidopsis lyrata subsp. lyrata] | 99 | 100 | 0 |
| | araly_AL6G32080.t1 | probable 2-oxoglutarate-dependent dioxygenase At5g05600 [Arabidopsis lyrata subsp. lyrata] | 99 | 84.52 | 6e-96 |
| | araly_AL6G36120.t1 | 2-oxoglutarate (2OG) and Fe(II)-dependent oxygenase superfamily protein [Arabidopsis thaliana] | 99 | 97.52 | 2e-114 |
| | araip_XP_016191494.1 | zinc finger protein WIP2-like isoform X1 [Arachis ipaensis] | 100 | 100 | 0 |
| | araip_XP_016197624.1 | heme oxygenase 1, chloroplastic [Arachis ipaensis] | 100 | 100 | 0 |
| | arath_AT5G05270 | Chalcone-flavanone isomerase family protein [Arabidopsis thaliana] | 99 | 100 | 4e-151 |
| | arath _AT5G05600 | 2-oxoglutarate (2OG) and Fe(II)-dependent oxygenase superfamily protein [Arabidopsis thaliana] | 99 | 100 | 0 |
| | arath _AT5G17030 | UDP-glucosyl transferase 78D3 [Arabidopsis thaliana] | 99 | 100 | 0 |
| | arath _AT5G17040 | UDP-Glycosyltransferase superfamily protein [Arabidopsis thaliana] | 99 | 100 | 0 |
| | arath _AT5G17050 | UDP-glucosyl transferase 78D2 [Arabidopsis thaliana] | 99 | 100 | 0 |
| | arath _AT5G07480 | KAR-UP oxidoreductase 1 [Arabidopsis thaliana] | 99 | 95.51 | 0 |
| | arath _AT5G07990 | Cytochrome P450 superfamily protein [Arabidopsis thaliana] | 99 | 100 | 0 |
| | arath _AT5G08640 | flavonol synthase 1 [Arabidopsis thaliana] | 99 | 100 | 0 |
| | arath _AT5G18200 | putative galactose-1-phosphate uridyltransferase [Arabidopsis thaliana] | 99 | 100 | 0 |
| | arath _AT5G24530 | 2-oxoglutarate (2OG) and Fe(II)-dependent oxygenase superfamily protein [Arabidopsis thaliana] | 99 | 100 | 0 |
| | braol_BO3G003410.1 | flavonol synthase/flavanone 3-hydroxylase-like [Brassica oleracea var. oleracea] | 88 | 100 | 0 |
| | brara_Brara.B00184.1.p | flavonol synthase/flavanone 3-hydroxylase isoform X2 [Brassica rapa] | 99 | 100 | 0 |
| | brara_Brara.B00247.1.p | flavanone 3-dioxygenase 3-like isoform X1 [Brassica napus] | 99 | 94.74 | 0 |
| | brara_Brara.B00690.1.p | PREDICTED: UDP-glycosyltransferase 78D2-like isoform X3 [Brassica rapa] | 99 | 98.03 | 0 |
| | brara_Brara.B03677.1.p | flavanone 3-dioxygenase-like [Brassica oleracea var. oleracea] | 99 | 97.07 | 0 |
| | brara_Brara.B03857.1.p | flavonol synthase 3-like [Brassica rapa] | 99 | 98.71 | 0 |
| | brara _Brara.C00218.1.p | flavonol synthase/flavanone 3-hydroxylase-like [Brassica rapa] | 99 | 100 | 0 |
| | brara _Brara.J01576.1.p | codeine O-demethylase-like [Brassica rapa] | 99 | 100 | 0 |
| | brara _Brara.J01743.1.p | ADP-glucose phosphorylase [Brassica rapa] | 99 | 100 | 0 |
| | brara _Brara.J02408.1.p | flavonol synthase/flavanone 3-hydroxylase [Brassica napus] | 94 | 100 | 0 |
| | camsa_XP_010452429.1 | probable chalcone--flavonone isomerase 3 [Camelina sativa] | 100 | 100 | 3e-152 |
| | camsa_XP_010452770.1 | flavonoid 3'-monooxygenase [Camelina sativa] | 100 | 100 | 0 |
| | camsa_XP_010452467.1 | flavonol synthase/flavanone 3-hydroxylase isoform X2 [Camelina sativa] | 100 | 100 | 0 |
| | camsa_XP_010455373.1 | transcription factor TT8-like [Camelina sativa] | 100 | 100 | 0 |
| | camsa_XP_010491060.1 | probable chalcone--flavonone isomerase 3 [Camelina sativa] | 100 | 100 | 2e-151 |
| | camsa_XP_010491092.1 | flavonol synthase/flavanone 3-hydroxylase-like [Camelina sativa] | 100 | 100 | 0 |
| | camsa_XP_010491421.1 | flavonoid 3'-monooxygenase-like [Camelina sativa] | 100 | 100 | 0 |
| | cicar_Ca_08111 | cytochrome P450 98A2 [Cicer arietinum] | 100 | 100 | 0 |
| | cicar_Ca_09533 | vestitone reductase isoform X2 [Cicer arietinum] | 100 | 100 | 0 |
| | cicar_Ca_11884 | 2-hydroxyisoflavanone dehydratase-like [Cicer arietinum] | 100 | 100 | 0 |
| | cicar_Ca_11885 | 2-hydroxyisoflavanone dehydratase-like [Cicer arietinum] | 100 | 100 | 0 |
| | cicar_Ca_19207 | NAD(P)H-dependent 6'-deoxychalcone synthase-like [Cicer arietinum] | 100 | 100 | 0 |
| | cicar_Ca_19221 | NAD(P)H-dependent 6'-deoxychalcone synthase [Cicer arietinum] | 100 | 100 | 0 |
| | cicar_Ca_19222 | NAD(P)H-dependent 6'-deoxychalcone synthase-like [Cicer arietinum] | 100 | 100 | 0 |
| | citcl_XP_006423717.1 | vestitone reductase [Citrus clementina] | 100 | 100 | 0 |
| | citcl_XP_006423723.1 | vestitone reductase [Citrus clementina] | 100 | 96.17 | 0 |
| | citcl_XP_006433965.1 | protein TRANSPARENT TESTA 1 [Citrus clementina] | 100 | 100 | 0 |
| | citcl_XP_006434471.1 | sterol 3-beta-glucosyltransferase UGT80B1 [Citrus clementina] | 100 | 100 | 0 |
| | citcl_XP_006434819.1 | probable inactive heme oxygenase 2, chloroplastic [Citrus clementina] | 100 | 100 | 0 |
| | citcl_XP_006446775.1 | Protein TRANSPARENT TESTA 12 [Morella rubra] | 96 | 55.68 | 2e-147 |
| | citcl_XP_006446841.1 | TT12-2 MATE transporter [Theobroma cacao] | 99 | 87.45 | 0 |
| | citsi_Cs2g15080.1 | TT12-2 MATE transporter [Theobroma cacao] | 98 | 80 | 0 |
| | citsi_Cs2g15120.1 | TT12-2 MATE transporter [Theobroma cacao] | 99 | 86.85 | 0 |
| | citsi_Cs2g15610.1 | TT12-2 MATE transporter [Theobroma cacao] | 87 | 62.23 | 6e-80 |
| | citsi_Cs3g18120.1 | protein TRANSPARENT TESTA 1 [Citrus sinensis] | 99 | 100 | 0 |
| | citsi_Cs3g19280.1 | flavonol synthase/flavanone 3-hydroxylase-like [Citrus sinensis] | 99 | 100 | 0 |
| | citsi_Cs3g21440.1 | sterol 3-beta-glucosyltransferase UGT80B1 isoform X1 [Citrus sinensis] | 99 | 95.17 | 0 |
| | citsi_Cs3g23830.1 | probable inactive heme oxygenase 2, chloroplastic [Citrus sinensis] | 99 | 100 | 0 |
| | dauca_DCAR_018993 | bergaptol O-methyltransferase-like [Daucus carota subsp. sativus] | 78 | 99.43 | 0 |
| | dauca_DCAR_018995 | bergaptol O-methyltransferase-like [Daucus carota subsp. sativus] | 86 | 100 | 0 |
| | eucgr_Eucgr.H03555.1.p | 2-hydroxyisoflavanone dehydratase isoform X1 [Eucalyptus grandis] | 85 | 100 | 0 |
| | eutsa_Thhalv10015523m | flavanone 3-dioxygenase 3 [Eutrema salsugineum] | 99 | 95.57 | 0 |
| | eutsa_Thhalv10013289m | flavonoid 3'-monooxygenase [Eutrema salsugineum] | 99 | 100 | 0 |

| | | | | | |
|---|---|---|---|---|---|
| flavonoid biosynthetic process | eutsa_Thhalv10013472m | UDP-glycosyltransferase 78D2 [Eutrema salsugineum] | 99 | 100 | 0 |
| | eutsa_Thhalv10013861m | probable 2-oxoglutarate-dependent dioxygenase At5g05600 [Eutrema salsugineum] | 99 | 100 | 0 |
| | eutsa_Thhalv10013989m | codeine O-demethylase-like [Brassica rapa] | 99 | 87.93 | 0 |
| | eutsa_Thhalv10013996m | ADP-glucose phosphorylase [Eutrema salsugineum] | 99 | 100 | 0 |
| | eutsa_Thhalv10014054m | flavonol synthase/flavanone 3-hydroxylase [Eutrema salsugineum] | 99 | 100 | 0 |
| | eutsa_Thhalv10014675m | probable chalcone--flavonone isomerase 3 isoform X1 [Eutrema salsugineum] | 99 | 100 | 4e-151 |
| | frave_mrna11126.1-v1.0-hybrid | flavonol synthase/flavanone 3-hydroxylase [Fragaria vesca subsp. vesca] | 76 | 100 | 2e-124 |
| | glyma_Glyma.06G110600.1.p | flavonol synthase/flavanone 3-hydroxylase [Glycine max] | 99 | 100 | 0 |
| | glyma_Glyma.06G137000.1.p | DOWNY MILDEW RESISTANCE 6 [Glycine max] | 99 | 100 | 0 |
| | glyma_Glyma.06G143000.1.p | chalcone isomerase 4A [Glycine max] | 99 | 100 | 2e-147 |
| | glyma_Glyma.11G004200.1.p | 2-hydroxyisoflavanone dehydratase [Glycine soja] | 97 | 97.20 | 0 |
| | glyma_Glyma.19G126000.1.p | cytochrome P450 98A2 [Glycine max] | 99 | 100 | 0 |
| | gosra_Gorai.009G243600.1 | flavonol synthase/flavanone 3-hydroxylase-like [Gossypium raimondii] | 99 | 100 | 0 |
| | gosra_Gorai.009G254800.1 | sterol 3-beta-glucosyltransferase UGT80B1 isoform X1 [Gossypium raimondii] | 99 | 100 | 0 |
| | jatcu_JCDBP15895 | flavonol synthase/flavanone 3-hydroxylase [Jatropha curcas] | 100 | 100 | 0 |
| | lepme_evm.model.scaffold98.478 | flavone synthase [Arabidopsis thaliana] | 100 | 95.24 | 0 |
| | lepme_evm.model.scaffold98.544 | flavonoid 3'-monooxygenase [Capsella rubella] | 97 | 95.65 | 3e-101 |
| | lepme_evm.model.scaffold98.739 | flavonoid 3'-monooxygenase [Capsella rubella] | 100 | 88.74 | 0 |
| | lotja_Lj1g3v0775780.1 | flavonol synthase/flavanone 3-hydroxylase [Medicago truncatula] | 100 | 84.64 | 8e-172 |
| | lotja_Lj1g3v4081860.1 | p-coumaroyl-shikimate 3'-hydroxylase [Trifolium pratense] | 100 | 91.94 | 0 |
| | lupan_XP_019421751.1 | 2-hydroxyisoflavanone dehydratase-like [Lupinus angustifolius] | 100 | 100 | 0 |
| | lupan_XP_019422242.1 | 2-hydroxyisoflavanone dehydratase-like [Lupinus angustifolius] | 100 | 100 | 0 |
| | manes_Manes.18G074900.1.p | flavonol synthase/flavanone 3-hydroxylase-like [Manihot esculenta] | 99 | 100 | 0 |
| | manes_Manes.18G092800.1.p | sterol 3-beta-glucosyltransferase UGT80B1 isoform X1 [Manihot esculenta] | 99 | 100 | 0 |
| | medtr_XP_003602395.1 | DOWNY MILDEW RESISTANCE 6 [Medicago truncatula] | 100 | 100 | 0 |
| | medtr_XP_003610679.1 | putative carboxylesterase, 2-hydroxyisoflavanone dehydratase [Medicago truncatula] | 100 | 100 | 0 |
| | medtr_XP_003610683.1 | 2-hydroxyisoflavanone dehydratase [Medicago truncatula] | 100 | 100 | 0 |
| | medtr_XP_003610685.1 | 2-hydroxyisoflavanone dehydratase [Medicago truncatula] | 100 | 100 | 0 |
| | medtr_XP_003612166.1 | naringenin 3-dioxygenase (flavanone-3-hydroxylase) [Medicago truncatula] | 100 | 100 | 0 |
| | medtr_XP_003623716.1 | vestitone reductase isoform X2 [Medicago truncatula] | 100 | 100 | 0 |
| | medtr_XP_003623718.2 | dihydroflavonol reductase [Medicago truncatula] | 100 | 100 | 0 |
| | medtr_XP_003623719.1 | vestitone reductase [Medicago truncatula] | 100 | 100 | 0 |
| | medtr_XP_003625205.1 | cytochrome P450 93A3 [Medicago truncatula] | 100 | 100 | 0 |
| | medtr_XP_003625207.1 | cytochrome P450 93A3 [Medicago truncatula] | 100 | 100 | 0 |
| | medtr_XP_013461509.1 | chalcone-flavanone isomerase family protein [Medicago truncatula] | 100 | 100 | 6e-166 |
| | momch_XP_022141387.1 | flavonol synthase/flavanone 3-hydroxylase [Momordica charantia] | 100 | 100 | 0 |
| | orysa_LOC_OS04G01140 | cytochrome P450 93G1-like [Oryza sativa Japonica Group] | 100 | 100 | 0 |
| | orysa_LOC_OS04G09680 | probable inactive methyltransferase Os04g0175900 [Oryza sativa Japonica Group] | 88 | 77.62 | 2e-107 |
| | orysa_LOC_OS10G02490 | deoxymugineic acid synthase 1-D [Oryza sativa Japonica Group] | 100 | 100 | 0 |
| | orysa_LOC_OS10G16974 | flavonoid 3'-monooxygenase CYP75B4-like [Oryza sativa Japonica Group] | 100 | 100 | 0 |
| | orysa_LOC_OS10G17260 | flavonoid 3'-monooxygenase CYP75B3-like [Oryza sativa Japonica Group] | 100 | 100 | 0 |
| | oryru_ORUFI10G05810.1 | flavonoid 3'-monooxygenase CYP75B4-like [Oryza sativa Japonica Group] | 100 | 99.81 | 0 |
| | oryru_ORUFI10G05920.1 | flavonoid 3'-monooxygenase CYP75B3-like [Oryza sativa Japonica Group] | 100 | 99.81 | 0 |
| | papso_XP_026451891.1 | uncharacterized protein LOC113352262 [Papaver somniferum] | 100 | 100 | 0 |
| | phavu_ESW29153 | 2-hydroxyisoflavanone dehydratase-like [Vigna angularis] | 88 | 92.24 | 0 |
| | pyrbr_Pbr020886.1 | cytochrome P450 98A2-like [Pyrus x bretschneideri] | 99 | 100 | 0 |
| | pyrbr_Pbr020888.1 | cytochrome P450 98A2-like [Pyrus x bretschneideri] | 99 | 100 | 0 |
| | pyrbr_Pbr020890.1 | cytochrome P450 98A2 isoform X1 [Pyrus x bretschneideri] | 96 | 94.38 | 0 |
| | ruboc_BRAS_G01409 | flavonol synthase [Fragaria x ananassa] | 100 | 93.73 | 0 |
| | selmo_Smo12440\|PACid_15405409 | flavonoid 3'-monooxygenase [Selaginella moellendorffii] | 100 | 100 | 0 |
| | selmo_Smo85404\|PACid_15419886 | flavonoid 3'-monooxygenase [Selaginella moellendorffii] | 99 | 98.80 | 0 |
| | solpe_Sopen05G030780.1 | probable chalcone--flavonone isomerase 3 [Solanum pennellii] | 100 | 100 | 3e-147 |
| | soltu_PGSC0003DMG400011655 | probable chalcone--flavonone isomerase 3 [Solanum tuberosum] | 99 | 100 | 4e-152 |
| | solyc_Solyc05G052240.3.1 | chalcone--flavonone isomerase 3 [Solanum lycopersicum] | 99 | 100 | 2e-151 |
| | thepa_Tp6g26120 | ADP-glucose phosphorylase [Eutrema salsugineum] | 100 | 89.68 | 0 |
| | thepa_Tp6g27110 | UDP-glycosyltransferase 78D2 [Eutrema salsugineum] | 99 | 85.93 | 0 |
| | thepa_Tp6g34220 | flavonol synthase/flavanone 3-hydroxylase [Eutrema salsugineum] | 100 | 94.64 | 0 |
| | thepa_Tp6g34960 | flavonoid 3'-monooxygenase [Eutrema salsugineum] | 99 | 92.59 | 0 |
| | thepa_Tp6g37260 | probable 2-oxoglutarate-dependent dioxygenase At5g05600 [Eutrema salsugineum] | 100 | 94.37 | 0 |

| | | | | | |
|---|---|---|---|---|---|
| | thepa_Tp6g37560 | probable chalcone--flavonone isomerase 3 [Raphanus sativus] | 100 | 94.15 | 2e-127 |
| | tripr_Tp57577_TGAC_v2_mRNA39268 | putative carboxylesterase 2-like protein [Trifolium pratense] | 99 | 99.70 | 0 |
| | tripr_Tp57577_TGAC_v2_mRNA39273 | putative carboxylesterase 2-like protein [Trifolium pratense] | 99 | 100 | 0 |
| **regulation of anthocyanin catabolic process** | araly_AL6G15770.t1 | UDP-glycosyltransferase 76C2 [Arabidopsis lyrata subsp. lyrata] | 100 | 100 | 0 |
| | araly_AL6G15780.t1 | UDP-glycosyltransferase 76C1 [Arabidopsis lyrata subsp. lyrata] | 100 | 100 | 0 |
| | araly_AL6G15810.t1 | UDP-glycosyltransferase 76C5 [Arabidopsis lyrata subsp. lyrata] | 100 | 100 | 0 |
| | araly_AL6G15820.t1 | UDP-glycosyltransferase 76C3 [Arabidopsis lyrata subsp. lyrata] | 99 | 100 | 0 |
| | arath_AT5G05860 | UDP-glucosyl transferase 76C2 [Arabidopsis thaliana] | 99 | 100 | 0 |
| | arath_AT5G05870 | UDP-glucosyl transferase 76C1 [Arabidopsis thaliana] | 99 | 100 | 0 |
| | arath_AT5G05880 | UDP-Glycosyltransferase superfamily protein [Arabidopsis thaliana] | 99 | 100 | 0 |
| | arath_AT5G05890 | UDP-Glycosyltransferase superfamily protein [Arabidopsis thaliana] | 99 | 100 | 0 |
| | arath_AT5G05900 | UDP-Glycosyltransferase superfamily protein [Arabidopsis thaliana] | 99 | 100 | 0 |
| | braol_BO3G003530.1 | UDP-glycosyltransferase 76C4-like [Brassica oleracea var. oleracea] | 100 | 100 | 0 |
| | camsa_XP_010452495.1 | UDP-glycosyltransferase 76C2 [Camelina sativa] | 100 | 100 | 0 |
| | camsa_XP_010491132.1 | UDP-glycosyltransferase 76C1-like [Camelina sativa] | 100 | 100 | 0 |
| | camsa_XP_010491133.1 | UDP-glycosyltransferase 76C2-like [Camelina sativa] | 100 | 100 | 0 |
| | camsa_XP_010495081.2 | UDP-glycosyltransferase 76C5-like [Camelina sativa] | 100 | 100 | 0 |
| | camsa_XP_010495082.1 | UDP-glycosyltransferase 76C3 [Camelina sativa] | 100 | 100 | 0 |
| | eutsa_Thhalv10013500m | UDP-glycosyltransferase 76C1 [Eutrema salsugineum] | 99 | 100 | 0 |
| | eutsa_Thhalv10013508m | UDP-glycosyltransferase 76C3 isoform X2 [Eutrema salsugineum] | 99 | 100 | 0 |
| | eutsa_Thhalv10013513m | UDP-glycosyltransferase 76C1 [Eutrema salsugineum] | 99 | 100 | 0 |
| | eutsa_Thhalv10015913m | UDP-glycosyltransferase 76C4 [Eutrema salsugineum] | 99 | 100 | 0 |
| | lepme_evm.model.scaffold98.712 | UDP-glycosyltransferase 76C3 isoform X2 [Eutrema salsugineum] | 99 | 74.34 | 0 |
| | lepme_evm.model.scaffold98.720 | UDP-glycosyltransferase 76C2 [Capsella rubella] | 100 | 86.18 | 0 |
| | lepme_evm.model.scaffold98.719 | UDP-glycosyltransferase 76C1 [Camelina sativa] | 99 | 82.97 | 0 |
| | thepa_Tp6g37010 | UDP-glycosyltransferase 76C5-like [Brassica oleracea var. oleracea] | 100 | 81.58 | 0 |
| | thepa_Tp6g37020 | UDP-glycosyltransferase 76C3 isoform X2 [Eutrema salsugineum] | 100 | 85.24 | 0 |
| | thepa_Tp6g37030 | UDP-glycosyltransferase 76C1 [Camelina sativa] | 100 | 83.15 | 0 |
| | thepa_Tp6g37040 | UDP-glycosyltransferase 76C2 [Eutrema salsugineum] | 100 | 92.05 | 0 |
| **regulation of flavonol biosynthetic process** | araly_AL1G36170.t1 | transcription factor MYB3 [Arabidopsis lyrata subsp. lyrata] | 100 | 100 | 0 |
| | araly_AL6G28270.t1 | glutathione S-transferase F12 [Arabidopsis lyrata subsp. lyrata]/TT19 | 100 | 100 | 9e-159 |
| | araly_AL7G16230.t1 | transcription factor MYB32 [Arabidopsis lyrata subsp. lyrata] | 100 | 100 | 0 |
| | arath_AT4G34990 | myb domain protein 32 [Arabidopsis thaliana] | 99 | 100 | 0 |
| | arath_AT5G17220 | glutathione S-transferase phi 12 [Arabidopsis thaliana]/TT19 | 99 | 100 | 1e-158 |
| | brana_GSBRNA2T00114880001 | glutathione S-transferase F12 [Brassica napus] | 96 | 97.58 | 1e-150 |
| | brara_Brara.A00330.1.p | transcription factor MYB32-like [Brassica rapa] | 99 | 100 | 0 |
| | brara_Brara.B00698.1.p | glutathione S-transferase F12 [Brassica napus] | 99 | 100 | 5e-159 |
| | brara_Brara.J01826.1.p | glutathione S-transferase F12-like [Brassica rapa] | 99 | 99.53 | 3e-156 |
| | camsa_XP_010420427.1 | glutathione S-transferase F12 [Camelina sativa] | 100 | 100 | 2e-158 |
| | camsa_XP_010453905.1 | glutathione S-transferase F12-like [Camelina sativa] | 100 | 100 | 2e-158 |
| | camsa_XP_010463766.1 | glutathione S-transferase F11-like isoform X1 [Camelina sativa] | 100 | 100 | 0 |
| | camsa_XP_010492664.1 | glutathione S-transferase F12 [Camelina sativa] | 100 | 100 | 1e-158 |
| | camsa_XP_010507366.1 | glutathione S-transferase F11 isoform X2 [Camelina sativa] | 100 | 100 | 1e-156 |
| | capgr_Cagra.2350S0045.1.p | transcription factor MYB32 [Capsella rubella] | 99 | 99.65 | 0 |
| | capru_Carubv10005438m | transcription factor MYB32 [Capsella rubella] | 99 | 100 | 0 |
| | capru_Carubv10014371m | transcription factor MYB7 [Capsella rubella] | 99 | 100 | 0 |
| | capru_Carubv10014608m | glutathione S-transferase F11 [Capsella rubella] | 99 | 100 | 0 |
| | citcl_XP_006434293.1 | glutathione S-transferase TCHQD [Citrus clementina] | 100 | 100 | 0 |
| | citla_Cla015834 | myb-related protein 308 [Cucumis sativus] | 100 | 93.23 | 2e-170 |
| | citsi_Cs3g20220.1 | glutathione S-transferase TCHQD [Citrus sinensis] | 99 | 100 | 0 |
| | cucme_MELO3C010516P1 | glutathione S-transferase TCHQD [Cucumis melo] | 99 | 100 | 0 |
| | cucsa_XP_004142639.1 | glutathione S-transferase TCHQD [Cucumis sativus] | 100 | 100 | 0 |
| | eutsa_Thhalv10008456m | transcription factor MYB3 [Eutrema salsugineum] | 99 | 100 | 0 |
| | eutsa_Thhalv10014654m | glutathione S-transferase F12 [Eutrema salsugineum] | 99 | 100 | 2e-158 |
| | eutsa_Thhalv10025926m | transcription factor MYB32 [Eutrema salsugineum] | 99 | 100 | 0 |
| | lepme_evm.model.scaffold70.605 | transcription factor MYB32 [Eutrema salsugineum] | 98 | 79.51 | 1e-150 |
| | lepme_evm.model.scaffold236.1050 | transcription factor MYB32 [Eutrema salsugineum] | 99 | 80.57 | 7e-154 |
| | glyma_Glyma.06G117800.2.p | glutathione S-transferase TCHQD-like [Glycine max] | 99 | 100 | 0 |
| | glyma_Glyma.06G160500.1.p | MYB transcription factor MYB56 [Glycine max] | 99 | 100 | 0 |
| | medtr_XP_013461627.1 | myb-related protein 308 [Medicago truncatula] | 100 | 100 | 0 |
| | ruboc_BRAS_G01502 | glutathione S-transferase TCHQD [Rosa chinensis] | 100 | 94.38 | 0 |
| | thepa_Tp1g20150 | transcription factor MYB3 [Brassica napus] | 100 | 86.57 | 9e-173 |

| | | | | | |
|---|---|---|---|---|---|
| | thepa_Tp6g27010 | glutathione S-transferase F12 [Brassica napus] | 99 | 90.61 | 1e-143 |
| | thepa_Tp7g32750 | transcription factor MYB32-like [Raphanus sativus] | 100 | 79.65 | 3e-151 |
| | zosma_Zosma3G00870.1 | Transcription factor MYB32 [Zostera marina] | 99 | 100 | 5e-143 |

Table S8: Enriched GO terms of category ‘Biological process’ directly linking to flavonoid metabolism using the set with the background ‘all genes’. PKS genes were removed prior to the enrichment analysis. The genes are listed according to their enriched processes ‘isoflavonoid metabolic process’ and ‘regulation of flavonol biosynthetic process’.

| | Gene | Description | Query cover [%] | Identity [%] | E value |
|---|---|---|---|---|---|
| **isoflavonoid metabolic process** | aradu_XP_015964809.1 | 2-hydroxyisoflavanone dehydratase-like [Arachis duranensis] | 100 | 100 | 0 |
| | aradu_XP_015964810.1 | 2-hydroxyisoflavanone dehydratase [Arachis duranensis] | 100 | 100 | 0 |
| | aradu_XP_015964830.1 | 2-hydroxyisoflavanone dehydratase-like [Arachis duranensis] | 100 | 100 | 0 |
| | cicar_Ca_11884 | 2-hydroxyisoflavanone dehydratase-like [Cicer arietinum] | 100 | 100 | 0 |
| | cicar_Ca_11885 | 2-hydroxyisoflavanone dehydratase-like [Cicer arietinum] | 100 | 100 | 0 |
| | eucgr_Eucgr.H03555.1.p | 2-hydroxyisoflavanone dehydratase isoform X1 [Eucalyptus grandis] | 85 | 100 | 0 |
| | glyma_Glyma.11G004200.1.p | 2-hydroxyisoflavanone dehydratase [Glycine soja] | 97 | 97.20 | 0 |
| | lupan_XP_019421751.1 | 2-hydroxyisoflavanone dehydratase-like [Lupinus angustifolius] | 100 | 100 | 0 |
| | lupan_XP_019422242.1 | 2-hydroxyisoflavanone dehydratase-like [Lupinus angustifolius] | 100 | 100 | 0 |
| | medtr_XP_003610679.1 | putative carboxylesterase, 2-hydroxyisoflavanone dehydratase [Medicago truncatula] | 100 | 100 | 0 |
| | orysa_LOC_OS10G12050 | Cytochrome P450 [Oryza sativa Japonica Group] | 17 | 100 | 1e-13 |
| | papso_XP_026449092.1 | (RS)-norcoclaurine 6-O-methyltransferase-like [Papaver somniferum] | 100 | 100 | 0 |
| | phavu_ESW29153 | 2-hydroxyisoflavanone dehydratase-like [Vigna angularis] | 88 | 92.24 | 0 |
| | solpe_Sopen01G050800.1 | 2-hydroxyisoflavanone dehydratase-like [Solanum pennellii] | 100 | 100 | 0 |
| | soltu_PGSC0003DMG400040651 | 2-hydroxyisoflavanone dehydratase-like [Solanum tuberosum] | 99 | 100 | 0 |
| | tripr_Tp57577_TGAC_v2_mRNA39273 | putative carboxylesterase 2-like protein [Trifolium pratense] | 99 | 100 | 0 |
| **regulation of flavonol biosynthetic process** | araly_AL1G36170.t1 | transcription factor MYB3 [Arabidopsis lyrata subsp. lyrata] | 100 | 100 | 0 |
| | araly_AL6G28270.t1 | glutathione S-transferase F12 [Arabidopsis lyrata subsp. lyrata]/TT19 | 100 | 100 | 9e-159 |
| | araly_AL7G16230.t1 | transcription factor MYB32 [Arabidopsis lyrata subsp. lyrata] | 100 | 100 | 0 |
| | arath_AT4G34990 | myb domain protein 32 [Arabidopsis thaliana] | 99 | 100 | 0 |
| | arath_AT5G17220 | glutathione S-transferase phi 12 [Arabidopsis thaliana]/TT19 | 99 | 100 | 1e-158 |
| | brana_GSBRNA2T00114880001 | glutathione S-transferase F12 [Brassica napus] | 96 | 97.58 | 1e-150 |
| | brara_Brara.A00330.1.p | transcription factor MYB32-like [Brassica rapa] | 99 | 100 | 0 |
| | brara_Brara.B00698.1.p | glutathione S-transferase F12 [Brassica napus] | 99 | 100 | 5e-159 |
| | brara_Brara.J01826.1.p | glutathione S-transferase F12-like [Brassica rapa] | 99 | 99.53 | 3e-156 |
| | camsa_XP_010420427.1 | glutathione S-transferase F12 [Camelina sativa] | 100 | 100 | 2e-158 |
| | camsa_XP_010453905.1 | glutathione S-transferase F12-like [Camelina sativa] | 100 | 100 | 2e-158 |
| | camsa_XP_010463766.1 | glutathione S-transferase F11-like isoform X1 [Camelina sativa] | 100 | 100 | 0 |
| | camsa_XP_010492664.1 | glutathione S-transferase F12 [Camelina sativa] | 100 | 100 | 1e-158 |
| | camsa_XP_010507366.1 | glutathione S-transferase F11 isoform X2 [Camelina sativa] | 100 | 100 | 1e-156 |
| | capgr_Cagra.2350S0045.1.p | transcription factor MYB32 [Capsella rubella] | 99 | 99.65 | 0 |
| | capru_Carubv10005438m | transcription factor MYB32 [Capsella rubella] | 99 | 100 | 0 |
| | capru_Carubv10014371m | transcription factor MYB7 [Capsella rubella] | 99 | 100 | 0 |
| | capru_Carubv10014608m | glutathione S-transferase F11 [Capsella rubella] | 99 | 100 | 0 |
| | citcl_XP_006434293.1 | glutathione S-transferase TCHQD [Citrus clementina] | 100 | 100 | 0 |
| | citla_Cla015834 | myb-related protein 308 [Cucumis sativus] | 100 | 93.23 | 2e-170 |
| | citsi_Cs3g20220.1 | glutathione S-transferase TCHQD [Citrus sinensis] | 99 | 100 | 0 |
| | cucme_MELO3C010516P1 | glutathione S-transferase TCHQD [Cucumis melo] | 99 | 100 | 0 |
| | cucsa_XP_004142639.1 | glutathione S-transferase TCHQD [Cucumis sativus] | 100 | 100 | 0 |
| | eutsa_Thhalv10008456m | transcription factor MYB3 [Eutrema salsugineum] | 99 | 100 | 0 |
| | eutsa_Thhalv10014654m | glutathione S-transferase F12 [Eutrema salsugineum] | 99 | 100 | 2e-158 |
| | eutsa_Thhalv10025926m | transcription factor MYB32 [Eutrema salsugineum] | 99 | 100 | 0 |
| | glyma_Glyma.06G117800.2.p | glutathione S-transferase TCHQD-like [Glycine max] | 99 | 100 | 0 |
| | glyma_Glyma.06G160500.1.p | MYB transcription factor MYB56 [Glycine max] | 99 | 100 | 0 |
| | lepme_evm.model.scaffold70.605 | transcription factor MYB32 [Eutrema salsugineum] | 98 | 79.51 | 1e-150 |
| | lepme_evm.model.scaffold236.1050 | transcription factor MYB32 [Eutrema salsugineum] | 99 | 80.57 | 7e-154 |
| | medtr_XP_013461627.1 | myb-related protein 308 [Medicago truncatula] | 100 | 100 | 0 |
| | ruboc_BRAS_G01502 | glutathione S-transferase TCHQD [Rosa chinensis] | 100 | 94.38 | 0 |
| | thepa_Tp1g20150 | transcription factor MYB3 [Brassica napus] | 100 | 86.57 | 9e-173 |
| | thepa_Tp6g27010 | glutathione S-transferase F12 [Brassica napus] | 99 | 90.61 | 1e-143 |
| | thepa_Tp7g32750 | transcription factor MYB32-like [Raphanus sativus] | 100 | 79.65 | 3e-151 |
| | zosma_Zosma3G00870.1 | Transcription factor MYB32 [Zostera marina] | 99 | 100 | 5e-143 |

#### Supplementary Text

##### PKS copy numbers widely vary among different plant and green algae species

Looking at the species-specific distribution (Supplementary Figure 2), we found differences in the number of type III PKS copies among the analyzed species. While we detected PKS protein signatures in all land plant species and various algae [e.g. *Aureococcus aneophagefferens* (Pelagomonodales, Pelagophyta), *Ostreococcus lucimarinus* (Mamiellalles, Chlorophyta) and *Coccomyxa sp. C169* (Chlorococcales, Chlorophyta)], the protein sequences were either shorter (< 300 amino acids) or longer (>550 amino acids) than the typical length of type III PKS proteins (380-430 amino acids) (with the exception of the three *Ectocarpus siliculosus*, Ectocarpales, Phaeophyta, sequences, with 399, 414 and 419 amino acids). Too short sequences will probably result in non-functional proteins due to lacking folds/active sites, though we cannot rule out that a misassembly of the genome may contribute to some of them. Too long sequences are probably β-ketoacyl ACP synthases that show similarity to type III PKS (the catalytic capability was not tested here). Genomes of species belonging to Chlorophyta, except the afore-mentioned, and Charophyta, which were available at the time of conducting the analysis, did not have any type III PKS (Supplementary Table S2). All analyzed vascular plants showed more than two copies of type III PKS. One clade within the Brassicales showed with *Capsella rubella* the lowest number of type III PKS in its proteome (*n*=2/2, OrthoFinder/MCL, *Arabidopsis thaliana*: *n*=4/4, AT1G02050/LAP6, AT4G00040, AT4G34850/LAP5, AT5G13930/TT4). Clades with a high number of type III PKSs (≥ 200 amino acids, Supplementary Table S1) were Fabales (12/12 to 33/33, Supplementary Figure 2), members of the BEP and PACMAD clade of the

Poaceae (10/10 *to* 50/54, Supplementary Figure 2*)*, members of gymnosperms (11/11 to 42/44, Supplementary Figure 2) and members of the Marchantiales (*Marchantia polymorpha*) and the Funariales (*Physcomitrella patens*).

Recently, the full genomes of two members of the Mesostigmaphyceae and Chlorokybophyceae of the Charophyta were released[90]. To test for the presence of type III PKS in these species, we blasted the CDS sequences of the eleven type III PKS homologs of *Penium margaritaceum*[91] against the whole genome shotgun sequences of *Mesostigma viride* and *Chlorokybus atmophyticus*. This analysis resulted in few hits with a maximum query cover of 2% for *Mesostigma viride* and 16% for *Chlorokybus atmophyticus*. We also queried the CDS sequences of *Penium margaritaceum* against all known sequences of the Coleochaetaphyceae yielding hits with ≤ 13% query cover (at the current moment, no full genome of a member belonging to the Coleochaetaphyceae is available). This indicates that there are no type III PKS sequences in the reported genomes of the Chlorokybophyceae Mesostigmaphyceae or Coleochaetaphyceae. The software tool pPAP[92] was used to accurately classify unknown type III PKS sequences (for a more detailed description of reaction types refer to Fig. 1 in the main text). The pPAP analysis of *Penium* PKS sequences yielded functional diversification (one of 'R-4-A', two of type 'R-4-C', eight of 'Other), however, some of the *Penium* sequences did not fall within the range of generic type III PKS sequences and could represent non-functional genes or β-ketoacyl ACP synthases. All of the *Penium* sequences were located at the base of the tree before the divergence in the LAP and CHS clade (Fig. 3) indicating that only with the conquest of the terrestrial space did type III PKS sequences diversify into the two major clades of the tree.

Other, more basal, algal taxa possess type I PKS, type II KS proteins and NRPs: Shelest et al.[93] studied the distribution of type I PKS, type II KS proteins and NRPs in algae for *Chlamydomonas reinhardtii* ($n_{type\ I\ PKS}$=1, $n_{type\ II\ KS}$=4, $n_{NRP}$=0), *Coccomyxa subellipsoidea* (10, 3, 1), *Ostreococcus lucimarinus* (3, 3, 0), *Volvox carteri* (1, 3, 0; the four former from Chlorophyta), *Cyanophora paradoxa* (0, 0, 1; Glaucophyta), *Aureococcus anophagefferens* (1, 56, 3), *Ectocarpus siliculosus* (1, 5, 0; the two former from Heterokontophyta), *Porphyridium purpureum* (0, 0, 0), *Cyanidioschyzon merolae* (0, 2, 0; the two former from Rhodophyta) and *Klebsormidium flaccidum* (0, 3, 0; Streptophyta)[93], suggesting that these algal taxa possess type I PKS and type II KS (with the exception of *Porhyridium purpureum*) instead of type III PKS, which evolved with or after the conquest of terrestrial habitats.

Typically, the number of 'R-4-C'-type sequences is limited per species to few copies (e.g. one copy in *Arabidopsis thaliana*, two copies in *Solanum lycopersicum*, one copy in *Oryza sativa*, two copies in *Zea mays*), although their genomes underwent multiple duplication and/or triplication events[94,95]. This indicates that deletion of duplicated genes or genomic segments containing CHS sequences limited the number of CHS sequences in a species. Deletion of genes after whole-genome multiplication events was reported previously[96-100].

It is interesting to note, that for 'R-4-A'-type *PKS* referring to the stilbene synthase function we find massive tandem gene duplications in the genome regions (*Vitis vinifera*, *Arachis sp.*, Fig. 2), while other 'R-4-A'-type *PKS* sequences do not show gene duplication. Tandem duplication of 'R-4-A'-type *PKS* sequences of the type STS, as the main principle for the generation of STS genes, was also found for mulberry (*Morus spp.*) previously[101].

#### Evolution of valerophenone synthase in *Humulus lupulus* and olivetol synthase in *Cannabis sativa*

Next to our analysis of the evolution of chalcone synthase (CHS), acyl-CoA/hydroxyalkyl pyrone/less adhesive pollensynthase (LAP) and stilbene synthase (STS), we analyzed the evolution of valerophenone synthase (VPS) from *Humulus lupulus* and olivetol synthase (OLS) from *Cannabis sativa* (both species from the Cannabaceae family). VPS and OLS are classified according to pPAP as ‘Other’. The characterized VPS and OLS sequences (see the labels 13 and 23 in the Fig. 3) are located in a separate clade from the ‘R-4-C’-type sequences of *Humulus lupulus* and *Cannabis sativa* in the phylogenetic tree.

However, also the CHS-type sequences that correspond to CHS_H1 in the study of Novak et al.[102] (Humlu_CAC19808, humlu_HL.SW.v1.0.G018947.1) utilize isovaleryl-CoA and isobutyryl-CoA as substrates, although primarily catalyzing the formation of naringenin chalcone. The two corresponding enzymes are annotated as ‘R-4-C’ by pPAP here. These two sequences are located in the clade containing ‘R-4-C’ sequences from *Cannabis sativa* and *Humulus lupulus*. The enzyme with the sequence corresponding to VPS (here: humlu_BAB12012) catalyzes the formation of naringenin chalcone, albeit at lower rates than that encoded by *CHS_H1* indicating that the sequences in the clade containing humlu_BAB12012 are VPSs and not CHSs. This suggests that VPS and OLS evolved, together with the other sequences of the type ‘Other’, before the speciation of the two species, forming a monophyletic clade (Fig. 3) since this clade is distinct from the clade containing the ‘R-4-C’-type sequences of the two species. Novak et al.[103] studied the evolution of CHS-like homologues and found that three type III PKS sequences (VPS, CHS3 and CHS4) including VPS are located in the same genomic region. The sequences correspond in this study to humlu_BAB12102, humlu_HL_SW_v1.0_G013555.1, humlu_HL_SW_v1.0_G038363.1 (VPS, type 'Other'), to humlu_BAB47196, humlu_HL_SW_v1.0_G005266.1 (CHS3, 'Other') and to humlu_CAD23044, humlu_ACM17226, humlu_HL_SW_v1.0_G013566.1 (CHS4, 'Other'). This indicates that the clade containing *Humulus lupulus* and *Cannabis sativa* sequences of the type 'Other' originated by gene tandem duplication and that the genomic region is distinct from the sequences of the 'R-4-C' type (CHS_H1 in Novak et al.[103] corresponding to humlu_CAC19808, humlu_CAK19319, humlu_CAK19318, humlu_ACM17224, humlu_HL_SW_v1.0_G018947.1, see Fig. 3).

**Evolution of aleosone, chromone and octaketide synthases in *Aloe arborescens***

*Aloe arborescens* has the enzymatic capability to synthesize (i) 5,7-dihydroxy-2-methylchromone, from five molecules of malonyl-CoA, by a chromone synthase[104], (ii) octaketides SEK4 and SEK4b, from seven molecules of malonyl-CoA, by an octaketide synthase[105,106] and (iii) aleosone, from seven molecules of malonyl-CoA, by an aleosone synthase[106]. The corresponding sequences (Aloar_AAT48709, Aloar_AAX35541, Aloar_ABS72373, Aloar_ACR19997, Aloar_ACR19998, see the labels 16, 17 and 18 in Fig. 3) form a monophyletic clade within the phylogenetic tree, indicating that they evolved from the same sequence, possibly by recent tandem duplication or segmental duplication (no full genome is currently available for *Aloe arborescens* to further test these hypotheses). Furthermore, the sequences locate to a clade that contains mainly sequences of the 'Other' type from monocots, indicating that these sequences are distinct from the 'R-4-C' type and that they share a similar macroevolutionary trajectory compared to monocot type III PKS sequences belonging to the syntenic clusters 4, 5, 10 and 22 (Fig. 3).

**_CHS_-containing syntenic regions lack flavonoid biosynthetic gene cluster**

To further scrutinize the possibility of flavonoid biosynthetic gene cluster formation within the syntenic regions containing *CHS* genes, we analyzed the co-expression of *CHS* genes with genes in the syntenic regions. To this end, we employed the STRING database to detect functional protein association networks and used the syntenic regions from the model species *Arabidopsis thaliana*, *Oryza sativa* and *Solanum lycopersicum* and *Vitis vinifera* as input. Specifically, co-expression analysis was conducted using previously characterized or annotated type III *PKS* (*AT5G13930/TT4*, *Os11g32650*, *Solyc05g053550/SlCHS2*, *Solyc09g091510/SlCHS1*, *Solyc12g098100*, *GSVIVT01032968001*; Supplementary Figure 10). Although co-expression networks were detected for all analyzed genes within the PKS containing regions, they did not show any co-expression with *CHS* except for *AT5G13930/TT4*, which is located in a very large syntenic region on chromosome 5 in *Arabidopsis thaliana* (799 genes were used for STRING database analysis). However, the two co-expressed genes, glutathione S-transferase phi 12 (*AT5G17220*) and UDP-glycosyl transferase 78D2 (*AT5G17050*), were not located in close vicinity to *AT5G13930/TT4*. Generally, we detected functional links using the co-expression that was in agreement with the GO enrichment analysis (e.g. genes that were related to translation and transcription), but the analysis was alleviated by poor annotation of the genes within the syntenic regions for the studied species. Taken together with the GO enrichment analysis, where we found only few terms linked to direct flavonoid biosynthetic process, that were enriched for mainly

Brassicales species that have long syntenic regions or only present in a subset of species, we conclude that there are no flavonoid-specific biosynthetic gene clusters in the *CHS*-containing regions.

It is important to note that for all Gymnosperm species, no information on the syntenic relationship could be obtained due to low scaffold quality of their assembled genomes. From the phylogenetic analysis it seems most probable that Gymnosperms should be located in the syntenic cluster 5 (containing members of Bryophyta, Marchantiophyta and Polypodiopsida) or syntenic cluster 4 (containing members of Lycopodiophyta and Pteridophyta).

**Interaction of chalcone synthase with chalcone reductase might drive macroevolution of chalcone synthases in the Fabales**

Intriguingly for the Fabales clade, the concerted biosynthesis of CHS and chalcone reductase (CHR) was reported to be involved in the biosynthesis of isoflavone that are required for the establishment of the symbiosis between root and associated bacteria[49,107-113]. Hereby, CHS interacts with CHR indirectly via 2-hydroxyisoflavonoe synthase isozymes in an isoflavone metabolon[107].

To our knowledge, only in *Glycine max* specific candidate CHS genes involved in isoflavone biosynthesis interacting with CHR were proposed (CHS1 and CHS7[107] *and* CHS7 and CHS8[110,113]). CHS1, CHS7 and CHS8 of *Glycine max* located in our study to syntenic cluster 5, the cluster showing 'R-4-C'-type sequences for all Fabales species (Supplementary Figure 5). It is an interesting hypothesis if other 'R-4-C'-type sequences in this syntenic cluster are also involved in the biosynthesis of isoflavones and if syntenic clusters contain specifically those 'R-4-C'-type sequences required for isoflavone biosynthesis indicating that a duplication event facilitated the biosynthesis of isoflavones in the Fabales.

**Multi-tier evolution of PKS: relaxed conservation of gene expression on *CHS* orthologs and conservation of expression on *LAP5/6* orthologs within syntenic clusters**

Besides the evolution changes in a genome-wide context and the divergence of gene sequences, gene expression might also alter, especially after duplication events[114,115]. To test the relationship between gene expression divergence and gene duplication, we evaluated if genes in the same syntenic network cluster entails similar expression patterns across tissues (Supplementary Figure 11) and if the expression of type III PKS genes tend to alter after duplication events. To this end, we obtained expression data of type III PKS from the model species *Arabidopsis thaliana*, *Oryza sativa*, *Selaginella moellendorffii*, *Solanum lycopersicum*, *Vitis vinifera* and *Zea mays* from the CoNekT database[116] and categorized the expression data by tissue across the six species. Based on their Pearson-correlation values (Supplementary Figure 11 A) and affinity propagation clustering, the expression data was partitioned into four expression clusters (I to IV) that are specific to different tissues (Supplementary Figure 11 C). To some extent, gene expression was conserved for the 'R-4-C'-type *PKS* (corresponding to *CHS*) as exemplified by the genes *AT5G13930*/*TT4* from *Arabidopsis thaliana*, *LOC_Os11g32650* from *Oryza sativa*, *Zm00001d007403*/*Whp1* and *Zm00001d052673*/*C2* from *Zea mays* that were members of expression cluster I showing high expression in flowers and members of the syntenic cluster 4 (Supplementary Figure 11 B). However, other 'R-4-C'-type

*PKS* detected in syntenic cluster 2 and 4 (Fig. 2) were found in expression cluster III having high expression in reproductive tissues and stem/shoot tissue (*Solyc05g053550.3.1/SlCHS2* and *Solyc09g091510.3.1/SlCHS1* from *Solanum lycopersicum*, *GSVIVT01032968001* from *Vitis vinifera*). Similarly, 'R-4-A'-type *PKS*, assigned to the syntenic cluster 5, were found in both the expression clusters II and III (Supplementary Figure 11 B). These results suggest that 'R-4-C' and 'R-4-A' type *PKS* from even the same syntenic cluster(s) can be expressed differently.

LAP5 and LAP6 homologues (*AT1G02050/LAP6*, *AT4G00040*, *AT4G34850/LAP5* from *Arabidopsis thaliana*, *Smo122361|PACid_15419808* and *Smo231846|PACid_15419824* from *Selaginella moellendorffii*, *Solyc01g090600.3.1* and *Solyc01g111070.3.1* from *Solanum lycopersicum*, *GSVIVT01018219001*, *GSVIVT01024107001* from *Vitis vinifera*, *Zm00001d013991*, *Zm00001d019478*, *Zm00001d032662* from *Zea mays*) showed high expression in flower, fruit/siliques/ear/strobilus/spores or pollen (Supplementary Figure 11, for *Selaginella moellendorffii* expression information is only available for fruit/siliques/ear/strobilus/spores and leaves, for *Vitis vinifera* no expression information is available for pollen and roots/rhizoids). These transcripts were found in the expression clusters I (*Arabidopsis thaliana*, *Solanum lycopersicum*, *Zea mays*), III (*Solanum lycopersicum*, *Vitis vinifera*) and IV (*Selaginella moellendorffii*) (Supplementary Figure 11). Expression values for flower and fruit/siliques/ear/strobilus/spores were not available for *LOC_Os07g22850* and *LOC_Os10g34360* and the two transcripts from *Oryza sativa* showed highest expression in stem/shoot.

The syntenic cluster membership does not dictate the gene expression pattern for the CHS family members analyzed here. Given the occurrence of flavonoids in diverse plant tissues, this observation is likely because the genomic maintenance of syntenic region is overlaid by evolution of transcriptional regulation and that transcriptional adaptation is happening faster than the loss of synteny of genomic regions. Similarly in *Nicotiana tabacum*, in nicotine biosynthesis the non-syntenic *quinolate phosphoribosyltransferase* (*QPT*) *1* and *QPT2*, which evolved by gene duplication in a *Nicotiana* ancestor and have a sequence identity of 94%, exhibit different expression profiles and different response to stress stimuli due to *cis*-regulatory divergence[117,118].

The expression of *LAP5* and *LAP6* orthologs coincided with the expected gene expression pattern in all species where expression data was available for all necessary tissues. *LAP5* and *LAP6* orthologs showed expression in flower, fruit/siliques/ear/strobilus/spores or pollen that overlaps with the localization of sporopollenin biosynthesis. *LAP5* homologs locate to the syntenic clusters 1 and 27, while *LAP6* homologs locate to the syntenic clusters 3 and 11 (cf. Fig. 2 and Supplementary Table S3). LAP5 and LAP6 form two distinct clades in the phylogenetic tree (Fig. 3), however, the clades do not show inter-cluster synteny. *LAP5/6* confer a tissue-specific role in the biosynthesis of sporopollenin, conserving its expression over different syntenic clusters.

1 Emms, D. M. & Kelly, S. OrthoFinder: phylogenetic orthology inference for comparative genomics. *Genome Biol* **20**, 238, doi:10.1186/s13059-019-1832-y (2019).

2 Lemoine, F. *et al.* Renewing Felsenstein's phylogenetic bootstrap in the era of big data. *Nature* **556**, 452-+, doi:10.1038/s41586-018-0043-0 (2018).

3 Kim, S. S. *et al.* LAP6/POLYKETIDE SYNTHASE A and LAP5/POLYKETIDE SYNTHASE B Encode Hydroxyalkyl alpha-Pyrone Synthases Required for Pollen Development and Sporopollenin Biosynthesis in Arabidopsis thaliana. *Plant Cell* **22**, 4045-4066, doi:10.1105/tpc.110.080028 (2010).

4 Dobritsa, A. A. *et al.* LAP5 and LAP6 Encode Anther-Specific Proteins with Similarity to Chalcone Synthase Essential for Pollen Exine Development in Arabidopsis. *Plant Physiol* **153**, 937-955, doi:10.1104/pp.110.157446 (2010).

5 Koenig, D. *et al.* Comparative transcriptomics reveals patterns of selection in domesticated and wild tomato. *P Natl Acad Sci USA* **110**, E2655-E2662, doi:10.1073/pnas.1309606110 (2013).

6 Besseau, S. *et al.* Flavonoid accumulation in Arabidopsis repressed in lignin synthesis affects auxin transport and plant growth. *Plant Cell* **19**, 148-162, doi:10.1105/tpc.106.044495 (2007).

7 Bohl, M., Czupalla, C., Tokalov, S. V., Hoflack, B. & Gutzeit, H. O. Identification of actin as quercetin-binding protein: An approach to identify target molecules for specific ligands. *Anal Biochem* **346**, 295-299, doi:10.1016/j.ab.2005.08.037 (2005).

8 Bohl, M. *et al.* Flavonoids affect actin functions in cytoplasm and nucleus. *Biophys J* **93**, 2767-2780, doi:10.1529/biophysj.107.107813 (2007).

9 McLusky, S. R. *et al.* Cell wall alterations and localized accumulation of feruloyl-3 '-methoxytyramine in onion epidermis at sites of attempted penetration by Botrytis allii are associated with actin polarisation, peroxidase activity and suppression of flavonoid biosynthesis. *Plant J* **17**, 523-534, doi:DOI 10.1046/j.1365-313X.1999.00403.x (1999).

10 Hsieh, K. & Huang, A. H. C. Tapetosomes in Brassica tapetum accumulate endoplasmic reticulum-derived flavonoids and alkanes for delivery to the pollen surface. *Plant Cell* **19**, 582-596, doi:10.1105/tpc.106.049049 (2007).

11 Thompson, E. P., Wilkins, C., Demidchik, V., Davies, J. M. & Glover, B. J. An Arabidopsis flavonoid transporter is required for anther dehiscence and pollen development. *J Exp Bot* **61**, 439-451, doi:10.1093/jxb/erp312 (2010).

12 Quilichini, T. D., Samuels, A. L. & Douglas, C. J. ABCG26-Mediated Polyketide Trafficking and Hydroxycinnamoyl Spermidines Contribute to Pollen Wall Exine Formation in Arabidopsis. *Plant Cell* **26**, 4483-4498, doi:10.1105/tpc.114.130484 (2014).

13 Pearce, S., Ferguson, A., King, J. & Wilson, Z. A. FlowerNet: A Gene Expression Correlation Network for Anther and Pollen Development. *Plant Physiol* **167**, 1717-U1923, doi:10.1104/pp.114.253807 (2015).

14 Peer, W. A. & Murphy, A. S. Flavonoids and auxin transport: modulators or regulators? *Trends Plant Sci* **12**, 556-563, doi:10.1016/j.tplants.2007.10.003 (2007).

15 Ringli, C. *et al.* The modified flavonol glycosylation profile in the Arabidopsis rol1 mutants results in alterations in plant growth and cell shape formation. *Plant Cell* **20**, 1470-1481, doi:10.1105/tpc.107.053249 (2008).

16 Santelia, D. *et al.* Flavonoids redirect PIN-mediated polar auxin fluxes during root gravitropic responses. *J Biol Chem* **283**, 31218-31226, doi:10.1074/jbc.M710122200 (2008).

17 Tan, J. F. *et al.* A Genetic and Metabolic Analysis Revealed that Cotton Fiber Cell Development Was Retarded by Flavonoid Naringenin. *Plant Physiol* **162**, 86-95, doi:10.1104/pp.112.212142 (2013).

18 Mo, Y. Y., Nagel, C. & Taylor, L. P. Biochemical Complementation of Chalcone Synthase Mutants Defines a Role for Flavonols in Functional Pollen. *P Natl Acad Sci USA* **89**, 7213-7217, doi:DOI 10.1073/pnas.89.15.7213 (1992).

19 Zuk, M. *et al.* Chalcone Synthase (CHS) Gene Suppression in Flax Leads to Changes in Wall Synthesis and Sensing Genes, Cell Wall Chemistry and Stem Morphology Parameters. *Front Plant Sci* **7**, doi:ARTN 894 10.3389/fpls.2016.00894 (2016).
20 Lepikson-Neto, J. *et al.* Flavonoid supplementation affects the expression of genes involved in cell wall formation and lignification metabolism and increases sugar content and saccharification in the fast-growing eucalyptus hybrid E. urophylla x E. grandis. *Bmc Plant Biol* **14**, 301, doi:10.1186/s12870-014-0301-8 (2014).
21 Schulz, E., Tohge, T., Zuther, E., Fernie, A. R. & Hincha, D. K. Natural variation in flavonol and anthocyanin metabolism during cold acclimation in Arabidopsis thaliana accessions. *Plant Cell Environ* **38**, 1658-1672, doi:10.1111/pce.12518 (2015).
22 Schulz, E., Tohge, T., Zuther, E., Fernie, A. R. & Hincha, D. K. Flavonoids are determinants of freezing tolerance and cold acclimation in Arabidopsis thaliana. *Sci Rep* **6**, 34027, doi:10.1038/srep34027 (2016).
23 Yamasaki, H., Sakihama, Y. & Ikehara, N. Flavonoid-peroxidase reaction as a detoxification mechanism of plant cells against H2O2. *Plant Physiol* **115**, 1405-1412, doi:DOI 10.1104/pp.115.4.1405 (1997).
24 Park, S. S., Bae, I. & Lee, Y. J. Flavonoids-induced accumulation of hypoxia-inducible factor (HIF)-1 alpha/2 alpha is mediated through chelation of iron. *J Cell Biochem* **103**, 1989-1998, doi:10.1002/jcb.21588 (2008).
25 Loreti, E. *et al.* Gibberellins, jasmonate and abscisic acid modulate the sucrose-induced expression of anthocyanin biosynthetic genes in Arabidopsis. *New Phytol* **179**, 1004-1016, doi:10.1111/j.1469-8137.2008.02511.x (2008).
26 Lucho-Constantino, G. G. *et al.* Antioxidant responses under jasmonic acid elicitation comprise enhanced production of flavonoids and anthocyanins in Jatropha curcas leaves. *Acta Physiol Plant* **39**, doi:ARTN 165 10.1007/s11738-017-2461-2 (2017).
27 Lu, Y. F. *et al.* Flavonoid Accumulation Plays an Important Role in the Rust Resistance of Malus Plant Leaves. *Front Plant Sci* **8**, doi:ARTN 1286 10.3389/fpls.2017.01286 (2017).
28 Nikiforova, V. *et al.* Transcriptome analysis of sulfur depletion in Arabidopsis thaliana: interlacing of biosynthetic pathways provides response specificity. *Plant J* **33**, 633-650, doi:DOI 10.1046/j.1365-313X.2003.01657.x (2003).
29 Jackson, T. L. *et al.* Large Cellular Inclusions Accumulate in Arabidopsis Roots Exposed to Low-Sulfur Conditions. *Plant Physiol* **168**, 1573-U1857, doi:10.1104/pp.15.00465 (2015).
30 Nakabayashi, R., Mori, T. & Saito, K. Alternation of flavonoid accumulation under drought stress in Arabidopsis thaliana. *Plant Signal Behav* **9**, e29518, doi:10.4161/psb.29518 (2014).
31 Nakabayashi, R. *et al.* Enhancement of oxidative and drought tolerance in Arabidopsis by overaccumulation of antioxidant flavonoids. *Plant J* **77**, 367-379, doi:10.1111/tpj.12388 (2014).
32 Adato, A. *et al.* Fruit-Surface Flavonoid Accumulation in Tomato Is Controlled by a SlMYB12-Regulated Transcriptional Network. *Plos Genet* **5**, doi:ARTN e1000777 10.1371/journal.pgen.1000777 (2009).
33 Espana, L. *et al.* Transient Silencing of CHALCONE SYNTHASE during Fruit Ripening Modifies Tomato Epidermal Cells and Cuticle Properties. *Plant Physiol* **166**, 1371-+, doi:10.1104/pp.114.246405 (2014).
34 Heredia, A., Heredia-Guerrero, J. A. & Dominguez, E. CHS silencing suggests a negative cross-talk between wax and flavonoid pathways in tomato fruit cuticle. *Plant Signal Behav* **10**, doi:10.1080/15592324.2015.1019979 (2015).
35 Ng, J. L. P. *et al.* Flavonoids and Auxin Transport Inhibitors Rescue Symbiotic Nodulation in the Medicago truncatula Cytokinin Perception Mutant cre1. *Plant Cell* **27**, 2210-2226, doi:10.1105/tpc.15.00231 (2015).

36 Toffolatti, S. L., Venturini, G., Maffi, D. & Vercesi, A. Phenotypic and histochemical traits of the interaction between Plasmopara viticola and resistant or susceptible grapevine varieties. *Bmc Plant Biol* **12**, doi:Artn 124
10.1186/1471-2229-12-124 (2012).

37 Blount, J. W., Dixon, R. A. & Paiva, N. L. Stress Responses in Alfalfa (Medicago-Sativa L) .16. Antifungal Activity of Medicarpin and Its Biosynthetic Precursors - Implications for the Genetic Manipulation of Stress Metabolites. *Physiol Mol Plant P* **41**, 333-349, doi:Doi 10.1016/0885-5765(92)90020-V (1992).

38 Hain, R. *et al.* Disease Resistance Results from Foreign Phytoalexin Expression in a Novel Plant. *Nature* **361**, 153-156, doi:DOI 10.1038/361153a0 (1993).

39 Leckband, G. & Lorz, H. Transformation and expression of a stilbene synthase gene of Vitis vinifera L. in barley and wheat for increased fungal resistance. *Theor Appl Genet* **96**, 1004-1012, doi:DOI 10.1007/s001220050832 (1998).

40 Hipskind, J. D. & Paiva, N. L. Constitutive accumulation of a resveratrol-glucoside in transgenic alfalfa increases resistance to Phoma medicaginis. *Mol Plant Microbe In* **13**, 551-562, doi:Doi 10.1094/Mpmi.2000.13.5.551 (2000).

41 GrantPetersson, J. & Renwick, J. A. A. Effects of ultraviolet-B exposure of Arabidopsis thaliana on herbivory by two crucifer-feeding insects (Lepidoptera). *Environ Entomol* **25**, 135-142, doi:DOI 10.1093/ee/25.1.135 (1996).

42 Misra, P. *et al.* Modulation of Transcriptome and Metabolome of Tobacco by Arabidopsis Transcription Factor, AtMYB12, Leads to Insect Resistance. *Plant Physiol* **152**, 2258-2268, doi:10.1104/pp.109.150979 (2010).

43 Onkokesung, N. *et al.* Modulation of flavonoid metabolites in Arabidopsis thaliana through overexpression of the MYB75 transcription factor: role of kaempferol-3,7-dirhamnoside in resistance to the specialist insect herbivore Pieris brassicae. *J Exp Bot* **65**, 2203-2217, doi:10.1093/jxb/eru096 (2014).

44 Fernie, A. R. Evolution: An Early Role for Flavonoids in Defense against Oomycete Infection. *Curr Biol* **29**, R688-R690, doi:10.1016/j.cub.2019.06.028 (2019).

45 Brown, D. E. *et al.* Flavonoids act as negative regulators of auxin transport in vivo in Arabidopsis. *Plant Physiol* **126**, 524-535, doi:DOI 10.1104/pp.126.2.524 (2001).

46 Gao, Y. F. *et al.* Tomato SlAN11 regulates flavonoid biosynthesis and seed dormancy by interaction with bHLH proteins but not with MYB proteins. *Hortic Res-England* **5**, doi:ARTN 27
10.1038/s41438-018-0032-3 (2018).

47 Gu, X. Y. *et al.* Association Between Seed Dormancy and Pericarp Color Is Controlled by a Pleiotropic Gene That Regulates Abscisic Acid and Flavonoid Synthesis in Weedy Red Rice. *Genetics* **189**, 1515-+, doi:10.1534/genetics.111.131169 (2011).

48 Sotelo-Silveira, M. *et al.* Cytochrome P450 CYP78A9 Is Involved in Arabidopsis Reproductive Development. *Plant Physiol* **162**, 779-799, doi:10.1104/pp.113.218214 (2013).

49 Gupta, R. *et al.* A Multi-Omics Analysis of Glycine max Leaves Reveals Alteration in Flavonoid and Isoflavonoid Metabolism Upon Ethylene and Abscisic Acid Treatment. *Proteomics* **18**, doi:ARTN 1700366
10.1002/pmic.201700366 (2018).

50 Fasano, R. *et al.* Role of Arabidopsis UV RESISTANCE LOCUS 8 in Plant Growth Reduction under Osmotic Stress and Low Levels of UV-B. *Mol Plant* **7**, 773-791, doi:10.1093/mp/ssu002 (2014).

51 Buer, C. S. & Muday, G. K. The transparent testa4 mutation prevents flavonoid synthesis and alters auxin transport and the response of Arabidopsis roots to gravity and light. *Plant Cell* **16**, 1191-1205, doi:DOI 10.1105/tpc.020313 (2004).

52 Buer, C. S., Sukumar, P. & Muday, G. K. Ethylene modulates flavonoid accumulation and gravitropic responses in roots of Arabidopsis. *Plant Physiol* **140**, 1384-1396, doi:10.1104/pp.105.075671 (2006).

53 Yang, H. X. *et al.* Reduction of root flavonoid level and its potential involvement in lateral root emergence in Arabidopsis thaliana grown under low phosphate supply. *Funct Plant Biol* **36**, 564-573, doi:10.1071/Fp08283 (2009).
54 Maloney, G. S., DiNapoli, K. T. & Muday, G. K. The anthocyanin reduced Tomato Mutant Demonstrates the Role of Flavonols in Tomato Lateral Root and Root Hair Development. *Plant Physiol* **166**, 614-U254, doi:10.1104/pp.114.240507 (2014).
55 Thomas, H., Huang, L., Young, M. & Ougham, H. Evolution of plant senescence. *Bmc Evol Biol* **9**, doi:Artn 163
10.1186/1471-2148-9-163 (2009).
56 Liang, D. *et al.* Exogenous Melatonin Application Delays Senescence of Kiwifruit Leaves by Regulating the Antioxidant Capacity and Biosynthesis of Flavonoids. *Front Plant Sci* **9**, doi:ARTN 426
10.3389/fpls.2018.00426 (2018).
57 Chen, M. J. *et al.* System Analysis of an Arabidopsis Mutant Altered in de Novo Fatty Acid Synthesis Reveals Diverse Changes in Seed Composition and Metabolism. *Plant Physiol* **150**, 27-41, doi:10.1104/pp.108.134882 (2009).
58 Chen, M. X. *et al.* The Effect of TRANSPARENT TESTA2 on Seed Fatty Acid Biosynthesis and Tolerance to Environmental Stresses during Young Seedling Establishment in Arabidopsis. *Plant Physiol* **160**, 1023-1036, doi:10.1104/pp.112.202945 (2012).
59 Watkins, J. M., Chapman, J. M. & Muday, G. K. Abscisic Acid-Induced Reactive Oxygen Species Are Modulated by Flavonols to Control Stomata Aperture. *Plant Physiol* **175**, 1807-1825, doi:10.1104/pp.17.01010 (2017).
60 Koyama, R. *et al.* Exogenous Abscisic Acid Promotes Anthocyanin Biosynthesis and Increased Expression of Flavonoid Synthesis Genes in Vitis vinifera x Vitis labrusca Table Grapes in a Subtropical Region. *Front Plant Sci* **9**, doi:ARTN 323
10.3389/fpls.2018.00323 (2018).
61 Petridis, A., Doll, S., Nichelmann, L., Bilger, W. & Mock, H. P. Arabidopsis thaliana G2-LIKE FLAVONOID REGULATOR and BRASSINOSTEROID ENHANCED EXPRESSION1 are low-temperature regulators of flavonoid accumulation. *New Phytol* **211**, 912-925, doi:10.1111/nph.13986 (2016).
62 Li, X. *et al.* Nitric oxide mediates brassinosteroid-induced flavonoid biosynthesis in Camellia sinensis L. *J Plant Physiol* **214**, 145-151, doi:10.1016/j.jplph.2017.04.005 (2017).
63 Hinderer, W., Petersen, M. & Seitz, H. U. Inhibition of Flavonoid Biosynthesis by Gibberellic-Acid in Cell-Suspension Cultures of Daucus-Carota L. *Planta* **160**, 544-549, doi:Doi 10.1007/Bf00411143 (1984).
64 Ribeiro, D. M., Araujo, W. L., Fernie, A. R., Schippers, J. H. M. & Mueller-Roeber, B. Translatome and metabolome effects triggered by gibberellins during rosette growth in Arabidopsis. *J Exp Bot* **63**, 2769-2786, doi:10.1093/jxb/err463 (2012).
65 Kubasek, W. L. *et al.* Regulation of Flavonoid Biosynthetic Genes in Germinating Arabidopsis Seedlings. *Plant Cell* **4**, 1229-1236, doi:Doi 10.2307/3869409 (1992).
66 Davies, K. M., Bradley, J. M., Schwinn, K. E., Markham, K. R. & Podivinsky, E. Flavonoid Biosynthesis in Flower Petals of 5 Lines of Lisianthus (Eustoma-Grandiflorum Grise). *Plant Sci* **95**, 67-77, doi:Doi 10.1016/0168-9452(93)90080-J (1993).
67 Kazuma, K., Noda, N. & Suzuki, M. Flavonoid composition related to petal color in different lines of Clitoria ternatea. *Phytochemistry* **64**, 1133-1139, doi:10.1016/S0031-9422(03)00504-1 (2003).
68 Li, J. Y., Oulee, T. M., Raba, R., Amundson, R. G. & Last, R. L. Arabidopsis Flavonoid Mutants Are Hypersensitive to Uv-B Irradiation. *Plant Cell* **5**, 171-179, doi:DOI 10.1105/tpc.5.2.171 (1993).
69 Vanhaelewyn, L. *et al.* Differential UVR8 Signal across the Stem Controls UV-B-Induced Inflorescence Phototropism. *Plant Cell* **31**, 2070-2088, doi:10.1105/tpc.18.00929 (2019).

70 Coe, E. H., Mccormick, S. M. & Modena, S. A. White Pollen in Maize. *J Hered* **72**, 318-320, doi:DOI 10.1093/oxfordjournals.jhered.a109514 (1981).
71 Ylstra, B. *et al.* Flavonols Stimulate Development, Germination, and Tube Growth of Tobacco Pollen. *Plant Physiol* **100**, 902-907, doi:DOI 10.1104/pp.100.2.902 (1992).
72 Muhlemann, J. K., Younts, T. L. B. & Muday, G. K. Flavonols control pollen tube growth and integrity by regulating ROS homeostasis during high-temperature stress. *P Natl Acad Sci USA* **115**, E11188-E11197, doi:10.1073/pnas.1811492115 (2018).
73 French, C. J. & Towers, G. H. N. Inhibition of Infectivity of Potato-Virus X by Flavonoids. *Phytochemistry* **31**, 3017-3020, doi:Doi 10.1016/0031-9422(92)83438-5 (1992).
74 Silva-Navas, J. *et al.* Flavonols Mediate Root Phototropism and Growth through Regulation of Proliferation-to-Differentiation Transition. *Plant Cell* **28**, 1372-1387, doi:10.1105/tpc.15.00857 (2016).
75 Wan, J. P. *et al.* UV-B Radiation Induces Root Bending Through the Flavonoid-Mediated Auxin Pathway in Arabidopsis. *Front Plant Sci* **9**, doi:ARTN 618 10.3389/fpls.2018.00618 (2018).
76 Coberly, L. C. & Rausher, M. D. Analysis of a chalcone synthase mutant in Ipomoea purpurea reveals a novel function for flavonoids: amelioration of heat stress. *Mol Ecol* **12**, 1113-1124, doi:DOI 10.1046/j.1365-294X.2003.01786.x (2003).
77 Hajrah, N. H. *et al.* Transcriptomic analysis of salt stress responsive genes in Rhazya stricta. *Plos One* **12**, doi:ARTN e0177589 10.1371/journal.pone.0177589 (2017).
78 Pi, E. X. *et al.* Quantitative Phosphoproteomic and Metabolomic Analyses Reveal GmMYB173 Optimizes Flavonoid Metabolism in Soybean under Salt Stress. *Mol Cell Proteomics* **17**, 1209-1224, doi:10.1074/mcp.RA117.000417 (2018).
79 Sarker, U. & Oba, S. Salinity stress enhances color parameters, bioactive leaf pigments, vitamins, polyphenols, flavonoids and antioxidant activity in selected Amaranthus leafy vegetables. *J Sci Food Agr* **99**, 2275-2284, doi:10.1002/jsfa.9423 (2019).
80 Li, W. Q. *et al.* The karrikin receptor KAI2 promotes drought resistance in Arabidopsis thaliana. *Plos Genet* **13**, doi:ARTN e1007076 10.1371/journal.pgen.1007076 (2017).
81 Nelson, D. C. *et al.* Karrikins enhance light responses during germination and seedling development in Arabidopsis thaliana. *Proc Natl Acad Sci U S A* **107**, 7095-7100, doi:10.1073/pnas.0911635107 (2010).
82 Bottomle.W, Smith, H. & Galston, A. W. A Phytochrome Mediated Effect of Light on Hydroxylation Pattern of Flavonoids in Pisum Sativum Var Alaska. *Nature* **207**, 1211-& (1965).
83 Jiao, Y. T., Xu, W. R., Duan, D., Wang, Y. J. & Nick, P. A stilbene synthase allele from a Chinese wild grapevine confers resistance to powdery mildew by recruiting salicylic acid signalling for efficient defence. *J Exp Bot* **67**, 5841-5856, doi:10.1093/jxb/erw351 (2016).
84 Gondor, O. K. *et al.* Salicylic Acid Induction of Flavonoid Biosynthesis Pathways in Wheat Varies by Treatment. *Front Plant Sci* **7**, doi:ARTN 1447 10.3389/fpls.2016.01447 (2016).
85 Dehghan, S. *et al.* Differential inductions of phenylalanine ammonia-lyase and chalcone synthase during wounding, salicylic acid treatment, and salinity stress in safflower, Carthamus tinctorius. *Bioscience Rep* **34**, 273-282, doi:ARTN e00114 10.1042/BSR20140026 (2014).
86 Kuhn, B. M. *et al.* 7-Rhamnosylated Flavonols Modulate Homeostasis of the Plant Hormone Auxin and Affect Plant Development. *J Biol Chem* **291**, 5385-5395, doi:10.1074/jbc.M115.701565 (2016).
87 Kurepa, J., Shull, T. E., Karunadasa, S. S. & Smalle, J. A. Modulation of auxin and cytokinin responses by early steps of the phenylpropanoid pathway. *Bmc Plant Biol* **18**, doi:ARTN 278

10.1186/s12870-018-1477-0 (2018).
88 Duellpfaff, N. & Wellmann, E. Involvement of Phytochrome and a Blue-Light Photoreceptor in Uv-B Induced Flavonoid Synthesis in Parsley (Petroselinum-Hortense Hoffm) Cell-Suspension Cultures. *Planta* **156**, 213-217, doi:Doi 10.1007/Bf00393727 (1982).
89 Taulavuori, K., Hyoky, V., Oksanen, J., Taulavuori, E. & Julkunen-Tiitto, R. Species-specific differences in synthesis of flavonoids and phenolic acids under increasing periods of enhanced blue light. *Environ Exp Bot* **121**, 145-150, doi:10.1016/j.envexpbot.2015.04.002 (2016).
90 Wang, S. *et al.* Genomes of early-diverging streptophyte algae shed light on plant terrestrialization. *Nat Plants*, doi:10.1038/s41477-019-0560-3 (2019).
91 Jiao, C. *et al.* The Genome of the Charophyte Alga Penium margaritaceum Bears Footprints of the Evolutionary Origins of Land Plants. *bioRXiv*, doi:10.1101/835561 (2019).
92 Shimizu, Y., Ogata, H. & Goto, S. Discriminating the reaction types of plant type III polyketide synthases. *Bioinformatics* **33**, 1937-1943, doi:10.1093/bioinformatics/btx112 (2017).
93 Shelest, E., Heimerl, N., Fichtner, M. & Sasso, S. Multimodular type I polyketide synthases in algae evolve by module duplications and displacement of AT domains in trans. *Bmc Genomics* **16**, doi:ARTN 1015
10.1186/s12864-015-2222-9 (2015).
94 Van de Peer, Y., Mizrachi, E. & Marchal, K. The evolutionary significance of polyploidy. *Nat Rev Genet* **18**, 411-424, doi:10.1038/nrg.2017.26 (2017).
95 Clark, J. W. & Donoghue, P. C. J. Whole-Genome Duplication and Plant Macroevolution. *Trends Plant Sci* **23**, 933-945, doi:10.1016/j.tplants.2018.07.006 (2018).
96 Thomas, B. C., Pedersen, B. & Freeling, M. Following tetraploidy in an Arabidopsis ancestor, genes were removed preferentially from one homeolog leaving clusters enriched in dose-sensitive genes. *Genome Res* **16**, 934-946, doi:10.1101/gr.4708406 (2006).
97 Schnable, J. C., Springer, N. M. & Freeling, M. Differentiation of the maize subgenomes by genome dominance and both ancient and ongoing gene loss. *P Natl Acad Sci USA* **108**, 4069-4074, doi:10.1073/pnas.1101368108 (2011).
98 Cheng, F. *et al.* Biased Gene Fractionation and Dominant Gene Expression among the Subgenomes of Brassica rapa. *Plos One* **7**, doi:ARTN e36442
10.1371/journal.pone.0036442 (2012).
99 Renny-Byfield, S., Gong, L., Gallagher, J. P. & Wendel, J. F. Persistence of Subgenomes in Paleopolyploid Cotton after 60 My of Evolution. *Mol Biol Evol* **32**, 1063-1071, doi:10.1093/molbev/msv001 (2015).
100 Moghe, G. D. *et al.* Consequences of Whole-Genome Triplication as Revealed by Comparative Genomic Analyses of the Wild Radish Raphanus raphanistrum and Three Other Brassicaceae Species. *Plant Cell* **26**, 1925-1937, doi:10.1105/tpc.114.124297 (2014).
101 Li, H. *et al.* Evolutionary and functional analysis of mulberry type III polyketide synthases. *Bmc Genomics* **17**, doi:ARTN 540
10.1186/s12864-016-2843-7 (2016).
102 Novak, P., Krofta, K. & Matousek, J. Chalcone synthase homologues from Humulus lupulus: some enzymatic properties and expression. *Biol Plantarum* **50**, 48-54, doi:10.1007/s10535-005-0073-y (2006).
103 Novak, P., Matousek, J. & Briza, J. Valerophenone synthase-like chalcone synthase homologues in Humulus lupulus. *Biol Plantarum* **46**, 375-381, doi:Doi 10.1023/A:1024326102694 (2003).
104 Abe, I. *et al.* A plant type III polyketide synthase that produces pentaketide chromone. *J Am Chem Soc* **127**, 1362-1363, doi:10.1021/ja0431206 (2005).

105 Abe, I., Oguro, S., Utsumi, Y., Sano, Y. & Noguchi, H. Engineered biosynthesis of plant polyketides: chain length control in an octaketide-producing plant type III polyketide synthase. *J Am Chem Soc* **127**, 12709-12716, doi:10.1021/ja053945v (2005).
106 Mizuuchi, Y. *et al.* Novel type III polyketide synthases from Aloe arborescens. *Febs J* **276**, 2391-2401, doi:10.1111/j.1742-4658.2009.06971.x (2009).
107 Mameda, R., Waki, T., Kawai, Y., Takahashi, S. & Nakayama, T. Involvement of chalcone reductase in the soybean isoflavone metabolon: identification of GmCHR5, which interacts with 2-hydroxyisoflavanone synthase. *Plant J* **96**, 56-74, doi:10.1111/tpj.14014 (2018).
108 Dakora, F. D. & Phillips, D. A. Diverse functions of isoflavonoids in legumes transcend anti-microbial definitions of phytoalexins. *Physiol Mol Plant P* **49**, 1-20, doi:DOI 10.1006/pmpp.1996.0035 (1996).
109 Graham, T. L., Graham, M. Y., Subramanian, S. & Yu, O. RNAi silencing of genes for elicitation or biosynthesis of 5-deoxyisoflavonoids suppresses race-specific resistance and hypersensitive cell death in Phytophthora sojae infected tissues. *Plant Physiol* **144**, 728-740, doi:10.1104/pp.107.097865 (2007).
110 Sepiol, C. J., Yu, J. J. & Dhaubhadel, S. Genome-Wide Identification of Chalcone Reductase Gene Family in Soybean: Insight into Root-Specific GmCHRs and Phytophthora sojae Resistance. *Front Plant Sci* **8**, doi:ARTN 2073 10.3389/fpls.2017.02073 (2017).
111 Subramanian, S., Stacey, G. & Yu, O. Endogenous isoflavones are essential for the establishment of symbiosis between soybean and Bradyrhizobium japonicum. *Plant J* **48**, 261-273, doi:10.1111/j.1365-313X.2006.02874.x (2006).
112 Wasson, A. P., Pellerone, F. I. & Mathesius, U. Silencing the flavonoid pathway in Medicago truncatula inhibits root nodule formation and prevents auxin transport regulation by rhizobia. *Plant Cell* **18**, 1617-1629, doi:10.1105/tpc.105.038232 (2006).
113 Yi, J., Derynck, M. R., Chen, L. & Dhaubhadel, S. Differential expression of CHS7 and CHS8 genes in soybean. *Planta* **231**, 741-753, doi:10.1007/s00425-009-1079-z (2010).
114 Chaudhary, B. *et al.* Reciprocal silencing, transcriptional bias and functional divergence of homeologs in polyploid cotton (gossypium). *Genetics* **182**, 503-517, doi:10.1534/genetics.109.102608 (2009).
115 Liu, S. L., Baute, G. J. & Adams, K. L. Organ and Cell Type-Specific Complementary Expression Patterns and Regulatory Neofunctionalization between Duplicated Genes in Arabidopsis thaliana. *Genome Biol Evol* **3**, 1419-1436, doi:10.1093/gbe/evr114 (2011).
116 Proost, S. & Mutwil, M. CoNekT: an open-source framework for comparative genomic and transcriptomic network analyses. *Nucleic Acids Res* **46**, W133-W140, doi:10.1093/nar/gky336 (2018).
117 Shoji, T. & Hashimoto, T. Recruitment of a duplicated primary metabolism gene into the nicotine biosynthesis regulon in tobacco. *Plant J* **67**, 949-959, doi:10.1111/j.1365-313X.2011.04647.x (2011).
118 Moghe, G. D. & Last, R. L. Something Old, Something New: Conserved Enzymes and the Evolution of Novelty in Plant Specialized Metabolism. *Plant Physiol* **169**, 1512-1523, doi:10.1104/pp.15.00994 (2015).
